# Predictable Engineering of Signal-Dependent Cis-Regulatory Elements

**DOI:** 10.1101/2025.03.07.642002

**Authors:** Jake Cornwall-Scoones, Dirk Benzinger, Tianji Yu, Alberto Pezzotta, Andreas Sagner, Lina Gerontogianni, Shaun Bernadet, Elizabeth Finnie, Giulia L. M. Boezio, Hannah T. Stuart, Manuela Melchionda, Oliver C. K. Inge, Bianca Dumitrascu, James Briscoe, M. Joaquina Delás

## Abstract

Cis-regulatory elements (CREs) control how genes respond to external signals, but the principles governing their structure and function remain poorly understood. While differential transcription factor binding is known to regulate gene expression, how CREs integrate the amount and combination of inputs to secure precise spatiotemporal profiles of gene expression remains unclear. Here, we developed a high-throughput combinatorial screening strategy, that we term NeMECiS, to investigate signal-dependent synthetic CREs (synCREs) in differentiating mammalian stem cells. By concatenating fragments of functional CREs from genes that respond to Sonic Hedgehog in the developing vertebrate neural tube, we found that CRE activity follows hierarchical design rules. While individual 200-base-pair fragments showed minimal activity, their combinations generated thousands of functional signal-responsive synCREs, many exceeding the activity of natural sequences. Statistical modelling revealed CRE function can be decomposed into specific quantitative contributions in which sequence fragments combine through a multiplicative rule, tuned by their relative positioning and spacing. These findings provide a predictive framework for CRE redesign, which we used to engineer synthetic CREs that alter the pattern of motor neuron differentiation in neural tissue. These findings establish quantitative principles for engineering synthetic regulatory elements with programmable signal responses to rewire genetic circuits and control stem cell differentiation, providing a basis for understanding developmental gene regulation and designing therapeutic gene expression systems.

## Introduction

During embryonic development and tissue homeostasis, stem cells continuously undergo differentiation to generate a diverse array of specialised cell types. Such cell fate decisions are driven by external signals that regulate cell type specific gene expression programmes (*1*, *2*). This process relies on cis-regulatory elements (CREs) – non-coding regions of the genome that act as molecular control switches for gene expression. These DNA sequences function as information processing devices by recruiting signal-dependent and cell type specific transcription factors (TFs) to control the transcription of target genes.

Despite their critical role, our understanding of how CREs integrate inputs to drive cell-type specific gene expression remains incomplete and lacks unifying principles. Uncovering quantitative rules for the structure and sequence-composition of CREs would have far-reaching implications (*3*), advancing our understanding of how stem cells interpret signalling cues to establish and maintain complex tissues, and enabling the engineering of synthetic cis-regulatory elements (synCREs) to direct cell fates away from pathological states and toward regenerative outcomes.

Short DNA motifs within CREs bind specific TFs (*4*) and may be activating, repressing, or both depending on the context and signalling environment (*5–7*). While functional genomics has demonstrated that CRE activity depends on the presence of these motifs (*8*, *9*), we lack a fundamental understanding of how the structure and composition of CREs facilitate signal-dependent gene expression.

Various models for the design rules of CREs have been proposed (*10*). These fall along a spectrum (*11*) that place different emphasis on the relative positioning of motifs. At one extreme, the billboard model suggests that CREs are simply “bags of motifs” (*12*), where the activity contributions of motifs are independent of each other. At the other extreme, the enhanceosome model, exemplified by the IFN-β locus, argues that CRE activity depends on a specific physical conformation (*13*), where the “grammar” – the precise positioning and spacing of motifs – determines transcription (*14*, *15*). Indeed, there are clear examples of local rigid grammar (*16–18*), however for other CREs, grammar appears subtle and case-dependent (*19*).

Recent progress in applying deep-learning to construct novel synthetic CREs (*20–23*) points to a hidden structure in the configuration of CREs, which can be learned from assaying the regulatory activity of large sets of DNA sequences. These deep learning models are designed to handle potential complexities and higher-order interactions between TFs and CRE in a way that simpler mechanistic models cannot (*24*). It is possible that CRE structure is inherently complex, necessitating complex high dimensional models. Alternatively, the apparent complexity of deep-learning models may conceal a simpler set of rules or principles that provide an interpretable explanation of CRE function.

Synthetic genomics offers a path to understanding CRE activity (*25*). By going beyond natural variation, synthetic genomics allows for the systematic exploration of ‘design space’ (*26*, *27*), providing an understanding of how CRE activity depends on its sequence building-blocks. To realise this potential, however, novel high-throughput approaches for building and testing large cohorts of synthetic regulatory sequences are required (*24*).

The ability to rationally design synthetic regulatory sequences with precisely tuned activities has profound implications for therapeutic applications and tissue engineering. Understanding and exploiting the rules of CRE construction could allow the design of “super-physiological- enhancers” that drive gene expression more efficiently than their natural counterparts, or engineer tissue-specific regulatory elements that enable targeted gene therapy with minimal off-target effects (*28*, *29*). Such synthetic enhancers could be particularly valuable for fine-tuning the expression of therapeutic transgenes, reprogramming cell fates in regenerative medicine, or creating more effective gene therapy vectors that achieve therapeutic levels of expression in specific target tissues while remaining silent in others.

Here, we developed NeMECiS (Nested Modular Expressible Cis Screening), a novel combinatorial screening strategy to build and assay synCREs. Using vertebrate neural progenitor diversification as a model, in which different concentrations of the signal Sonic Hedgehog (Shh) instruct alternative neural cell types with distinct gene expression programmes (*30*), we generated exhaustive three-way fusions of fragments from a set of known CREs. The comprehensive, combinatorial coverage of synCRE design-space afforded by NeMECiS enabled inference of the rules of CRE activity. Surprisingly, individual 200-base-pair fragments of CREs showed minimal activity, yet in combination generated thousands of functional signal-decoding synCREs. Statistical modelling revealed the hierarchical rules of CRE activity, where specific sequence fragments contribute to signal-dependent activity, but are tuned by spacing-dependent pairwise interactions.

The learned rules extrapolate beyond the training design-space, enabling the principled re- design of CREs controlling stem cell diversification. As a proof-of-concept, we modified a hedgehog-responsive element controlling motor neuron differentiation (*31–33*), generating synthetic variants with enhanced or reduced activity compared to the endogenous sequence. These engineered CREs successfully directed alternative spatial patterning outcomes in neural tissue. This synthetic regulatory toolkit provides a versatile platform for programming cell fate decisions, offering opportunities to investigate pathological consequences of aberrant signal interpretation and to optimise therapeutic gene expression in target tissues.

## Results

### Complex Architectural Requirements for Hedgehog Dependent CRE Activity

To analyse CRE activity we took advantage of Shh signalling dependent gene expression in developing neural progenitors using an mouse embryonic stem cell (mESC) directed differentiation model of neural tube patterning (*34*, *35*) (Fig. S1A). Here, neural progenitors supplied with different concentrations of SAG, a Shh signalling pathway agonist, are directed to distinct identities that correspond to the *in vivo* pattern of progenitor domains (*36*). To assay CRE activity, we integrated a lentivirus containing regulatory sequences controlling the expression of a fluorescent reporter (ZsGreen)(*37*, *38*) and a constitutively expressed marker, mScarlet3, to identify transduced cells (Fig. 1A). Signal-dependent CRE activity was assayed by exposing cells to different SAG concentrations and analysing by flow cytometry at day 6 of the differentiation, when Shh-responsive ventral neural progenitors are usually present (*36*). When performed using a well-characterised CRE for the Shh-responsive TF Olig2, we observed an increase in ZsGreen levels as a function of SAG concentration (Fig. 1B), consistent with *in vivo* reporters (*31*, *32*).

**Figure 1.**
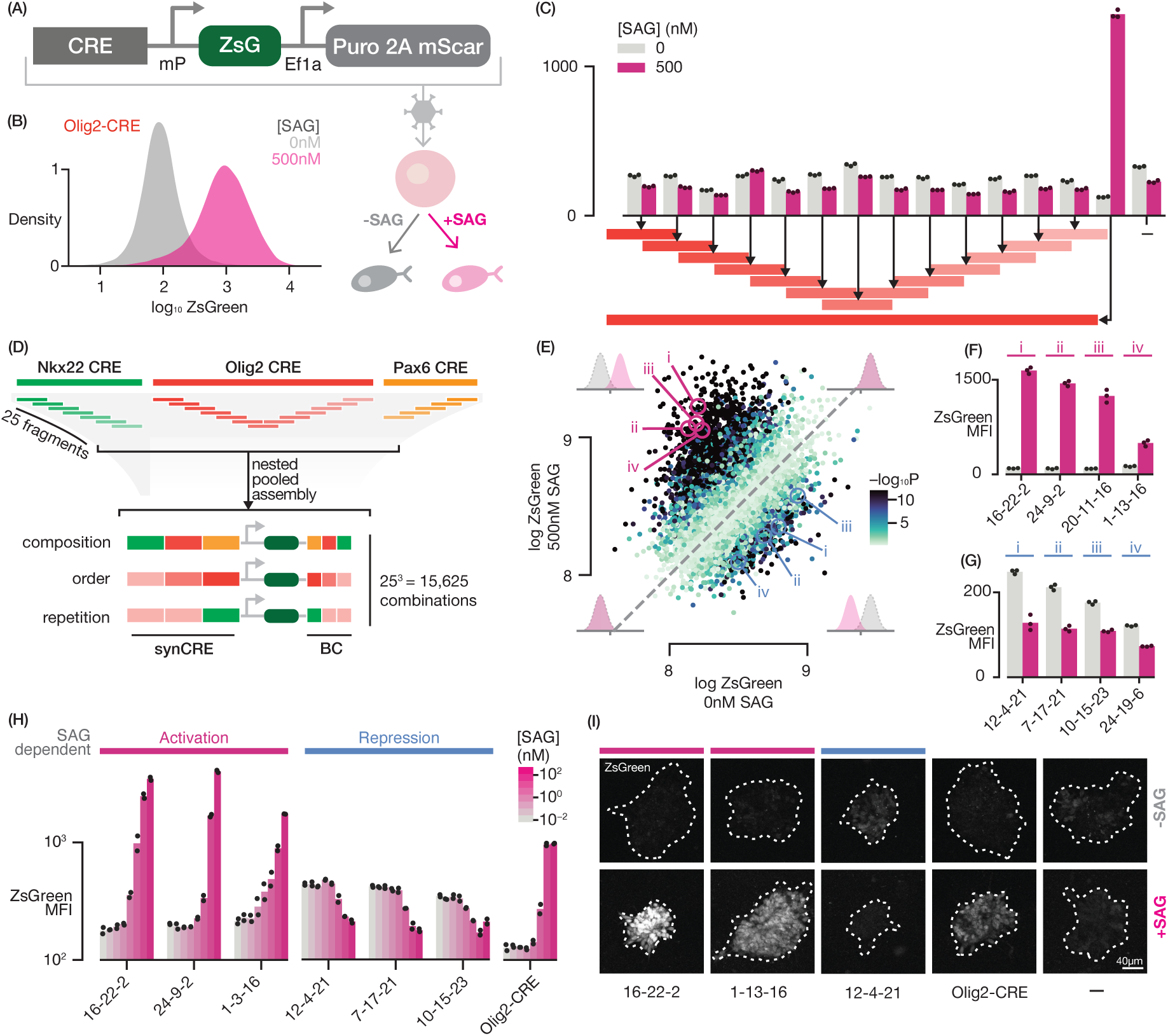
NeMECiS: A Combinatorial Screening Strategy Reveals Flexible Composability of Regulatory Elements. (A) Schematic of lentiviral reporter construct, consisting of a CRE upstream of the Shh minimal promoter driving the green fluorescent protein ZsGreen, also containing the selectable markers mScarlet3 and Puromycin resistance. Lentivirus is generated from the cloned constructs, used to infect differentiating mESCs supplied with variable SAG concentrations, allowing for assaying of Hedgehog-dependent CRE activity. (B) Live flow-cytometry of ZsGreen levels (kernel density estimate) in mScarlet3+ day-6 neural progenitor cells transduced with an Olig2-CRE reporter construct and treated with 0 or 500nM SAG. (C) Using an equivalent live-flow cytometry quantification procedure, ZsGreen mean-fluorescence intensities (MFI) for day-6 neural progenitor cells cultured in 0 or 500nM SAG transduced with either the entire Olig2-CRE, or 200bp fragments of the Olig2-CRE (100bp offset), or promoter-only constructs without any CRE (“–“). Bars represent mean across biological replicates, and points represent individual replicate datapoints (N=3). (D) Schematic of nested pooled sequential cloning strategy of NeMECiS, sequentially inserting one of 25 CRE fragments (with associated barcodes, BC) into a lentivirus backbone, yielding a pooled library of 15,625 combinations of CRE fragment triplet-fusions (synCREs), with barcode concatemers that reflect (in reverse) the composition and order of fragments by synCRE. (E) Inferred log ZsGreen levels for the NeMECiS library in 0nM and 500nM SAG, coloured by the significance level of a non-zero difference between SAG concentrations (–log_10_P, z-test; N=13,991). Dashed-line represents equal inferred fluorescence in 0nM and 500nM SAG. Inset are diagrams of the expected fluorescence distributions for 0nM (grey) and 500nM (magenta) SAG in the four quadrants of the graph. Validation constructs in Fig. 1F-G are encircled and annotated. (F) Independently constructed and tested validation constructs, assayed as in Fig. 1B-C, for synCREs predicted to show activation. (G) As in Fig. 1F, but for repression. (H) Serial dilution of SAG concentration (8 concentrations at 5-fold dilution starting at 500nM), assayed as above. (I) Representative images of day 6 neural progenitor colonies displaying ZsGreen levels for validation synCREs and controls, under 0nM and 500nM SAG. Magenta label represents “activation” synCREs and blue label represents “repression” synCREs. Colonies outlined (white dashed line) using a Sox2 immunofluorescence staining. Scale bar 40 μm.

Previous high-throughput screens of CRE activity have been limited to DNA elements <300bp in size, typically centred on ChIP-/ATAC-seq peaks (*39–43*). We therefore generated a nested set of reporters comprising 200bp fragments of the Olig2 CRE. Surprisingly, none of the fragments alone showed substantial SAG-dependent fold-changes in activity (Fig. 1C). This could indicate a rigid architecture of the CRE, consistent with the enhanceosome model, in which the CRE activity is dependent on its integrity. Alternatively, unknown rules of composability may describe how modules come together to generate CRE activity that is greater than the sum of its parts.

### Combinatorial assay of synCRE activity demonstrates flexible composability of regulatory information

To discriminate between alternative hypotheses, we tested whether combining CRE sequence fragments was sufficient to recover SAG-dependent activity. To test if activity can be recovered, and to infer the hidden rules of CRE composability in an unbiased manner, we designed an exhaustive and systematic approach. We term this approach NeMECiS (Nested Modular Expressible Cis Screening).

To this end we established a sequential, combinatorial pooled cloning strategy to fuse a library of CRE fragments in exhaustive permutations to establish synCREs (Fig. 1D). In this approach, 608bp synCREs, comprising three 200bp fragments connected by 4bp spacers, are assembled spanning the full permutation-set of CRE fragments (Fig. S1B). During the assembly a concatemer of 8bp barcodes is established, allowing for direct readout of synCRE composition by high-throughput sequencing (Materials and Methods) (Fig. S1D). Using three established CREs controlling neural progenitor genes Nkx2-2, Olig2 and Pax6 (*31*, *44*) (Fig. S1E), we generated a library comprising 25^3^=15,625 synCREs, the exhaustive permutation- set of twenty-five 200bp fragments from these CREs (*32*) (Fig. S2A-D).

To measure synCRE activity across this library, we used lentiviral transduction to integrate synCREs into mESCs (*41*) (MOI<1) and differentiated them to neural progenitors, treating cells with either 0nM or 500nM SAG (Fig. S1C). To quantify SAG-dependent synCRE activity, we flow-sorted neural progenitors using the ZsGreen reporter into four equally sized bins, representing different levels of activity, and sequenced the barcodes in each bin (Fig. S2E-G). From the counts of barcodes in each activity bin we used Bayesian inference to infer the mean ZsGreen level (Materials and Methods). The measurements showed good reproducibility across replicates (PCC: 0nM SAG = 0.45; 500nM SAG = 0.60; Fig. S3A) and established a set of 13,991 synCREs (90% of theoretical max) that pass quality-control.

The assay revealed a broad spectrum of signal-dependent behaviours (Fig. 1E), ranging from activation in SAG conditions (23% of synCREs) to repression (7% of synCREs) and insensitivity (Fig. S3B-E). We validated with flow cytometry and microscopy a selection of synCREs that were either induced or repressed by SAG (Fig. 1F-I & S3F-I). This showed tight SAG dose-response relationships (Fig. 1H). Strikingly, in response to 500nM SAG, some activating synCREs displayed activities that were 187% greater than the strong Olig2 CRE. Across the whole dataset, the majority of inducing (84%) and repressing (86%) synCREs comprised combinations of fragments from the CREs of the different genes.

We therefore conclude that despite the weak or absent activity of individual 200bp CRE fragments, a surprising proportion of fragment permutations showed a range of different activities. This argues that while CRE activity is greater than the sum of its parts, the rules of composability are not as rigid as the enhanceosome model but could be consistent with a modular mechanism in which individual fragments make a specific contribution to the overall activity of the synCRE.

### CRE activity can be decomposed into modular contributions

The possibility that CRE activity can be decomposed into a series of separable modules could explain why signal-dependent synCREs can be readily assembled by concatenating CRE fragments. To test this, we established a linear modelling framework (Materials and Methods). In this framework each fragment makes a discrete, predictable contribution to overall transcriptional output both with and without Hedgehog signalling, independently of the other fragments in the synCRE (Fig. 2A & S4A-C). This revealed clear statistical signatures of signal-dependent fragment activities, which we grouped into three categories based on their response to Hedgehog signalling: inducing, repressing and neutral (Fig. S4D). To validate these predictions, we compared the mean ZsGreen expression levels between synCREs containing a specific fragment versus those without it, measuring their responses to SAG (Fig. 2B & S4D). The observed changes closely matched the predictions from our linear modelling, indicating that a major component of the overall activity of each synCRE can be explained by individual fragment contributions Such modularity in the composition of synCREs is consistent with the extensive literature on TF-motif binding driving signal-dependent activities (*44–46*). To understand the sequence features underlying Hedgehog-responsive modules, we performed motif enrichment analysis on synCREs’ signal-dependent activity. This identified a cluster of motifs that included the DNA binding motif of Gli transcription factors (the transcriptional effectors of the Hedgehog pathway (*47*)) as the strongest positive correlate of logFC SAG (Fig. 2C, S5A-C). Indeed, when we mutated the Gli motif in a functional synCRE (Fig. S5D), we found that the Hedgehog signalling-dependent fold change of activity decreased substantially (Fig. 2D), both raising the activity in the absence of SAG and lowering the activity in its presence (*48*). This is consistent with the Gli proteins acting as transcriptional repressors in the absence of Hedgehog signalling and activators in its presence (*47*). Further, across the entire NeMECiS dataset, synCREs that contain Gli binding sites have elevated responses to SAG (Fig. 2E). However, the overlap in the distribution of SAG responsiveness of fragments with or without a binding motif for Gli proteins (Fig. 2E) suggested that motif composition alone is insufficient to explain the extent of synCRE signal-dependent activity: 25% of strongly inducing fragments lacked detectable Gli motifs, while 14% of non-responsive fragments contained them. Additional sequence motifs enriched in Hedgehog-responsive fragments suggest complex combinatorial control beyond Gli binding (Fig. S5A-C). These findings indicate that while Gli motifs contribute substantially to Hedgehog responsiveness, additional sequence features shape the signal-dependent activity of synCREs.

**Figure 2.**
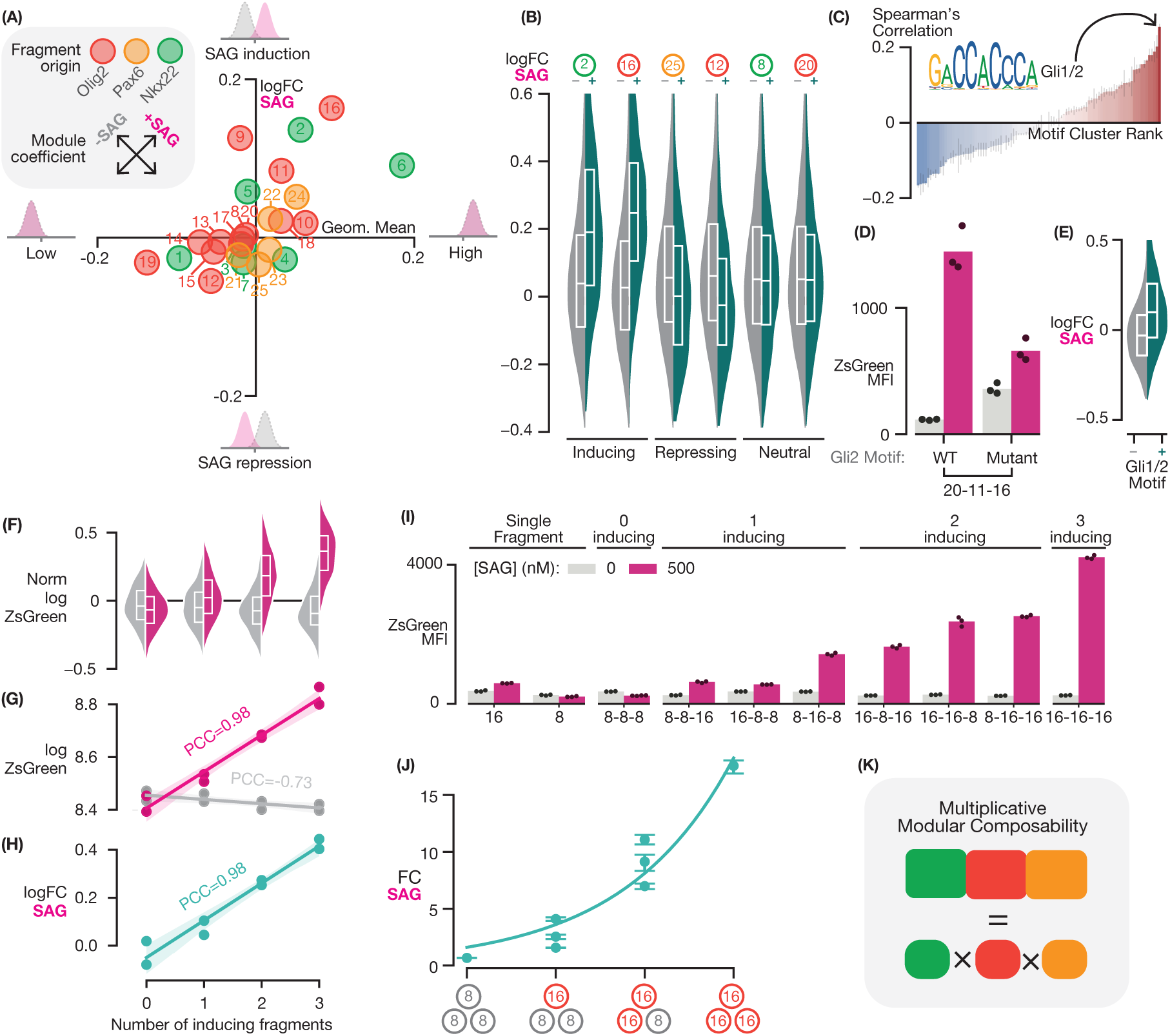
**Modular Contributions of CRE Fragments to Activity Follow a Multiplicative Rule.** (A) Fragment contributions (module coefficient) from linear modelling of log ZsGreen (±SAG) for each fragment in a synCRE. Each circle represents the contribution of a single fragment, coloured by their originating endogenous CRE. Each fragment has a weight for 0nM and 500nM SAG, which are used to compute logFC (logFC SAG (log fold change of ZsGreen = w_500_ – w_0_) and their geom. (geometric) mean ([w_0_ + w_500_]/2). The diagonals of the graph represent the raw fragment weights (see key). (B) Violin plots with associated quartiles for the logFC SAG of all synCREs without (grey) or with (teal) a given fragment, shown for two examples of each of the three classes of fragment activity. Fragments are denoted above the plot, coloured by their originating endogenous CREs. (C) Spearman’s correlation across synCREs of the presence of each DNA motif cluster with the corresponding logFC SAG. Performed separately for each replicate: bars represent mean across replicates; error bars indicate maximum/minimum. Annotated is the top hit motif cluster containing Gli1/2 and Zbtb7b/c, displaying the logo plot of the position weight matrix for Gli2 (JASPAR2024: MA0734.4 (*73*)). (D) ZsGreen mean fluorescence intensity (MFI) in 0nM (grey) and 500nM (magenta) SAG for a wildtype (WT) strong inducing synCRE (f20-f11-f16) and a version where the (only) Gli2 motif was mutated (see Fig. S5D). Bars represent means across three biological replicates, and points represent individual replicates. (E) Violin plots with associated quartiles for the logFC SAG of all synCREs without (grey) or with (teal) a motif found in the cluster containing Gli1/2 and Zbtb7b/c. (F) Violin-plot and associated quartiles of log ZsGreen for 0nM (grey) or 500nM (magenta) SAG for the set of synCREs that lack repressing fragments, stratified by the number of inducing fragments. Values are normalised by subtracting the population mean across SAG concentrations (i.e. the intercept of the linear model). (G) The mean logZsGreen for the stratification detailed in Fig. 2F (PCC = pearson’s correlation coefficient). Line represents replicate average, points represent individual replicates, shaded region represents 95% CI of a linear fit. (H) logFC SAG with the same stratification and depiction as Fig. 2F-G. (I) Individually built and assayed synCRE reporter constructs for the exhaustive set of ordered triplets that contain fragment 8 and 16, with corresponding single fragment reporter constructs. Data is plotted as in Fig. 2D. Number of inducing (i.e. fragment 16) fragments is annotated. (J) Fold change of mean fluorescence intensity of ZsGreen with respect to SAG (FC SAG) is plotted for the data in Fig. 2I. Each datapoint corresponds to a single synCRE, and the error bars represent the maximum/minimum FC SAG across all replicate-by-replicate comparisons. An exponential curve is fit to the data. (K) Schematic for the multiplicative modular composability model, where the activity of a synCRE is the product of the activities of each of its components.

### Signal-dependent CRE activity follows a multiplicative composability rule

We next sought to identify the modular composability rules. In principle this could follow several forms: Boolean, where activity depends on the presence of a fragment; or cumulative, where activity depends on the numbers of each fragment present in a synCRE. We grouped synCREs based on the number of inducing fragments, disregarding synCREs containing repressing fragments. This revealed the activity to be cumulative, with a substantial, monotonic increase in activity in response to 500nM SAG (Fig. 2F).

The cumulative behaviour could in principle be additive, multiplicative, or adopt an alternative non-linearity (*49*). Whereas for an additive model each fragment would add a fixed amount to the total activity, with a multiplicative model each additional fragment multiplies the existing activity by a factor, leading to exponential-like increases in overall activity. Consistent with a multiplicative model, we found a good fit of the mean log activities, and logFC, as a function of the number of activating fragments (Fig. 2G-H). A comparison of additive versus multiplicative models across all the dataset revealed the multiplicative model captured the behaviour better across fragment compositions (Fig. S6A-C). These results agree with the predictions of the well-established thermodynamic description of CRE activity (Materials and Methods)(*50–55*).

We tested the multiplicative model with an arrayed, individualised experiment, where we generated exhaustive permutations of the neutral fragment 8 (f8) and the activating fragment 16 (f16). Signal-dependent activity increases as a function of the number of activating fragments, minimal in f8-f8-f8 and maximal in f16-f16-f16 (Fig. 2I). In between these extremes, activities fit well to a multiplicative scaling rule (Fig. 2J). Repeating this approach with the activating (f2) and neutral (f20) fragments combined with either f8, f16, yielded equivalent results (Fig. S6E-G). A corollary of this rule is that a linear increase in inducing fragment frequency has an exponential effect on activity, helping explain why individual fragments show low activities yet readily combine to make functional synCREs (Fig. 2K).

### Synergies among modules shape activities

The multiplicative model assumes that transcription factors bind to fragments independently. Yet, this is likely an oversimplification, as transcription factors can bind cooperatively to DNA (*49*, *56–60*). We therefore tested whether a model that accounted for interactions between fragments, and not just their modular action, fit the data better. NeMECiS is uniquely positioned to statistically isolate these higher-order effects (Fig. 3A), as it assays every fragment in every combination and ordering.

**Figure 3.**
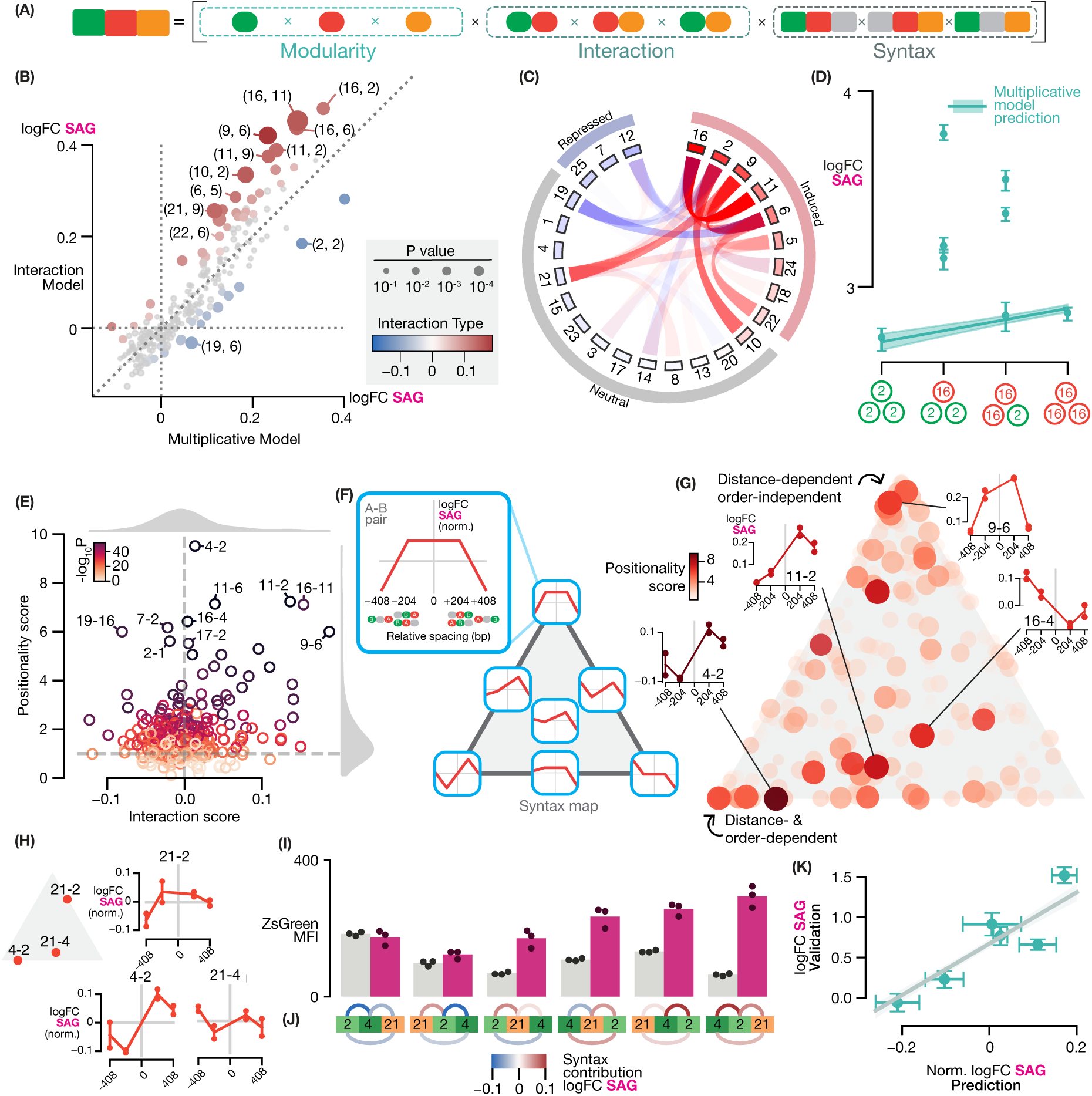
**Pairwise Interactions and Syntax Rules Shape Regulatory Element Activity.** (A) Schematic of the extended linear model incorporating modularity, interaction and syntax layers. (B) Average contribution of each unique pair of fragments to the log fold change with respect to SAG (logFC SAG) in the multiplicative model and the interaction model. Dot size corresponds to the significance level of a z-test for non-zero interaction coefficients, and dot colour corresponds to the deviation of interaction coefficients from zero. Points above the dashed-line diagonal (red) indicate positive interactions (i.e. logFC SAG for fA&fB > fA × fB) and points below (blue) indicate negative interactions (i.e. logFC SAG for fA&fB < fA × fB). (C) Circos plot for the distribution of positive (red) and negative (blue) significant (p<0.01) interactions between fragments. (D) Log fold change of ZsGreen mean fluorescence intensity with respect to SAG (logFC SAG) for individually built and assayed combinations of two activating fragments (f2 and f16), grouped by the number of each fragment. Datapoints represent mean across replicates for each synCRE, and error-bars represent the minimum and maximum value across exhaustive replicate comparison. Overlaid are the predicted activities from the multiplicative model for each of the combinations, based on fitting to data considering combinations of f2 and f16 individually with a neutral fragment f8, where the error-bar is the 95% confidence interval. (E) Distribution of interaction coefficient compared to positionality score (see Methods) for all heterologous fragment pairs. Top and right are kernel-density estimates of the marginal distributions. Points are coloured by significance of spacing dependent variance (Kruskal-Wallis H test for average logFC SAG across each possible spacing). (F) Key for syntax map, built by orthonormal basis decomposition. Each position on the triangular manifold represents the relative contribution of each of three orthonormal bases to the total spacing-dependent variance of logFC SAG. Overlaid are diagrams of the logFC SAG (normalised to its mean) across all possible relative fragment spacings (see pull-out for an example of an ordered pair of fragments A & B). (G) Orthonormal basis decomposition for all heterologous pairs of fragments, coloured by positionality score, with the shade and diameter of the points corresponding to the significance level in Fig. 3E. Inset are examples of spacing-dependent activity for different ordered fragment pairs. (H) Case study of spacing-dependent activity for all pairings of a trimer (f2, f4, f21), where dots on the triangle correspond to the orthonormal basis decomposition for each pairing. (I) Individually built and assayed exhaustively re-ordered synCREs comprising f2, f4, and f21 (MFI = mean fluorescence intensity; bars represent means across biological replicates; points represent individual replicates). (J) Corresponding syntax contributions for each spaced-pair by synCRE (corresponding to the values in Fig. 3H). (K) Comparison of predicted logFC SAG from linear modelling of NeMECiS to independently validated constructs for each of the six re-orderings of f2-f4-f21. Points represent means across replicates, and error bars represent maximum/minimum across replicates. Predicted logFC SAG is normalised by subtracting ‘modular’ and ‘interactional’ contributions.

Given the recent success of deep-learning in capturing regulatory sequence functionality (*20*, *23*, *61*, *62*) and in building synthetic CREs (*21*, *22*, *43*, *63*), we first trained several deep-learning models on our data (*20*, *23*, *64*, *65*) (Fig. S7A-B). These models generally perform better than our linear modular model (PCCs: 0.61-0.66 on held-out test-data compared to 0.56 of the linear model; Fig. S7B). This suggested that the linear modular multiplicative model, while a good first approximation, does not account for the full complexity of regulatory activity across our synCRE design space.

Because the deep learning models provided superior predictive power for CRE activity, but their complexity limited mechanistic interpretation, we developed an interpretable framework. We decomposed fragment interactions into two components. First, we quantified how pairs of fragments jointly influence CRE activity beyond their individual effects, which we term ’interaction’. Second, we measured how the spacing between fragments affects CRE activity independent of these pairwise interactions, termed ’syntax’ (Fig. 3A). This outperforms the simple multiplicative model across experimental replicates, revealing significant and reproducible effects for both interaction and syntax terms (Fig. S8).

Strikingly, the fit quality of our synergistic model matched those of the deep-learning models (Fig. S7B). There was a clear correspondence in predictions for synCRE activity between the synergistic model and the deep-learning models, which suggests that both classes of model are learning similar rules (Fig. S7A). This raises the possibility that sequence-to-activity deep-learning models are performing a two-step process, first identifying local regulatory modules, then applying a multiplicative rule modulated by pairwise interactions to predict the activity of a whole CRE from its collection of modules.

We compared the simple multiplicative model to the model incorporating pairwise interactions between fragments (Fig. 3B & S9A-B). Most fragment pairs were well described by the multiplicative behaviour, as shown by their distribution along the diagonal. However, there were significant instances of deviations above (positive interaction; 15% at P < 0.05) and below (negative interaction; 13% at P < 0.05) the diagonal. Analysis of all pairwise fragment combinations revealed that positive interactions occurred more frequently between pairs of inducing fragments (1.6-fold enrichment, permutation test), whereas negative interactions were more common between inducing-repressing pairs (1.3-fold enrichment; Fig. 3C, Fig. S9A-B).

To test the role of interactions in modular composition, we examined combinations of two activating fragments with different strengths: f2 (weaker) and f16 (stronger). The multiplicative model predicted that sequentially replacing f2 with f16 in the f2-f2-f2 synCRE would produce a monotonic increase in SAG-dependent activity. However, our experimental results showed that synCREs with combinations of f2 and f16 fragments produced higher activities than predicted, exceeding the f16-f16-f16 arrangement (Fig. 3D). This finding aligns with our global analysis showing interactions between inducing fragments, and with the strong positive interaction coefficient between f2-f16 predicted in our synergistic model (Fig. 3C). By combining both modular and synergistic effects, synCREs achieved activities up to 276% higher than the Olig2 CRE. These results confirm that while CREs can be broken down into modules, pairwise interactions can significantly influence their activity.

### Pairwise syntax rules explain CRE grammar

We next asked whether synCRE activity is dependent on the spacing and ordering of its fragments. By analogy to linguistics, we term such contributions syntax. Our exhaustive, combinatorial synCRE library contains every pair of fragments in every combination of spacing (adjacent vs separated by a fragment) and ordering (‘AB’ vs ‘BA’). These four classes act as components in the ‘syntax’ layer of our synergistic model (Fig. 3A). To identify contributions of syntax to signal-dependent CRE activity, we devised a positionality score (Materials and Methods), quantifying the relative variance of activity due to the spacing of a pair of fragments (positionality score > 1 implies spacing-dependent grammar). This revealed that, while most fragment pairs show ‘syntax-free’ interaction (positionality score ≈ 1), 86 fragment pairs showed significant spacing-dependent interactions (26.5%; positionality score > 2) (Fig. 3E).

Having identified a subset of fragment pairs displaying syntax effects, we sought to classify the different types of spacing-dependent activity. We transformed the measurements of spatial interactions into combinations of fundamental patterns (orthonormal bases) (Materials and Methods) (Fig. 3F). This allowed us to project the data into a 2D space, revealing examples of distance-dependent order-independent (e.g. f9-f6), and distance- and order-dependent (e.g. f4-f2), syntax (Fig. 3G-H). Similar results were found by k-means clustering of spacing-dependent synergies (Fig. S9D).

A consequence of these syntax rules is that the ordering of fragments in a synCRE should influence its activity. Specifically, permuting the order of fragments comprising a synCRE should maximise or minimise combined syntax-effects across the pairs of these fragment trimers. By building exhaustive six-way permutations of a series of fragment trimers (Fig. 3I, Fig. S10A), we found repeated examples of order-dependence, with the levels of ZsGreen induced by 500nM varying between 2.3 and 4.6-fold (6 triplet sets x 6 permutations). In some instances, these ordering effects are sufficiently influential that some orderings show strongly SAG-dependent activities, whereas other orderings are insensitive to SAG, despite being comprised of the same modules.

This order-dependence is explained by our measured pairwise syntax rules. For example, the ordering in f4-f2-f21 maximises spacing-dependent synergy for each comprising pair (Fig. 3H). Signal-dependent activity is maximised when: 1) f4 is immediately before f2; 2) when f2 and f21 are adjacent in either ordering; and (3) when f4 is 408bp upstream of f21 (Fig. 3I-J). Indeed, the learned spacing-dependence predicts well the relative signal-dependent activities when validated individually (Fig. 3K & S10B-C). This argues that the ordering-dependent differences in synCRE activity can be explained by the linear combination of comprising pairwise spacing rules, with deviations from this trend (e.g. f10-f11-f9, f11-f10-f9; Fig. S10) potentially indicating three-way interactions that defy pairwise reduction.

### Principled re-engineering of motor neuron progenitor patterning

The widespread examples of spacing-dependent synergies prompted us to ask whether endogenous Shh-responsive CREs have a fragment ordering that maximises expression. Analysis of 600bp windows from the Nkx2-2 and Olig2 CREs, divided into 200bp fragments and permuted exhaustively, showed that non-natural fragment orderings could achieve higher signal-dependent activity than natural orderings (e.g. f4-f2-f3 > f2-f3-f4; f9-f11-f10 > f9-f10-f11; Fig. S10). We then extended our prediction framework to evaluate signal-dependent activity across arbitrary-length fragment combinations (Materials and Methods; Fig. 4A). Applying this to the 1400bp Olig2 CRE sequence, we analysed systematically all possible arrangements of its seven 200bp fragments (5,040 permutations) to predict signal-dependent activity across the complete full-length synCRE (FL-synCRE) design-space (Fig. 4B-C & S11A). This analysis revealed that while the natural Olig2 CRE fragment ordering performs above average (73rd percentile; Fig. 4C & S11A), it does not achieve maximal signal-dependent activity.

**Figure 4.**
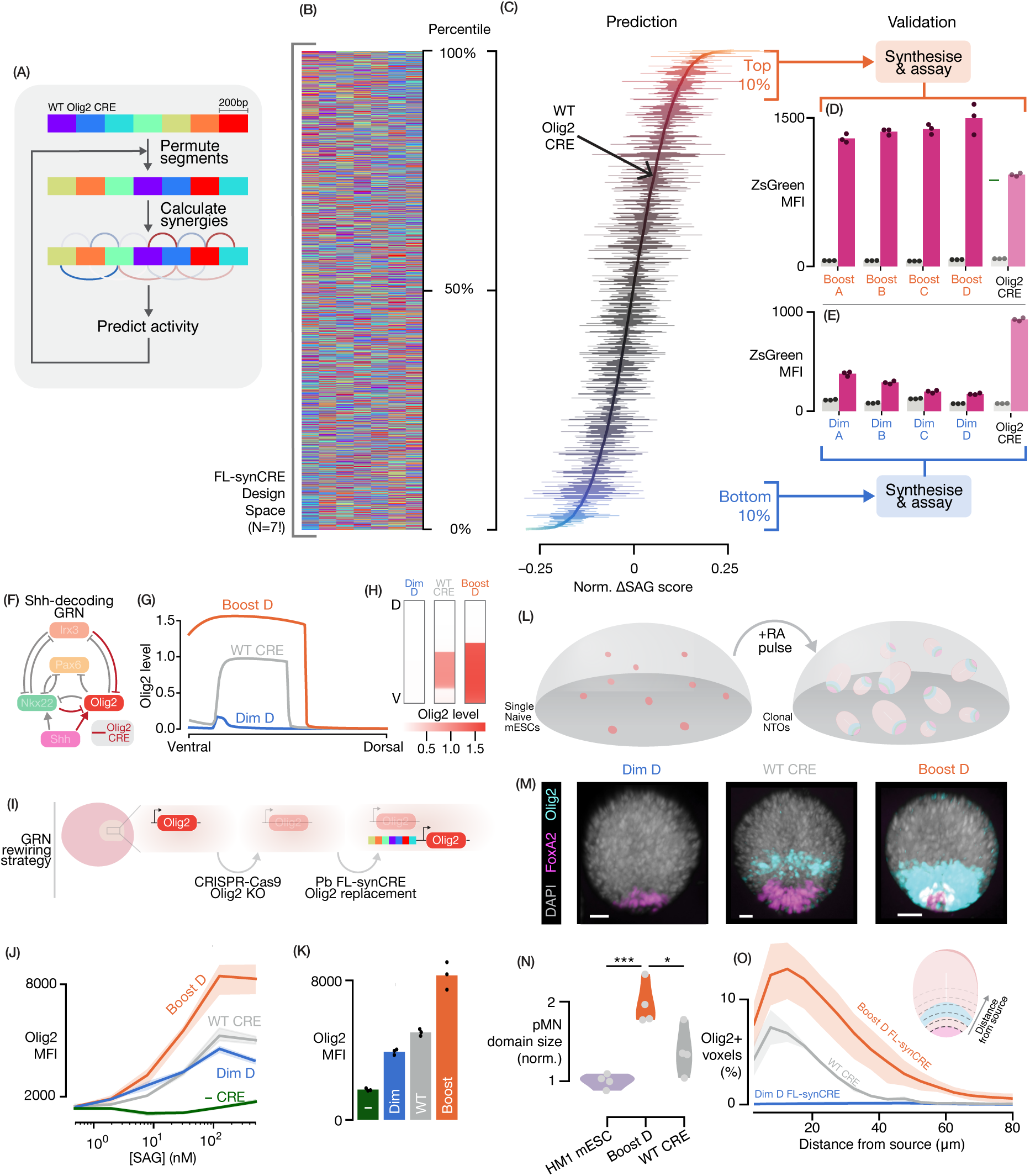
**Predictable Engineering of CREs to Control Neural Patterning.** (A) Schematic representing iterative shuffling of 200bp fragments of the WT Olig2 CRE to generate FL-synCREs. Curved lines connecting fragments represent syntax contributions (see Fig. S11A). (B) Full FL-synCRE design space of permutations (5040 combinations), coloured as in Fig. 4A, ordered by predicted (normalised) log-fold change of ZsGreen level in response to 500nM SAG (logFC SAG). (C) Corresponding logFC SAG for each FL-synCRE, normalised to consider only positional- dependent effects, ordered by the average predicted logFC SAG across replicates. Horizontal bars represent predictions from each of the replicates of NeMECiS. Predicted logFC SAG of the WT ordering of fragments that make up the Olig2 CRE is annotated. (D) FL-synCREs from the top 10% of predicted logFC SAG were individually cloned and assayed for reporter activity under 0nM (grey) and 500nM (magenta) SAG. Bars represent means across biological replicates, and points represent individual replicates. (E) As in Fig. 4D but for the bottom 10%. Dim; Diminished. (F) Schematic of the ventral neural patterning gene regulatory network (GRN). Regulatory interactions corresponding to Olig2 CRE are coloured in red. (G) In silico prediction of Olig2 spatial expression pattern when the WT CRE is replaced with either Boost D or Dim D FL-synCREs. (H) Spatial distribution of Olig2 expression patterns from simulations in Fig. 4G. (I) Schematic of GRN rewiring strategy, involving: CRISPR-Cas9 mediated homozygous knockout of the Olig2 gene; re-introduction of a synthetic cassette containing a CRE, a minimal promoter and the Olig2 coding sequence. (J) Olig2 (AlexaFluor 488) mean fluorescence intensity (MFI) for populations of day 6 differentiating neural progenitors derived from FL-synCRE GRN re-wired mESCs, subjected to a logarithmic range of SAG concentrations. Solid lines represent mean across biological replicates, and shaded regions represent maxima/minima (N=3 replicates). (K) As in Fig. 4J, plotted for 500nM SAG. Bars represent means across replicates, and points represent individual biological replicates. (L) Schematic of clonal neural tube organoid (NTO) derivation. Single naïve mESCs are embedded in a Matrigel droplet and resulting NTOs analysed at day 6 of the differentiation. (M) Representative immunofluorescence images of FoxA2 and Olig2 expression in NTOs derived from FL-synCRE GRN re-wired mESCs. Scale bar = 25µm. (N) Quantification of pMN (motor neuron progenitor) domain size for wildtype HM1 mESCs, and mESCs subjected to GRN rewiring using the WT Olig2 CRE and the Boost D FL-synCRE. Dots are replicates. (O) Proportion of voxels that are Olig2 positive, binned by distance from the FoxA2+ pole (see Methods).

Surprised by this observation, we selected a set of candidate FL-synCREs from the top and bottom 10% of the design space to test the predictions (Fig. 4D-E & S11B). Remarkably, we found that re-ordering the fragments of the Olig2 CRE can result in up to a 61% increase in SAG-dependent activity (Fig. 4D). Conversely, we found that candidates in the bottom 10% of activity predictions were able to reduce signal-dependent activity almost completely (Fig. 4E).

### Principled re-engineering of motor neuron progenitor patterning

We investigated whether such ‘optimised’ re-orderings of fragments of the Olig2 CRE could be used to modify Hedgehog signalling responses and reprogramme signal-dependent cell fate transitions. Specifically, natural CREs are embedded within signal-decoding gene regulatory networks (GRNs), hence modifying CRE design enables the creation of synthetic GRNs (synGRNs). As a proof of concept, we asked whether we could predictably rewire the Shh-decoding GRN regulating dorsoventral patterning of spinal cord neural progenitors by placing Olig2, the initiator of motor-neuron development, under FL-synCRE control (Fig. 4F).

We exploited a well-established Ordinary Differential Equation (ODE) model of dorsoventral neural patterning to predict the consequences of GRN rewiring with FL-synCREs (*33*, *66*, *67*). This model integrates both signal strength and spatial distribution to predict how modified CREs alter gene expression patterns across the tissue. We first calibrated the model using fluorescence measurements from FL-synCRE reporters (Fig. S11C), then used the fit parameters to predict the spatial distribution of Olig2 expression upon GRN re-wiring (Fig. S11D). This revealed that replacing Olig2 regulation with a diminished FL-synCREs should reduce the Olig2 expression, shrinking or eliminating the Olig2+ motor neuron progenitor domain, whereas a boosted FL-synCREs should enhance Olig2 expression and expand the domain of Olig2 (Fig. 4G-H & S11E-F). These predictions capture not just changes in expression level, but crucially, how synthetic CREs reshape the tissue-wide organisation of cell types through altered interpretation of the morphogen gradient (Fig. 4G-H & S11D-F).

To test these predictions, we devised a synthetic reconstitution strategy of the cis-regulation of Olig2 to generate synGRNs (Fig. 4I). We made an Olig2-null mutant mESC line, then integrated, with PiggyBac, a cassette driving an Olig2 coding region under the control of a minimal promoter and either WT or FL-synCREs. Differentiating cells under varying SAG concentrations and performing flow cytometry with intracellular staining revealed a rescue of Hedgehog-dependent Olig2 expression (Fig. 4J). Strikingly, under maximum SAG, Olig2 was tuned to different levels depending on the FL-synCRE, with the diminished FL-synCRE showing lower and the boosted FL-synCRE showing higher expression than their wild-type counterpart (Fig. 4K).

These results suggest that neural patterning can be predictably controlled by re-engineering TF control under synCREs. To test this, we generated neural tube organoids (NTOs) (*68*) from synGRN-containing mESCs. Here, single mESCs embedded in Matrigel and cultured appropriately (Materials and Methods) generate neural progenitor cysts, polarised around a central lumen. These self-organise to generate a Shh-producing floorplate that patterns the neural progenitors with ordering and proportions that mirror natural development (*69*) (Fig. 4L). To quantify in an unbiased manner the patterning consequences of GRN rewiring, we performed high-throughput imaging and automated 3D feature extraction using NTOs generated from our synGRN mESC lines (Materials and Methods; Fig. S12A). This revealed altered Olig2 patterning. The reduced FL-synCRE showed a near complete extinction of the Olig2 expressing domain, whereas the boosted FL-synCRE displayed an expanded domain of motor neuron progenitors (Fig. 4M-O). These patterns agree with the *in silico* predictions (Fig. 4G-H). Together, these results demonstrate that reordering of endogenous DNA fragments within a CRE, guided by quantitative design rules, serves as a tunable device for controlling developmental patterning. This finding establishes the programmable control of signal-dependent CRE activity for reshaping tissue pattern and redirecting cell fate decisions.

## Discussion

Precise gene expression responses to external stimuli are a hallmark of cellular information processing. The complexity and robustness of cell-fate decisions that play out during development and homeostasis relies on the capacity of cells to measure subtle changes in signalling. This is typified in morphogen patterning (*2*, *30*). A gradient of a single signalling molecule provides positional information to establish and organise distinct cell fates. CREs mediate the cellular decoding of spatial information by eliciting distinct gene expression programmes, determining when, where, and at which level, genes are turned on and off. Yet how the structure and composition of these sequences achieve precise signal decoding is poorly understood. Uncovering design principles of signal-dependent CRE activity has broad implications, from understanding how natural signal decoding is achieved, to providing insight into the evolutionary constraints on CREs, to applying composability rules to engineer synthetic CREs for programmable control of gene expression Here, we introduce NeMECiS, a novel, high-throughput combinatorial screening strategy to explore systematically the design principles governing CRE function. By enabling the assessment of sequences longer that the typical ∼200bp fragments assayed in many MPRAs (*41–43*), NeMECiS reveals the combinatorial logic of CREs. It bridges the gap between high-throughput reporter assays and more extensive genomic engineering approaches and suggests that the field’s historical focus on shorter fragments may have systematically underappreciated the modular composability rules of CREs. Our findings reveal that CRE activity can be decomposed into modular, interactional, and syntactical components, providing a predictive framework for engineering synthetic regulatory elements and offering new insights into GRNs. We demonstrate that a CRE can be decomposed into a set of modular building-blocks, in which the activity of the collective is the product of its individual components, explaining why individual 200bp fragments of regulatory sequence display minimal activity, yet combine to make functional signal-decoding synCREs. This contrasts with the rigid conformation-dependent model proposed by enhanceosome models and suggests that many CREs operate under a more relaxed set of constraints.

The finding that CRE activity follows a multiplicative rather than additive rule provides mechanistic insight into how regulatory elements achieve signal sensitivity (*49*, *70*). Individual 200bp fragments showed minimal activity, yet their combination generated thousands of functional signal-responsive elements. This multiplicative behaviour explains how weak individual contributions combine to produce strong overall activity, resembling the theoretical predictions of thermodynamic models of transcriptional regulation (*50*, *52–54*).

Some pairs of modules interact synergistically, such that the activity of the pair is higher or lower than the product of its components. This means heterotypic fusions of signal-responsive modules can exceed the activities of their homotypic counterparts, and even that of the full length natural regulatory sequences from which these synCREs were constructed. We also find that interactions among pairs of modules can display distinct spacing-dependent syntax rules, meaning that the ordering of modules in a synCRE can alter its signal-responsiveness. These interactions are crucial for achieving the complex regulatory outcomes observed in GRNs.

The ability to identify and quantify these synergies across a large design space enables the systematic exploration of regulatory interactions that are difficult to predict from sequence information alone. We show how the learned composability rules can be used to rationalise the activity of an endogenous CRE and reprogramme it to increase or decrease its signal-dependence. Such rational design enables the rewiring of signal-decoding GRNs: we use the FL-synCREs to re-engineer signal-dependent control of the motor neuron specifying TF Olig2, producing predictable changes in expression and patterning.

Neural network based models of CRE accessibility and activity (*20*, *62*, *71*) have provided renewed hope for understanding how non-coding sequences enable their cellular functions. High-throughput constructionist methods such as NeMECiS can help test these models beyond the natural training set, allowing one to “hold out the genome” (*24*). However, the challenge in the use of these models is that their size and opacity prohibit the interpretable derivation of design principles. Our simple linear regression model achieves comparable and very similar predictive performance to the more sophisticated neural network models, while offering interpretability. Whereas neural networks remain opaque black boxes, our results demonstrate that CRE activity can be explained through a small set of hierarchical, quantitative rules that provide clear mechanistic insights. This approach is intentionally coarse-grained to map the combinatorial space and hence capture the higher-order composability rules. The equivalent performance of our interpretable framework suggests that the opaque higher-order structures learned by neural network models may simply be approximating these fundamental rules. This has the potential to bridge the gap between black-box predictions and mechanistic understanding, offering a path toward rational design of synthetic CREs.

Synthetic regulatory sequences with programmable signal-responsiveness, cell type specificity and expression levels hold promise in reprogramming cell fates and targeting the delivery of gene therapy cargoes. The modular composability rules our study has uncovered suggest that practices from engineering can be applied to the design of regulatory sequences, allowing for independent optimisation of modules and their integration into bespoke synthetic sequences. This contrasts with the computationally and experimentally intractable task of a complete sequence space search that would be necessary in the absence of modularity. Practically, this will enable the construction of tailored synthetic regulatory sequences by combining the desired parts from a catalogue of modules, which can be adjusted or upgraded as this catalogue expands.

We demonstrate the power of this rule-based engineering approach by rewiring the signal-dependent expression of Olig2, allowing us to reprogramme the pattern of motor neuron progenitors. Moreover, the observation that the rearrangement of natural CREs results in tuneable changes in signal response suggests that evolutionary processes have optimised extant CREs to balance multiple constraints in addition to precisely titrated signal sensitivity to meet tissue patterning needs. Understanding these constraints could provide insight into both the evolution of gene regulation and the design of synthetic regulatory elements for specific applications.

Taken together, our study provides new insight into CRE function and its modular, interactional, and syntactical rules. By systematically exploring the design space of synCREs, we provide a robust framework for the rational engineering of synthetic regulatory elements. These findings have broad implications for synthetic biology and regenerative medicine, offering new avenues for the precise control of gene expression and cellular behaviour. As the field of synthetic genomics continues to progress, the principles uncovered here will serve as a foundation for the next generation of synthetic regulatory tools and therapeutic strategies.

Our approach provides a link between neural network models of CRE function and interpretable design principles, suggesting that a set of simple, quantitative rules can explain how the activity of a CRE depends on its constituent parts. This insight not only advances our understanding of natural regulatory sequences but also opens up new possibilities for engineering synthetic genomes with programmable functions.

## Acknowledgements

We thank Margarida Cardoso Moreira, Zena Hadjivasiliou, Greg Findlay, Jonathan Chubb, Rory Maizels and Hilary Knowles for critically reading the manuscript; Steven Lim, Muhammad Saeed, and the Flow Science Technology Platform for flow cytometry support; Elly Tanaka for advice on the NTO system; Laura Cubitt and the Advanced Sequencing Science Technology Platform for sequencing support; Advanced Light Microscopy Sequencing Science Technology Platform for imaging support; Molly Strom, Ana Cunha and Hilary Knowles for molecular biology advice; Ruben Perez-Carrasco for advice on theory and statistical analysis; Nicholas M. Luscombe for initial discussions; Thomas Frith for advice on mESC maintenance.

## Funding

This work was supported by the Francis Crick Institute, which receives its core funding from Cancer Research UK (CC001051), the UK Medical Research Council (CC001051) and the Wellcome Trust (CC001051); and by the Wellcome Trust (220379/D/20/Z). M.J.D. was supported by the Wellcome Trust Career Development Award (227326/Z/23/Z). J.C.S. was supported by a Boehringer Ingelheim Fonds and Francis Crick Institute PhD Fellowships. D.B (671-2020) and A.P (860-2019) were supported by EMBO ALTFs. G.L.M.B was supported by EMBO ALTF (792-2021) and UKRI (EP/X031225/1). H.T.S was supported by a Sir Henry Wellcome postdoctoral fellowship.

## Author contributions

J.C.S, M.J.D and J.B conceived the project, and acquired funding. J.C.S, D.B, M.J.D and J.B interpreted the data, and wrote the manuscript. J.C.S and D.B performed experiments. J.C.S established theoretical results. J.C.S and A.P performed dynamical modelling simulations. J.C.S, A.P, T.Y, B.D, L.G conceived and performed statistical analyses. A.S generated the Olig2 knockout HM1-TetOn mESC line. S.B. assisted in molecular cloning. E.F assisted with flow-cytometry. G.L.M.B and O.C.K.I assisted in conceptualisation. H.T.S established the NTO protocol. H.T.S. and M.M assisted with its implementation and analysis.

## Competing interests

The authors declare no competing or financial interests.

## Data and materials availability

Code for sequencing processing and statistical analysis is provided at https://github.com/jakesorel/NeMECiS. Accession number for raw and processed barcode sequencing data reported in this paper is GSE291283.

## Notes

### Competing Interest Statement

The authors have declared no competing interest.

