## Supplementary Information for "Predictable Engineering of Signal-Dependent Cis-Regulatory Elements"

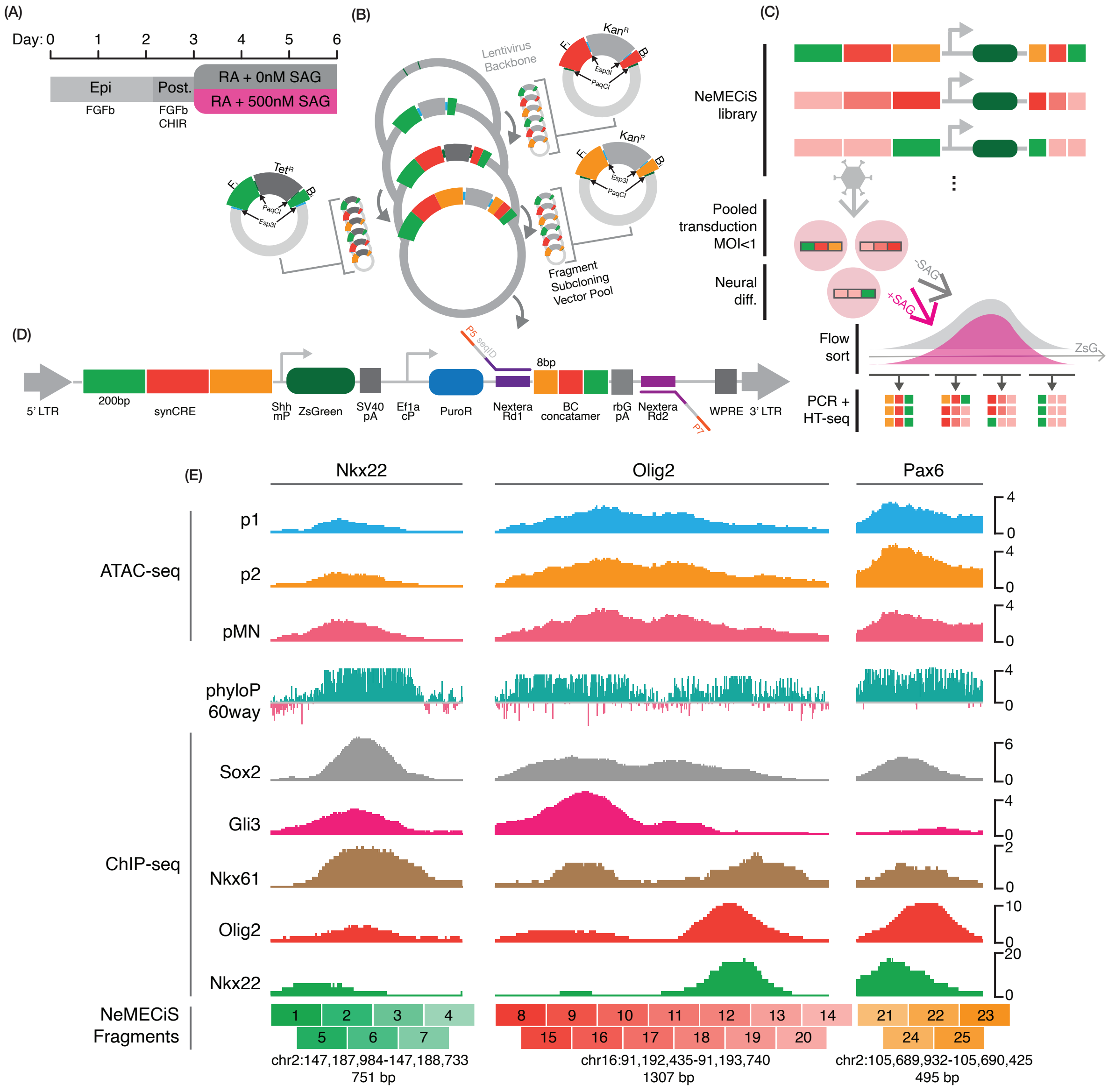


**Supplementary Figure 1. NeMECiS Library Construction Strategy.**

(A) Schematic of neural progenitor differentiation protocol (Epi = epiblast-like state; Post = posterior epiblast-like state; RA = retinoic acid; SAG = smoothened agonist).

(B) Schematic of nested, pooled cloning strategy. Three-part combinatorial synCRE library is cloned in four steps, the first three adding one fragment-barcode pair at a time, and the fourth adding the reporter and selection cassettes. Each step involves cloning a pool of subcloning vectors into the plasmid library by Golden Gate, followed by positive antibiotic selection (KanR = kanamycin resistance; TetR = tetracycline resistance; $F_i$, $B_i$ = CRE fragment "i" and corresponding barcode; Esp3I, PaqCI = TypeIIS restriction enzymes).

(C) Schematic of NeMECiS screening strategy: combinatorial plasmid library is used to generate a lentivirus library, which is integrated into mESCs at MOI<1; selected mESC pool is differentiated to neural progenitors under 0 or 500 nM SAG; then at day 6 is flow sorted into four bins across ZsGreen (ZsG) levels; from which barcodes are amplified from gDNA to quantify ZsGreen levels for all synCREs.

(D) Structure of NeMECiS lentiviral plasmids constructs (LTR = long terminal repeat; Shh mP = Sonic hedgehog minimal promoter; pA = poly-adenylation signal; PuroR = puromycin resistance gene; BC = barcode; rbG = rabbit beta-globin; WPRE = Woodchuck Hepatitis Virus Posttranscriptional Regulatory Element; P5/7 = Illumina flow cell binding sites; seqID = sample barcode for multiplexing sequencing).

(E) Genome-tracks for three neural progenitor patterning CREs (genome coordinates underneath), displaying: day 5 CATS-ATAC-seq for distinct dorsoventral progenitors (*72*); phylogenetic p-values from a 60-way vertebrate alignment (phyloP 60-way) (*75*); ChIP-seq for Sox2, Gli3 (*31*), Nkx61, Olig2, Nkx22 (*74*); and the genomic positions of the 25 NeMECiS fragments.

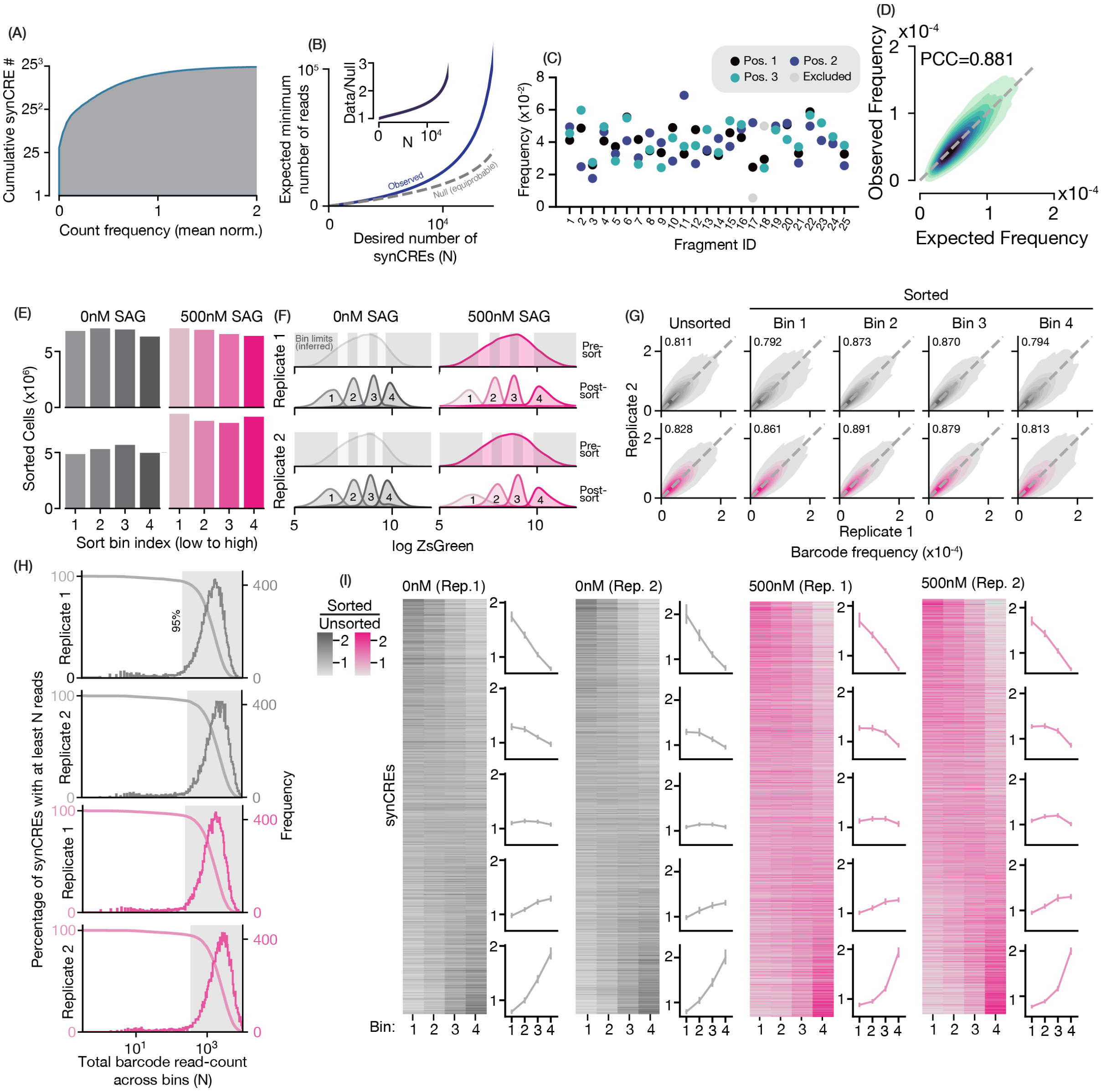


**Supplementary Figure 2. Quality Control and Data Processing for NeMECiS Library.**

(A) Cumulative barcode frequencies from amplicon sequencing of the NeMECiS plasmid library (theoretical max barcode number = 15,625) as a function of barcode count frequencies, normalised to its average (mean. norm).

(B) Simulation of minimum number of reads required to recover a given number of barcodes (coupon collector's problem) on measured barcode frequencies in the plasmid library, compared to a null model where barcodes are equiprobable (dashed line), indicating library evenness. Inset: A ratio of the expected number of reads required to recover a given number of barcodes given the measured distributions with respect to the null.

(C) Frequencies of each fragment in each position (colours), demonstrating even distribution of barcodes.

(D) Comparison of observed barcode frequencies against their frequencies expected from random sequential addition predicted from the cloning strategy (see Fig. S1B; i.e. frequency of synCRE ABC = product of frequencies of fragments A,B,C in positions 1,2,3 respectively; PCC = Pearson's correlation coefficient).

(E) Number of sorted cells, by sort bin, SAG concentration and replicate.

(F) By replicate and SAG concentration, ZsGreen distribution (live flow-cytometry, kernel density estimate), in the unsorted (pre-sort) day 6 NeMECiS neural progenitor pool, and in each of the four bins (post-sort), reporting sorting purity. Set on top are the estimated sort-bin extents (see Methods).
(G) Reproducibility over replicates, across SAG concentrations, of barcode frequencies in the unsorted pool and in each of the four sorted bins.
(H) Barcode recovery in sorted bins. For a given SAG concentration and replicate, taking the total number of reads for each barcode across bins, the inverse cumulative frequency (lighter colour, left axis) and the read-count histogram (darker colour, right axis) is plotted. 5$^{th}$ percentile of total barcode counts (i.e. 95$^{th}$ percentile of barcode recovery) is indicated.
(I) Barcode frequency by bin, normalised to unsorted barcode frequency, is plotted, sorted by the average median bin between replicates, for each SAG concentration. To the right of each heatmap, mean normalised barcode frequencies by bin for quintiles of the average median bin (error bars = 95% confidence interval of the estimate of the mean).

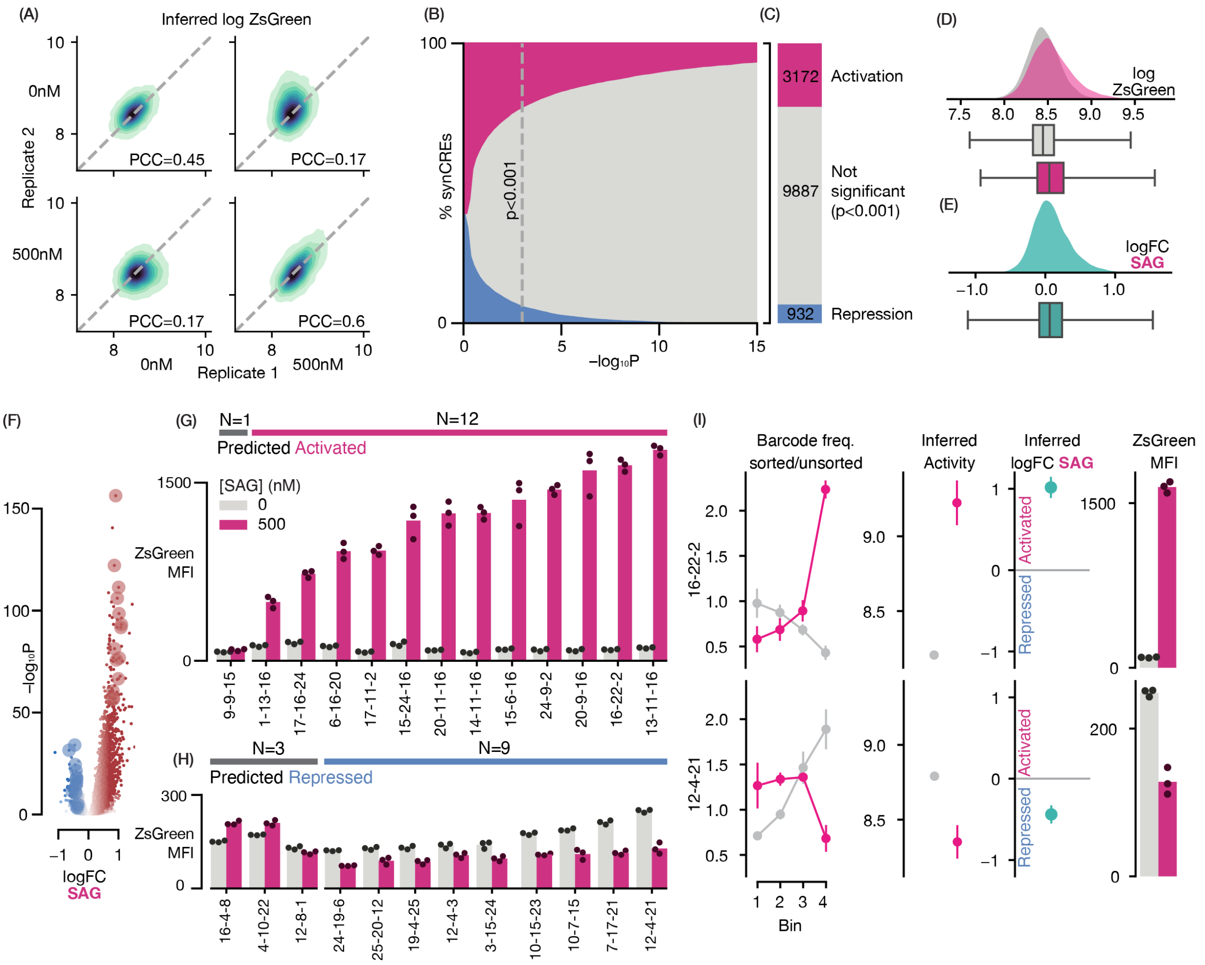


**Supplementary Figure 3. Distribution of Signal-Dependent Activities Across synCREs.**
(A) Comparing inferred log ZsGreen from Bayesian inference of barcode frequencies for the NeMECiS synCRE library across replicates and SAG concentrations, demonstrating reproducibility across replicates for the same (but not different) SAG concentration (PCC = Pearson's correlation coefficient).
(B) Frequency of synCREs determined to display activation (magenta) or repression (blue) for increasingly stringent significance cut-offs (one-tailed Z-test).
(C) A cross section of Fig. S3B for P<0.001 (indicated by the grey dashed line), enumerating synCREs displaying activation and repression.
(D). Distribution of inferred log ZsGreen levels across the NeMECiS synCRE library (grey = 0nM SAG; magenta = 500nM SAG), plotting kernel density estimates, and box-plots underneath.
(E) As above for inferred log fold change of inferred log ZsGreen levels with respect to SAG (logFC SAG).
(F) Volcano plot of logFC SAG against significance (calculated as above). Chosen validation constructs tested in Fig. S3G-H are encircled.
(G) Validation constructs built and assayed individually as in Fig. 1A tested for ZsGreen levels (MFI = mean fluorescence intensity) for 0nM and 500nM SAG. Bars indicate mean across replicates, and points indicate biological replicates (N=3). Above, the number that showed the expected behaviour are reported.
(H) As above, but for repressed validation constructs.
(I) Two case-studies of the workflow of data processing and validation for an activated (above, f16-f22-f2) and repressed (below, f12-f4-f21) synCRE. From left to right: frequencies of barcode reads by bin, normalised by barcode frequency in the unsorted pool (as in Fig. S2I; points indicate mean

across replicates, error bars indicate the maximum and minimum); inferred activity (i.e. log ZsGreen) of each synCRE for each SAG concentration (gray vs. magenta) from Bayesian inference; log fold change of ZsGreen with respect to SAG (logFC) computed from inferred activities; independently validated synCREs (repeating data from Fig. S3G-H for comparison).

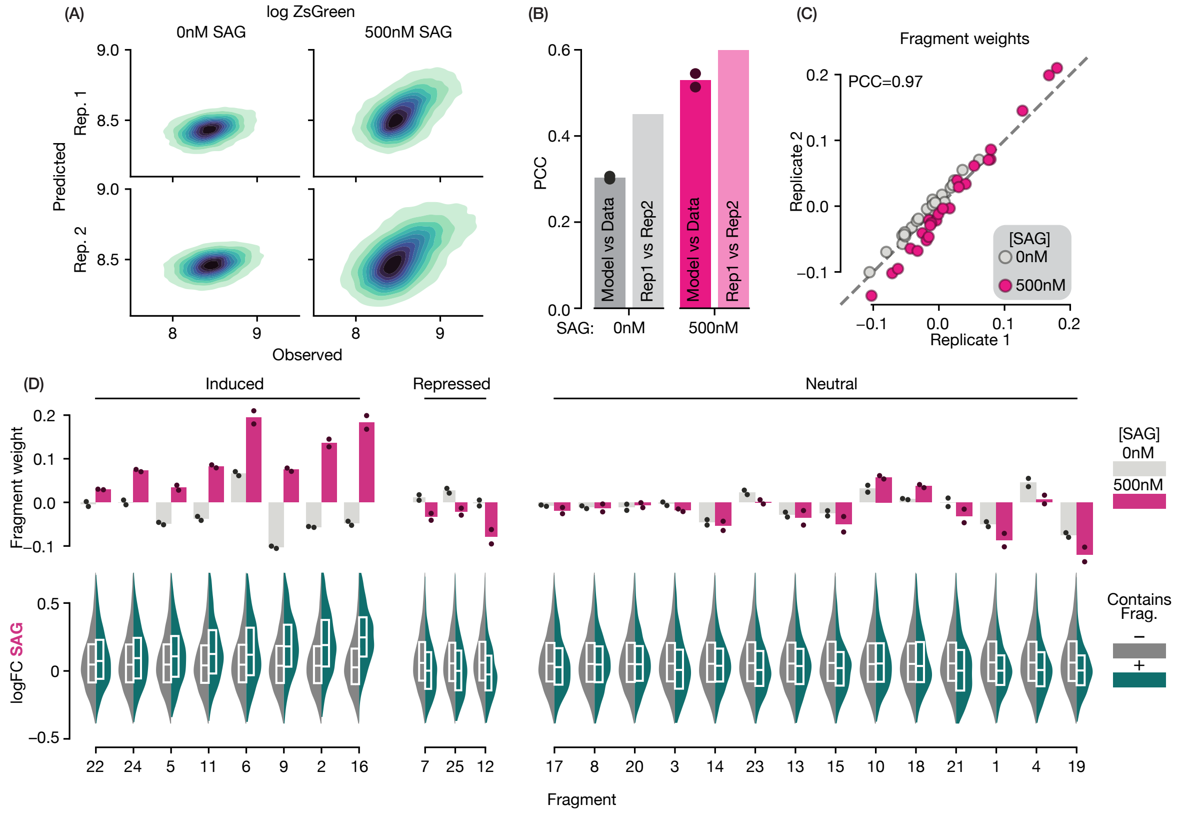


**Supplementary Figure 4. Linear Modelling Framework for Fragment Activity Analysis.**

(A) Comparison of observed versus predicted (under linear modelling) of log ZsGreen for all synCREs for 0nM and 500nM SAG, for each replicate (Rep.).

(B) Corresponding Pearson's correlation coefficients (PCC) between the linear modelling predictions and measurements (darker colours, bars represent means and points represent replicates). To benchmark these values, next to each the PCC of measured log ZsGreen between replicates is plotted.

(C) Predicted weights arising from linear modelling from each fragment, across replicates (dashed line diagonal corresponds to identity between replicates), coloured by SAG concentration.

(D) Above: fragment weights for 0nM and 500nM SAG (bar represents mean across replicates, and points represent each replicate). Below: violin-plots and associated quartiles for logFC SAG (log fold change of ZsGreen with respect to SAG) for all synCREs without (grey) or with (teal) a particular fragment, associated with Fig. 2B.

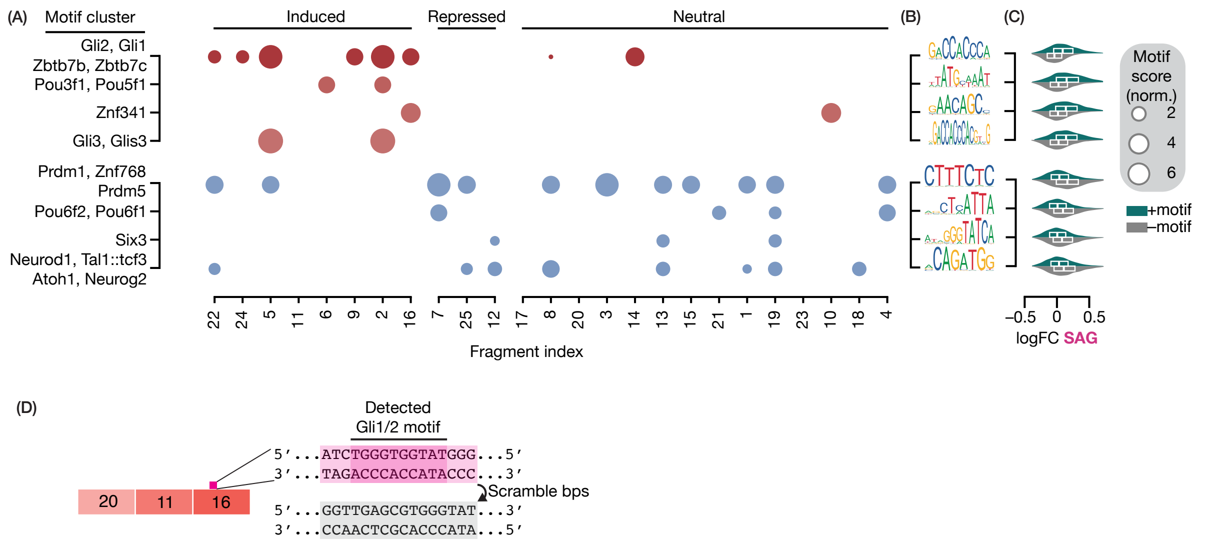


**Supplementary Figure 5. Sequence Motif Analysis of Hedgehog-Responsive Modules.**
(A) Dot-plot of motif clusters with the four most positive and four most negative Spearman's correlation coefficient in Fig. 2C, across all fragments, grouped by fragment class (induced, repressed, neutral). The presence of a dot indicates a motif of the corresponding motif cluster is found in the corresponding fragment (threshold MOODS hit-score = 6.5), with the diameter of the dot indicating how much that maximum MOODS hit-score exceeds the threshold (Motif score norm. in the legend).
(B) Corresponding logo-plot of the position weight matrices of a representative motif from each motif cluster (first motif listed in the left-hand key is plotted).
(C) Corresponding violin-plot and associated quartiles of the log fold change of ZsGreen with respect to SAG (logFC SAG) for all synCREs without (grey) or with (teal) an instance of the motif cluster.
(D) Schematic describing the Gli1/2 motif binding site mutagenesis strategy: the single MOODS-called motif in f20-f11-f16 (dark-magenta) is expanded by 3bp each side (light-magenta), and then base-pairs are scrambled (grey colouring beneath).

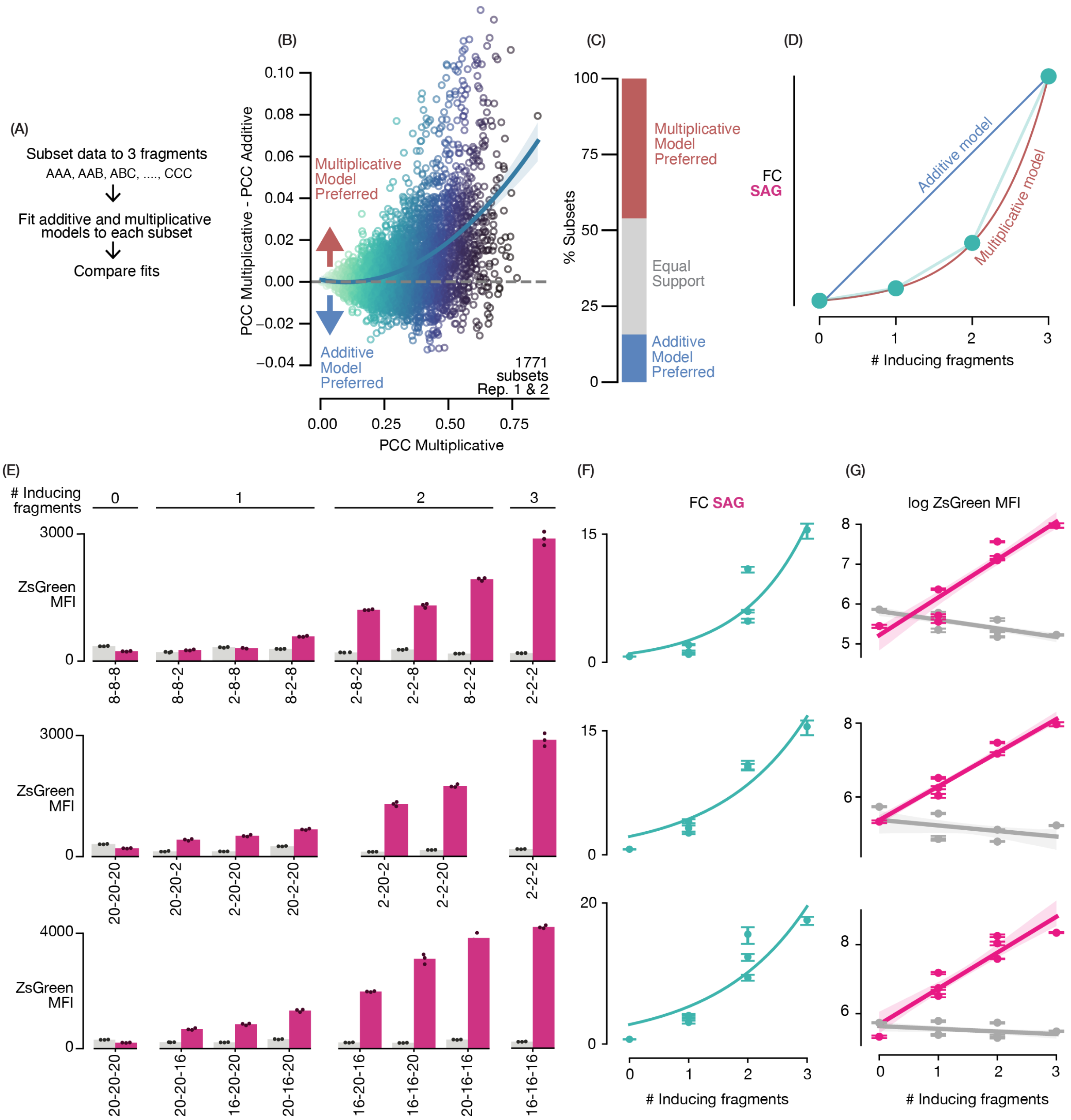


**Supplementary Figure 6. Validation of the Multiplicative Model for Fragment Combinations.** (A) Workflow of systematic comparison of additive and multiplicative models of modular composability. Fits are performed on linear-transformed inferred ZsGreen fold-changes without regularisation, with the multiplicative model fit with non-linear optimisation.

(B) Comparison of additive and multiplicative modular composability model fits (PCC = pearson's correlation coefficient), plotting the difference in their respective PCCs marginalised against the PCC of the multiplicative model fit. Each point corresponds to a three-fragment subset of the NeMECiS dataset for either replicate. Overlaid is the rolling mean (quadratic fit, error bars = 95% confidence interval).

(C) Proportion of three fragment subsets for which the multiplicative or additive model show better performance across biological replicates (equal support = preferred model across replicates diverge).

(D) Schematic of additive and multiplicative model predictions across exhaustive combinations of activating and neutral fragments.

(E) Individually built and assayed combinations of activating (f2, f16) and neutral (f8, f20) fragments, lending support for the multiplicative model, equivalent to Fig. 2I.
(F) Corresponding fold change of mean fluorescence intensity of ZsGreen with respect to SAG (FC SAG), grouped by the number (#) of inducing fragments.
(G) Corresponding log ZsGreen mean fluorescence intensities under 0nM (grey) and 500nM (magenta) SAG.

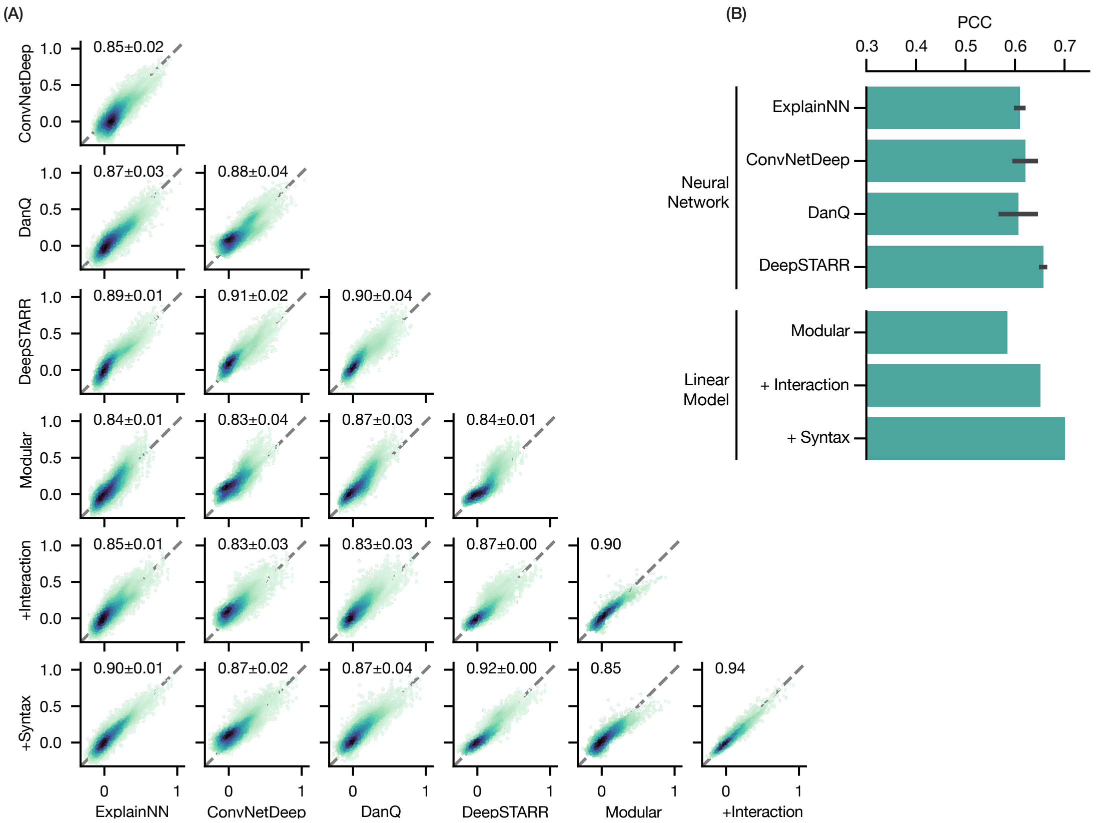


**Supplementary Figure 7. Comparison of Neural Network and Linear Models for Predicting synCRE Activity.**

(A) Comparison of predicted mean log fold change of ZsGreen with respect to SAG among models. For neural network models, data is stratified into test, train and validation, across five independent samplings and trainings. For linear models, the predictions for replicate 1 and 2 are averaged. All test data are plotted across sampling replicates. Mean pearson correlation coefficients across sampling replicates are reported (± standard deviation).

(B) Pearson correlation coefficient (PCC) of each model to the data. For neural network models, the mean PCC across sampling replicates of the test data is reported (± 95% CI).

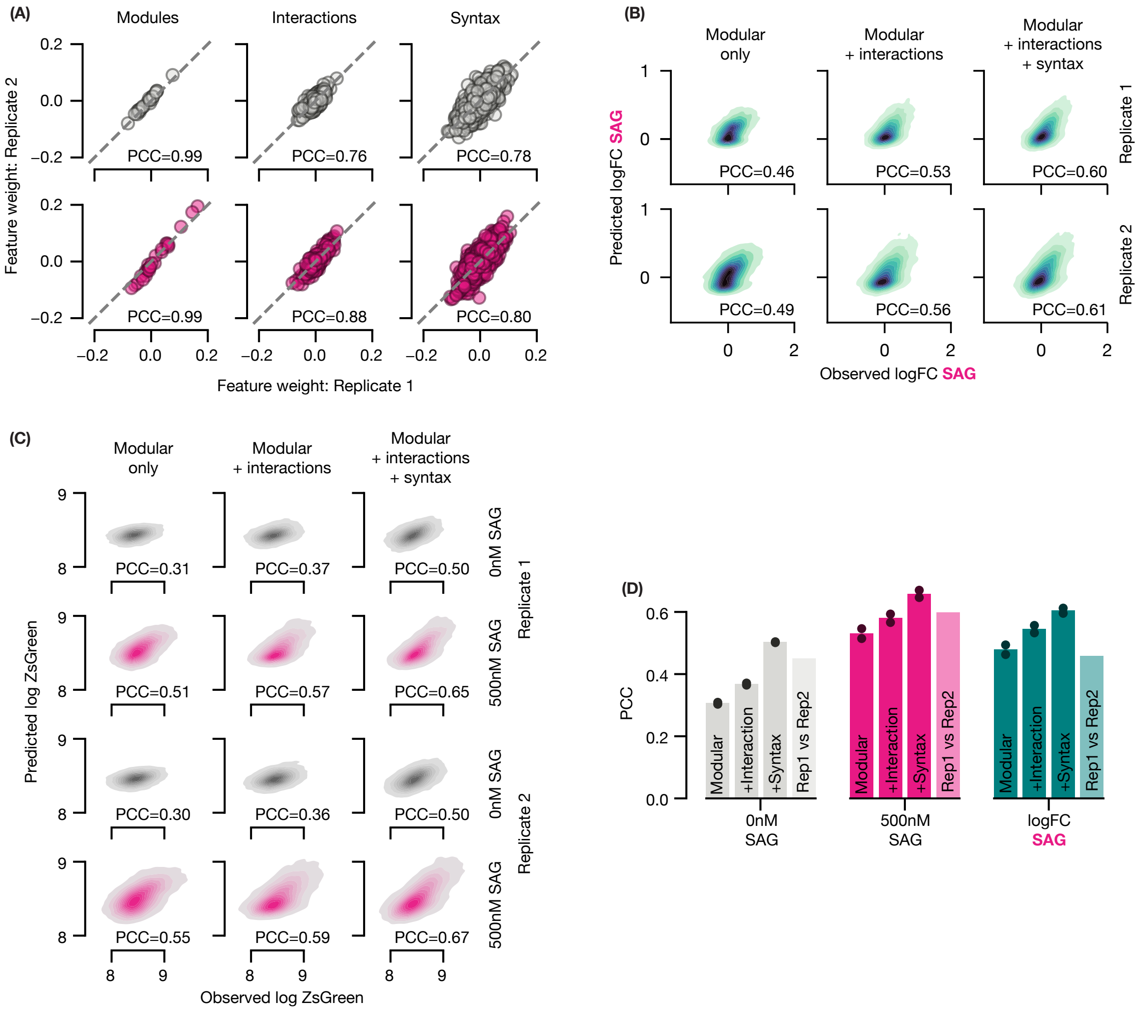


**Supplementary Figure 8. Interaction and Syntax Effects Across Replicates.**

(A) Reproducibility of feature weights from extended synergistic linear model, by feature class and SAG concentration (0nM = grey; 500nM = magenta).

(B) Observed versus predicted log fold change of ZsGreen with respect to SAG (logFC SAG) for full model (right-most) versus sequential parameter marginalisation to remove syntactical variance (middle) and interactional variance (left), by replicate. Pearson's correlation coefficient (PCC) for each is reported.

(C) As in Fig. S8B but reported individually for 0nM and 500nM SAG, plotting observed versus predicted log ZsGreen.

(D). Bar plot of PCCs for full model ("+syntax") and sequential truncations, for 0nM, 500nM and logFC SAG. For comparison the PCC between replicates for each measure is reported (lighter shade).

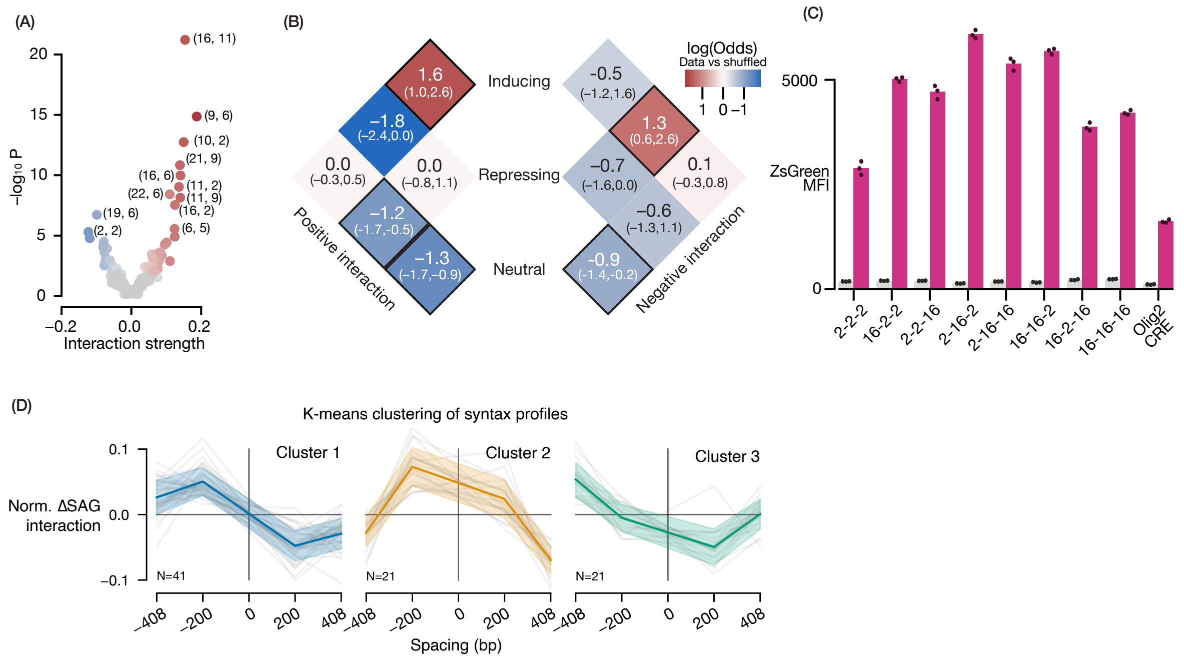


**Supplementary Figure 9. Characterisation of Pairwise Fragment Interactions and Syntax Dependencies.**

(A) Volcano plot testing for non-zero values of interaction coefficients (z-test on bootstraps).

(B) Permutation test for the distribution of significant (P<0.01) positive and negative interactions among fragment classes. An ensemble of randomised contact matrices was produced, then the distribution of interactions compared to the average of this null distribution were used to calculate a (continuity corrected) log Odds ratio. (ranges beneath each annotation define 5$^{th}$ and 95$^{th}$ percentiles of the log(odds).

(C) Individually built and assayed exhaustive combinations of two activating fragments (f2, f16), compared to the full Olig2 CRE. Plotted as in Fig. 2I. Corresponds to Fig. 3D. (D) K-means clustering (n=3) of spacing profiles of logFC SAG (normalised by its spatial mean) for heterologous pairs of fragments where the corresponding positionality score (Fig. 3E) exceeds 2.

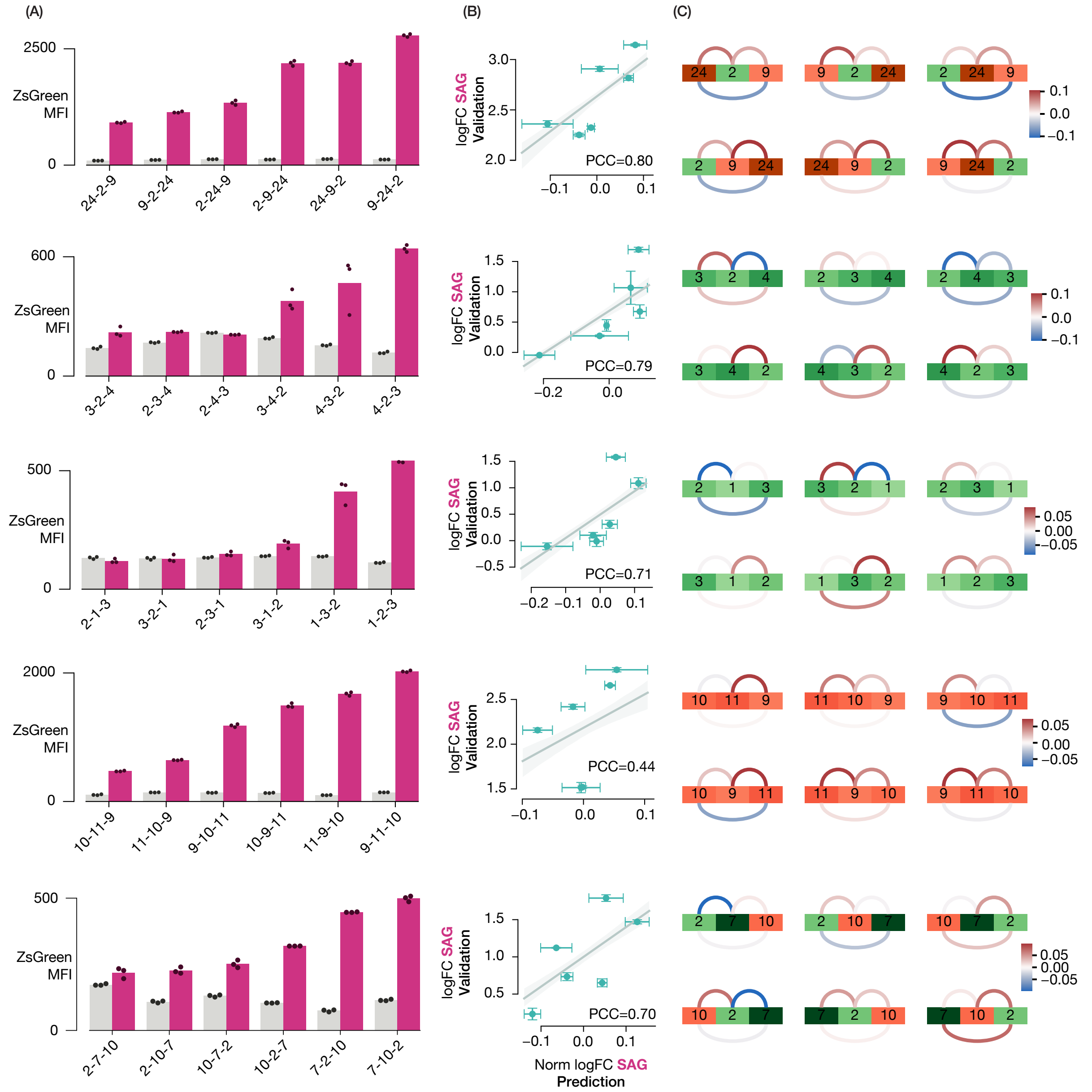


**Supplementary Figure 10. Order-Dependent Synergies in Fragment Combinations.**

(A) Individually built and assayed exhaustive re-orderings of heterologous trimers, plotted as in Fig. 2I.

(B) Comparison of predicted vs observed logFC SAG, plotted as in Fig. 3K.

(C) Syntax contributions to logFC SAG for exhaustive re-orderings, plotted as in Fig. 3J.

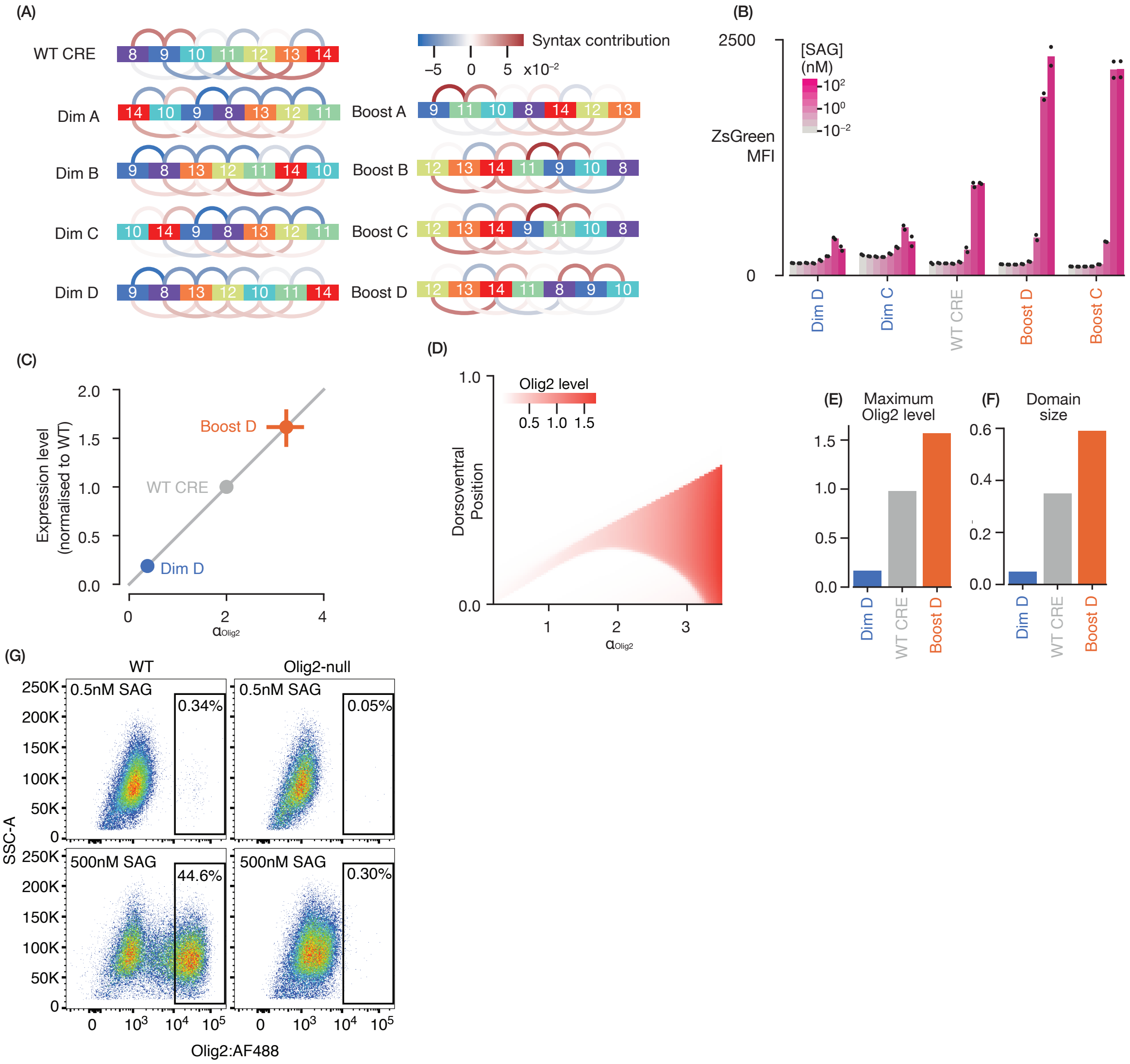

**Supplementary Figure 11. Computational Predictions for FL-synCRE Activities.**

(A) Syntax contributions for each pair of fragments in the WT Olig2 CRE, and re-orderings predicted to diminish or boost signal-responsiveness. Plotted as in Fig. S10C.

(B) SAG serial dilution (8 concentrations at 5-fold dilution starting at 500nM; performed and plotted as in Fig. 1H) for reporters of the WT Olig2 CRE and two representative re-orderings that boost or diminish signal responsiveness.

(C) Simulation output of a GRN model of neural tube patterning, adapted to measure relative reporter activity. Grey line plots the expected reporter level at 500nM SAG (see Methods) across a parameter sweep of $\alpha_{Olig2}$, a parameter controlling the propensity of bound transcriptionally activating transcription factors to initiate transcription. Overlaid points and error bars are the inferred parameter values for the FL-synCREs Dim D and Boost D (fit using 500nM SAG data from Fig. 4D-E).

(D) Parameter sweep for $\alpha_{Olig2}$, plotting the dorsoventral expression pattern of Olig2 (y-axis) for each parameter value (x-axis).

(E) Measured maximal Olig2 level when the simulated Olig2 CRE is swapped out with either Dim D and Boost D FL-synCREs (using fits from Fig. S11C). Corresponds to Fig. 4G.

(F) Corresponding domain sizes (defined as the dorsoventral extent for which the Olig2 level exceeds half of its maximum).

(G) Fixed and intracellularly stained flow cytometry data for 0.5nM and 500nM SAG differentiation, filtered by Sox2+ live neural progenitors, plotting Olig2 (AF88) mean fluorescence intensity (MFI), for WT and Olig2-null mESC lines.

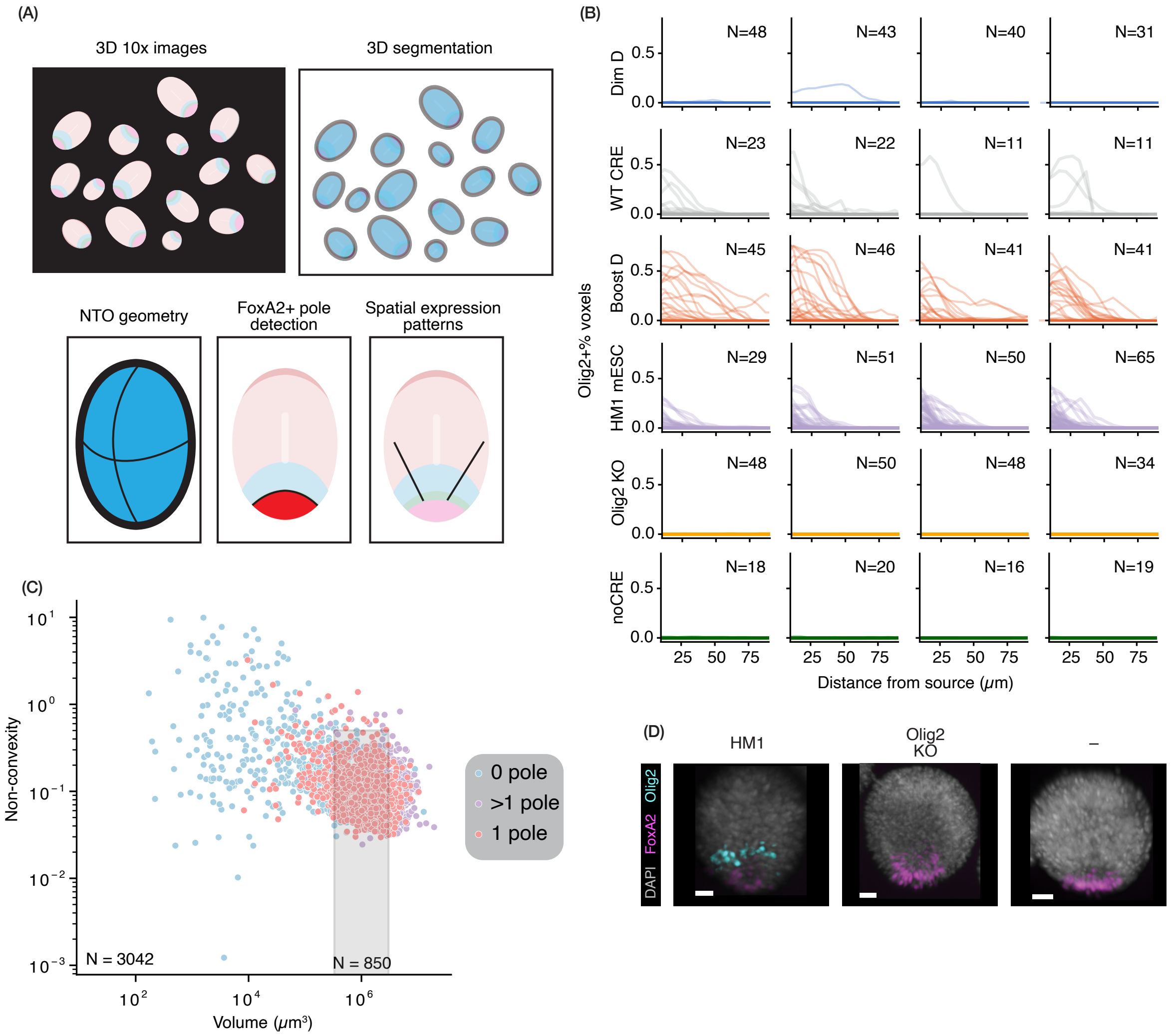


**Supplementary Figure 12. 3D Feature Extraction and Analysis of Neural Tube Organoids.**
(A) Schematic for the 3D neural tube organoid (NTO) segmentation and spatial analysis pipeline.
(B) Spatial distribution of the percentage of Olig2 positive voxels at different binned distances from the FoxA2+ source, for each cell line (rows) and for each of the four replicates (columns).
(C) NTO subsetting for downstream analysis. Each point represents a single segmented NTO, plotted with respect to their volume ($\mu m^3$) and non-convexity (i.e. volumetric ratio with respect to convex hull), coloured by the number of FoxA2+ poles. Red points lying in grey area represents samples used in downstream analysis.
(D) Representative 3D projections of NTOs from the following mESC lines: HM1 (wildtype); Olig2 knockout; minimal promoter only (noCRE) . Scale bar = 25$\mu$m.
