## Supplementary material for "Predictable Engineering of Signal-Dependent Cis-Regulatory Elements": Materials and Methods

### Predictable Engineering of Signal-Dependent Cis-Regulatory Elements
### Materials and Methods

**13 Tables** **32**

### 1 Cell culture

#### 1.1 Mouse ES cell maintenance

All experiments were carried out using the HM1 mouse embryonic stem cell (mESC) line [Magin et al., 1992], maintained at 37°C with 5% CO2. For all experiments besides the generation of neural tube organoids (NTOs), mESCs were maintained in ESGRO Complete PLUS Clonal Grade Medium (Merck Millipore, Cat. No. SF001-500P) supplemented with 1% Penicillin/Streptomycin (Gibco, Cat. No. 15140122) on CellBIND tissue-culture plates (Corning, Cat No. 3335) coated with 0.15% gelatin (Gibco, Cat. no. G1393-100ML). mESCs were passaged every two days at $10^4$ cells / $cm^2$ using StemPro Accutase (Gibco, Cat. No. A1110501) to dissociate cells. For generating NTOs, mESCs were maintained in 2i+LIF media at $3 \times 10^3$ cells / $cm^2$: DMEM/F12(-/-) (Gibco, Cat. No. 21331-020) and Neurobasal (Gibco, Cat. No. 21103-049) at 1:1, supplemented with 1xN2 (Gibco Cat No. 17502001), 1xB27 (Gibco Cat. No. 17504001), 0.1% 50mM 2-mercaptoethanol (Gibco, Cat. No. 31350-010), 1% 200mM L-Glutamine (Gibco, Cat. No. 25030024), 1% Penicillin/Streptomycin (Gibco, Cat. No. 15140122), 500U/ml LIF (Chemicon, Cat. No. Int ESG1107), 1$\mu$M PD0325901 (Axon Medchem, Cat. No. 1408), 3$\mu$M CHIR99021 (Axon Medchem, Cat. No. 1386).

#### 1.2 Olig2 CRISPR knockout HM1 line generation

A gRNA expression plasmid was built using the pX459v2 vector (Addgene #62988) and the following complementary 5' pre-phosphorylated oligonucleotides (CACCgTTTCTGCCCGCCCGGAGCAA, AAACTTGCTC-CGGGCGGGCAGAAAc, Integrated DNA Technologies), following [Ran et al., 2013]. 4$\mu$g of a maxiprep (Qiagen, Cat. No. 12162) of the expression plasmid was electroporated into 2i+LIF cultured HM1-TetON mESCs [Serafimidis et al., 2008] with the AMAXA nucleofector kit (Lonza Cat no. VPH-1001) following manufacturer's instructions, and plated onto 0.1% gelatin coated 10cm tissue culture plates. After 24 hours, media was changed into 2i+LIF supplemented with 1.5$\mu$g/mL Puromycin Dihydrochloride (Gibco, Cat. No. 12122530). After a further 48 hours, media was changed to 2i+LIF. Once visible, clones were expanded and genotyped (F:atggactcggacgccagc; R:gacggtgacttgagcaggc). An homozygous knockout clone was identified to carry a 23bp deletion (CGGAGCAAGGGGGGAAGCAGCAG), resulting in a 2bp frameshift after 24 amino acids, and a premature stop codon after 94 amino acids. Knockout was confirmed at the protein level by fixed intracellular flow-cytometry upon 6-day differentiation (Goat Olig2 unconjugated, 1:400; R&D, Cat. No. AF2418) (Fig. S11G).

#### 1.3 Lentivirus generation

To generate lentivirus, HEK293T were plated at 1.5x$10^6$ per 60mm tissue culture plate, cultured in DMEM (Gibco, Cat. No. 21969-035), supplemented with 10% Fetal Bovine Serum (PAN-Biotech P30-3031), 1% 200mM L-Glutamine (Gibco, Cat. No. 25030024), 1% Penicillin/Streptomycin (Gibco, Cat. No. 15140122). The next day, the media was changed (3.5mL). 3rd generation lentivirus packaging plasmids (0.24$\mu$g CMV-Rev, 0.46$\mu$g pMDLg, 0.34$\mu$g VSV-G) and 2.28$\mu$g of the transfer plasmid were mixed with 360$\mu$L Opti-Mem (Gibco, Cat. No. 31985062), 11.9$\mu$L XtremeGene-HP (Roche, Cat. No. 6366236001), vortexed briefly and incubated at room-temperature for 15 minutes. This was added dropwise to the HEK293T cells. 16 hours later, the media was changed into N2B27 media (2mL): DMEM-F12 (Gibco, Cat. No. 21331-020) and Neurobasal (Gibco, Cat. No. 21103-049) (1:1), supplemented with 1xN2 (Gibco, Cat No. 17502001), 1xB27 (Gibco, Cat. No. 17504001), 2mM L-glutamine (Gibco, Cat. No. 25030024), 40$\mu$g/ml BSA (Sigma-Aldrich, Cat. No. A7979-50ML), and 0.1mM 2-mercaptoethanol). After 30 hours, the supernatant was collected with a syringe and filtered through a 0.45$\mu$m filter. Lentivirus preps were scaled up and down by culture plate area. To generate stable mESC lines, lentivirus supernatant was added to mESCs at the time of passaging at MOI<1; after 48hrs and from then onwards, cells were maintained with 1$\mu$g /mL Puromycin Dihydrochloride (Gibco, Cat. No. 12122530).

#### 1.4 Piggybac transgenesis

To generate Piggybac transgenic mESC lines, 0.1$\mu$g of a mouse-codon optimised PiggyBac transposase plasmid [Cadinanos and Bradley, 2007] was vortexed in a 1.5mL Eppendorf microtube with 0.4$\mu$g of PiggyBac transfer plasmid, 25$\mu$L Opti-Mem (Gibco, Cat No. 31985062). In a separate microtube, 1$\mu$L of

Lipofectamine Stem Transfection Reagent (ThermoFischer, Cat. no. STEM00001) was vortexed with 25$\mu$L Opti-Mem, which were combined together and incubated for 10 minutes. mESCs were passaged in ES-GRO Complete PLUS Clonal Grade Medium (Merck Millipore, Cat. No. SF001-500P) onto 24-well tissue culture plates coated with 1% Vitronectin (Gibco, Cat. No. A31804), and the transfection mixture was added dropwise. After 48 hours, cells were maintained in 10$\mu$g/mL Blasticidin S HCl (Gibco, Cat. No. A1113903), including during 2D and 3D differentiations.

#### 1.5 2D neural progenitor differentiation

##### 1.5.1 Dual-colour differentiation for live flow cytometry

10cm tissue culture plates were coated with 1% Vitronectin (Gibco, Cat. No. A31804) in Phosphate Buffer Saline (PBS) (Gibco, Cat. No. 14190-094) for 1 hour. mESCs were dissociated using StemPro Accutase (Gibco, Cat. No. A1110501) and centrifuged and resuspended in N2B27 (see above). Cells were seeded at $3 \times 10^5$ cells per plate (8mL) supplemented with 10ng/ml bFGF (R&D, Cat. No. 100-18B). Two days later, 12-well tissue culture plates were coated for 1 hour with a 1:100 dilution of GelTrex (Gibco, Cat. No. A1413202) in DMEM-F12(-/-) (Cat. No. 21331046). After 48hrs, cells were dissociated with StemPro Accutase (Gibco, Cat. No. A1110501), centrifuged and resuspended in N2B27 supplemented with 10ng/ml bFGF (R&D, Cat. No. 100-18B) and 5$\mu$M CHIR99021 (Axon Medchem, Cat. No. 1386) and plated at a 1x 10cm plate to 1x 12 well-plate ratio (0.95mL media per well), supplemented additionally with 50$\mu$L lentivirus supernatant. 20 hours later, media was changed into N2B27 supplemented with 100nM RA (Sigma, Cat. No. R2625) and variable concentrations of SAG (Merck, Cat. No. 566660-5mg). Media was changed daily. 72 hours later (day 6 of the differentiation), cells were collected for flow cytometry analysis (see below).

##### 1.5.2 Neural progenitor colony differentiation imaging

The differentiation was set up in an identical manner to the above, but in 12-well tissue culture plates, plating $2.1 \times 10^4$ cells per well. After 48 hours, media was changed into 10ng/ml bFGF (R&D, Cat. No. 100-18B) and 5$\mu$M CHIR99021 (Axon Medchem, Cat. No. 1386). After 20 hours, media was changed into 100nM RA (Sigma, Cat. No. R2625) and variable concentrations of SAG (Merck, Cat. No. 566660-5mg). The next day, 24-well imaging plates (Miltenyi Biotec, Cat. No. 130-098-263) were coated with a 1:50 dilution of Matrigel (Corning, Cat. No. 356231) in DMEM-F12 (-/-) (Gibco, Cat. No. 21331-020). After 24 hours of further culture, cells were dissociated with StemPro Accutase (Gibco, Cat. No. A1110501) and resuspended in fresh N2B27 with the same concentrations of supplements, and re-plated from a 1$\times$ 12-well well to 8$\times$ 24-well wells, which were pre-coated for 1 hour with with 1:100 GelTrex (Gibco, Cat. No. A1413202) in DMEM-F12 (Gibco, Cat. No. 21331046), to obtain sparse colonies. Media was changed daily. After a further 48 hours, samples were washed twice with Phosphate Buffer Saline (PBS) (Gibco, Cat. No. 14190-094) and fixed for 10 minutes with 4% methanol-free paraformaldehyde (ThermoScientific, Cat. No. 28908), followed by two subsequent washes in PBS (Gibco, Cat. No. 14190-094). Cells were then prepared for immunofluorescence staining (see below).

##### 1.5.3 Intracellular fixed flow-cytometry

6-well tissue culture plates were coated for 1 hour with a 1:50 dilution of Matrigel (Corning, Cat. No. 356231) in DMEM-F12 (-/-) (Gibco, Cat. No. 21331-020). mESCs were dissociated and resuspended in N2B27 supplemented with bFGF as above and plated at $2 \times 10^4$ cells per well. Media was changed 48 hours later into bFGF and CHIR99021 as above. After 20 hours, media was changed into 100nM RA and variable SAG as above. Media was changed daily for another 72 hours, then cells were fixed for intracellular-stained flow cytometry (see below)

#### 1.6 Neural tube organoid differentiation

Following the protocol for neural tube organoid generation [Krammer et al., 2024], mESCs maintained in 2i+LIF were dissociated with 200$\mu$L StemPro Accutase (Gibco, Cat. No. A1110501), and resuspended in 2mL 'Advanced N2B27' (Advanced DMEM/F12 (Gibco, Cat. No. 12634-010) and Neurobasal (Gibco, Cat. No. 21103-049) at 1:1, supplemented with 1xN2 (Gibco Cat., No. 17502001), 1xB27 (Gibco Cat. No. 17504001), 0.1% 50 mM 2-mercaptoethanol (Gibco, Cat. No. 31350-010), 1% 200mM L-Glutamine

(Gibco, Cat. No. 25030024), 1% Penicillin/Streptomycin (Gibco, Cat. No. 15140122), 1% non-essential amino acids (Gibco, Cat. No. 11140-035). Cells were aliquoted into a new 15mL Falcon, centrifuged, aspirated dry and gently resuspended in ice-cold Matrigel at a concentration of $10^2$ cell $/\mu$L (Corning, Cat. No. 354234). In small batches, 4$\mu$L of the mix was plated per well of an optically clear flat-bottom 96-well imaging plate (PerkinElmer Cat. No. 6055302) by drawing an inwards spiral in the centre of each well, ensuring a flat droplet of Matrigel that does not touch the sides. Plates were incubated for 3 minutes in a 37℃ incubator, then supplemented with 100$\mu$L of Advanced N2B27. 48 hours later, 250nM all-trans Retinoic Acid (Merck, Cat. No. R2625) was supplemented into the media, and replaced with unsupplemented Advanced N2B27 after 18 hours. Media was changed daily. After 6 days from plating, 100$\mu$L of 4% paraformaldehyde (made from powder, as in [Krammer et al., 2024]) was added to each well on top of existing media and incubated at room temperature for 30 minutes. Wells were then washed twice with PBS.

### 2 Flow cytometry

#### 2.1 Live flow cytometry

For live-cell flow cytometry collection, 1$\mu$L of LIVE/DEAD Fixable Dead Cell Stain Near-IR fluorescent reactive dye (ThermoScientific, Cat. No. L34976) was added to each 12-well tissue-culture plate well (1:1000) dilution and incubated at 37℃ for 30 minutes. Wells were then washed twice with PBS, followed by dissociation with 100$\mu$L StemPro Accutase and resuspension in 120$\mu$L PBS+0.5% BSA (Sigma-Aldrich, Cat. No. A7979-50ML). Cells were transferred to a LoBind Microplate 96/V-PP 96-well plate (Eppendorf, Cat. No. 0030603303) and centrifuged at 500g for 5 minutes. Supernatant was removed by inversion of the plate. Cells were resuspended in 40$\mu$L of PBS + 2mM EDTA +0.5% BSA. Samples were run in high-throughput mode either on the BioRad ZE5 Cell Analyzer or the BD LSRFortessa Cell Analyzer, analysing the entire well for each. Samples that were infected with lentiviruses expressing ZsGreen or mScarlet3 singly and constitutively were run for compensation (without LIVE/DEAD staining), as were uninfected cells stained with LIVE/DEAD.

#### 2.2 Fixed flow cytometry

For fixed intracellular-stained flow cytometry collection, following [Delas et al., 2023], LIVE/DEAD Fixable Dead Cell Stain Near-IR fluorescent reactive dye (ThermoScientific, Cat. No. L34976) was incubated at a 1:1000 dilution at 37℃, then washed, as above. Cells were then dissociated with StemPro Accutase and resuspended in PBS + 0.5% BSA as above. Cells were then transferred into 1.5mL DNA LoBind Eppendorf microfuge tubes (Eppendorf, Cat. No. Z666548), centrifuged at 500g for 5 minutes, supernatant removed by aspiration, and resuspended in 100$\mu$L of 4% methanol-free paraformaldehyde (ThermoScientific, Cat. No. 28908). After a 10 minute room-temperature incubation, 1mL of PBS was supplemented to each tube, followed by a 5 minute centrifugation at 2000g, and subsequent aspiration of supernatant. Cells were then resuspended in PBS + 0.5% BSA and stored at 4℃. For subsequent intracellular staining, 1 x $10^6$ cells were transferred per well of a LoBind Microplate 96/V-PP 96-well plate (Eppendorf, Cat. No. 0030603303), and centrifuged for 5 minutes at 2000g, removing the supernatant by inversion. Cells were resuspended in a primary antibody mix, with a base of PBS + 0.5% BSA and 0.1% Triton-X100 (VWR Chemicals, Cat. No. 28817.295): Sox2-V450 (1:100; clone O30-678; BD Biosciences, Cat. No. 561610), Pax6-PerCPCy5.5 (1:100; clone 018-1330; BD Biosciences, Cat. No. 562388), Nkx6.1-AlexaFluor647 (1:100; clone R11-560; BD Biosciences, Cat. No. 563338), Goat Olig2 unconjugated (1:400; R&D, Cat. No. AF2418), Nkx2.2-PE (1:100; clone 74.5A5; BD Biosciences, Cat. No. 564730). The samples were incubated for 16 hours at 4℃, followed by two rounds of centrifugation, inversion and resuspension in PBS + 0.5% BSA and 0.1% Triton-X100. Cells were then resuspended in a secondary antibody mix with the same base: Donkey anti-goat AF488 (1:1000; Invitrogen, Cat. No. A11055). This was incubated for 40 minutes at room temperature, then washed twice as above. Cells were resuspended in 200$\mu$L of PBS + 0.5% BSA and run on the BD LSRFortessa Cell Analyzer, analysing 150$\mu$L of each sample. Single stain controls were additionally run, using UltraComp eBeads Plus Compensation Beads (Invitrogen, 01-3333-41) stained with the above conjugated antibodies.

##### 2.3 Flow cytometry analysis

For all flow-cytometry analysis, live and fixed, FCS files were analysed using FlowJo (BD Biosciences), gating in the following order to obtain live singlets: FSC-A vs SSC-A; SSC-A vs SSC-H; FSC-A vs FSC-H; FSC-A vs LIVE/DEAD Near-IR. For dual-colour live flow cytometry analysis for ZsGreen reporter experiments, mScarlet3 positive cells were further subset, then mean fluorescence intensities in the ZsGreen channel were calculated. For fixed intracellular flow cytometry analysis, live singlets were further gated on Sox2 positivity to subset on neural progenitors.

#### 3 Immunofluorescence (IF) staining

##### 3.1 IF for 2D differentiations

Samples were pre-incubated with PBS + 0.5% BSA and 0.1% Triton-X100 (VWR Chemicals, Cat. No. 28817.295). Generically, primary antibody mixes were generated with the same base, added to wells at 150$\mu$L per 24-well well and incubated for 16 hours at 4℃. Wells were washed three times for 10 minutes each with PBS + 0.5% BSA and 0.1% Triton-X100, then supplied with the secondary antibody mix with the same base, and incubated at room temperature for 1 hour. Finally, wells were washed three times as above, then supplemented with PBS. For the single-colony re-plating experiment, we used the following primary (anti-Sox2 (1:1000, Invitrogen, Cat. No. 14-9811-80), goat anti-Olig2 (1:1000, R&D, Cat. No. AF2418), and mouse anti-Pax7 (1:500, DSHB)) and secondary (CS405 donkey anti-rat (Biotium, Cat. No. 20419); AF647 donkey anti-goat (Invitrogen, Cat. No. A21447); AF568 donkey anti-mouse (Invitrogen, Cat. No. A10037); all 1:1000) antibody mixes. 2D replated single-colonies were imaged with the an inverted Olympus FV3000 laser-scanning confocal microscope with an Olympus UPlanSApo 10x-0.40NA objective.

##### 3.2 IF and clearing of neural tube organoids

For immunofluorescence staining of 3D neural tube organoids, samples were pre-incubated for 4 days at 4℃ with PBS + 0.5% BSA and 0.1% Triton-X100 (VWR Chemicals, Cat. No. 28817.295). 50$\mu$L of the primary antibody mix (1:800 Olig2 chicken (Aves lab Cat. No OLIG2-0020); 1:400 FoxA2 goat (R&D Cat No. AF2400); 1:2000 Nkx6.1 rabbit (Novus Cat No. NBP1-49672)) was supplied to each well and incubated overnight at 4℃. Samples were washed once for 10 minutes each with PBS + 0.5% BSA and 0.1% Triton-X100 (VWR Chemicals, Cat. No. 28817.295) then incubated at room temperature for 6 hours, followed by a subsequent wash. 50$\mu$L of the Alexa Fluor conjugated secondary antibody mix (donkey anti-goat AF647 (Invitrogen Cat No. A-21447); donkey anti-chick AF568 (Jackson Cat No. 703-575-155); donkey anti-rabbit AF488 (Invitrogen Cat No. A-21206; all 1:1000) and 1:1000 DAPI (Cell Signalling 4083) was then supplied to each well and incubated overnight at 4℃. Samples were then washed three times in PBS + 0.5% BSA and 0.1% Triton-X100 for 10 minutes each. To clear samples, the buffer was aspirated and replaced with 100$\mu$L CUBIC-R+(N) [Matsumoto et al., 2019], incubated for 15 minutes, replaced with fresh CUBIC-R+(N), and repeated once more.

3D neural tube organoids were imaged with Olympus CSU-W1 SoRa Spinning Disk using a 10x air objective (10x UPLXAPO 0.4NA). Per well, a 3x3 tile-scan was performed per Z-stack, acquiring 72 Z-stacks at 3$\mu$m separation, allowing for the acquisition of the entire 3D volume of every organoid in each well.

#### 4 NeMECiS screen

##### 4.1 NeMECiS mESC differentiation

In duplicate, $10^7$ mESCs were infected with the NeMECiS lentivirus library in ESGRO at MOI<1, maintained in CellBIND T175 flasks (Corning, Cat. No. 3292). After 48 hours, media was changed into ESGRO supplemented with 1ug/mL Puromycin Dihydrochloride (Gibco, Cat. No. 12122530). Cells were then expanded for one passage in ESGRO and Puromycin Dihydrochloride. 150mm tissue culture plates were coated with 1:100 Vitronectin (Gibco, Cat. No. A31804) diluted in DMEM-F12 (-/-) (Gibco, Cat. No. 21331-020) for 1 hour. Per replicate, a total of $6 \times 10^7$ cells were plated in N2B27 supplemented with bFGF as above. After

48 hours, media was changed into N2B27 supplemented with bFGF and CHIR99021 as above. After a further 20 hours, cells were dissociated with StemPro Accutase (Gibco, Cat. No. A1110501), centrifuged, and re-plated at the same density in N2B27 supplemented with 100nM RA (Sigma, Cat. No. R2625, additionally supplementing the media for half of the cells with 500nM SAG (Merck, Cat. No. 566660-5mg), onto 100mm tissue-culture plates coated with 1:100 GelTrex (Gibco, Cat. No. A1413202) diluted in DMEM-F12 (Gibco, Cat. No. 21331046). Media was changed daily. After 72 hours, cells were dissociated with StemPro Accutase, centrifuged and resuspended in 0.5% BSA (Sigma-Aldrich, Cat. No. A7979-50ML) in PBS (Gibco, Cat. No. 14190-094). 30% of each sample was immediately pelleted and frozen, and the remaining cells were flow sorted, as described below.

#### 4.2 NeMECiS flow sorting specifications

24 hours before flow sorting, 10mL polypropylene tubes were incubated with 100% Fetal Bovine Serum. Prior to sorting, samples were supplemented with 10 ng/ml DAPI (Invitrogen, Cat. No. D1306). Samples were run on MoFlo XDP and BD InFlux. Sorted events were gated on live singlets (FSC-A vs SSC-A; SSC-A vs SSC-H; FSC-A vs DAPI). Four bins of ZsGreen intensity were established for each sample (two replicates, each for 0nM and 500nM SAG), demarcating the following percentile bins, following an optimal binning strategy found in [de Boer et al., 2020]: $[0-15], [28.333-43.333], [56.666-71.666], [85-100]$. FBS in polypropylene tubes was decanted and samples were sorted according to the above gating strategy. To assess sort quality and calibrate downstream analysis, 10$\mu$L of each sorted sample, as well as the unsorted pool, were extracted and run on the BD LSRFortessa Cell Analyzer, as described above. The final number of sorted cells is reported in Fig. S2E.

#### 4.3 NeMECiS library preparation and sequencing

To extract genomic DNA, sorted cells for the NeMECiS screen were pelleted in 1.5mL DNA LoBind microfuge tubes (Eppendorf, Cat. No. Z666548) by centrifugation for 10 minutes at 500g. Genomic DNA was extracted for sorted and unsorted pellets using a Quick-DNA Miniprep Plus Kit according to the manufacturer's instructions (Zymo Research, Cat. No. D4069), dividing each sorted sample into six columns each. Each column was eluted in 75$\mu$L of the supplied elution buffer (10 mM Tris-HCl, pH 8.5, 0.1 mM EDTA) pre-warmed to 60℃, into 1.5mL DNA LoBind microfuge tubes, pooling the samples of the same origin to bring the total eluted gDNA to 450uL. Integrated barcode concatemers were amplified by PCR using NEBNext High-Fidelity 2X PCR Master Mix (New England Biolabs, Cat. No M0541L) by combining on ice: 450$\mu$L gDNA, 500$\mu$L 2X master mix, 5$\mu$L each of 100$\mu$M Nextera-adaptor primers (indexing each sample, see below), and 40$\mu$L nuclease-free water. Amplification was carried out under the following program: 5 minutes at 72℃, 30 seconds at 98℃, followed by 17 cycles of 30 seconds at 98℃, 30 seconds at 63 ℃ and 60 seconds at 72℃, and subsequently 5 minutes at 72℃. At this step, all PCR products were pooled. The pooled amplicon DNA was then cleaned up using 1.8x v/v of AMPureXP beads (Beckman Coulter, Cat. No. A63882) and eluted in 50$\mu$L EB buffer (10 mM Tris-Cl, pH 8.5). Library concentration was determined using the KAPA Library Quanitification Complete Kit Universal (Roche, Cat. No. 07960140001) according to manufacturer's instructions, using the QuantStudio 12K Flex Real-Time PCR System for qPCR (Applied Biosystems, Cat. No. 4471087). 50% of the amplified sequence was further amplified using NEBNext High-Fidelity 2X PCR Master Mix and P5 (AATGATACGGCGACCACCGA) and P7 primers (CAAGCAGAAGACGGCATACGAGAT) using the same programme as above, then re-quantified as above to establish a concentration in excess of 1nM. Size spectra were examined with a Bioanalyzer using the DNA High Sensitivity DNA Kit (Agilent, Part No. 5067-4626) to confirm library quality. The library was then sequenced using the Illumina NextSeq 500 Sequencer on a 'High Output' run using the NextSeq 500/550 High Output Kit v2.5 (300 Cycles) (Illumina, Cat. No. 20024908) according to the manufacturer's instructions. Sequence demultiplexing was performed with the built-in software by supplying the corresponding indexing sequences for each sample.

### 5 Molecular cloning

#### 5.1 Cloning lentivirus backbones

We based all lentivirus backbones on T3G-zsGreen-ultramiR-SFFV-rtTA (L3zUSR) [Delas et al., 2019]. Specifically, we generated a Golden Gate compatible lentivirus base vector – pLV-GG-WPRE – by cloning a bacterially expressible superfolder GFP flanked by Esp3I cut sites (derived from Addgene: 123943; [Fonseca et al., 2019]) between the CTS and WPRE sequences, replacing the existing contents of the plasmid with a Golden Gate landing-pad sequence (overhangs: 5'CTGA–AGCA 3'). This construct produces bacterial colonies that are visibly green under a blue-light box, whereas successful subsequent Golden Gate cloning with this vector produces non-green colonies.

We constructed reporter constructs through two alternative cloning strategies. We either inserted CREs into a backbone containing a minimal promoter, fluorescent protein, and one or more selectable marker; or we first inserted the CRE (or sequentially blocks thereof) into an 'empty' backbone, introducing the reporter and selectable cassettes in a second step (see synCRE construction).

For the first strategy, we built a series of base reporter vectors. We exclusively use the Sonic Hedgehog minimal promoter derived in [Kvon et al., 2020] (Addgene: 139098), optimised for high signal-to-noise in an in vivo mouse reporter context, to drive the expression of our reporter fluorescent protein ZsGreen, terminated by the SV40 polyA sequence. Upstream of the minimal promoter, we insert the same bacterially expressible superfolder GFP as above, this time flanked by PaqCI cut sites (overhangs: 5'CTGA–AGCA 3'), to act as a landing-pad for subsequent cloning. Downstream of the reporter cassette, we introduced one of two selectable cassettes, driven by an Ef1a promoter, and flanked by the 3' WPRE of the backbone: (1) Puromycin resistance gene (PuroR); (2) Puro-t2A-mScarlet3 (Addgene: 189753; [Gadella Jr et al., 2023]). These backbones are termed pLV-GG-CRErep-Puro and pLV-GG-CRErep-Puro2AmScar3 respectively. These backbones can be used in cloning reactions to insert any CRE with corresponding PaqCI golden-gate adaptors, where successful clonings can be screened by a lack of green fluorescence in bacterial colonies.

#### 5.2 Routine cloning, bacterial culture, plasmid DNA extraction and validation

All cloning was performed using Golden Gate [Engler et al., 2008], using T4 Ligase (New England Biolabs, Cat. No. M0202L) and either Esp3I (Thermo Scientific, Cat. No. ER0451) or PaqCI (New England Biolabs, Cat. No. R0745L) as Type IIS restriction enzymes. In every case, 10$\mu$L reactions were prepared using 25fmol of each part, 1$\mu$L T4 Ligase Buffer, 0.1$\mu$L T4 Ligase, and 0.5$\mu$L of the Type IIS restriction enzyme ($\pm$ 'PaqCI Activator' in the case of PaqCI), thermocycling the reaction with the following protocol: for 25 cycles, 37℃ for 2 minutes, 16℃ for 5 minutes; then 37℃ for 30 mins; then 50℃ for 30 mins; then 80℃ for 10 minutes.

Besides the pooled sequential cloning used to build the NeMECiS plasmid library, all bacterial transformations were performed using MAX Efficiency Stbl2 Competent Cells (Invitrogen, Cat. No. 10268019), according to manufacturer's instructions. All bacteria were grown at 37℃ overnight in L-Broth, supplemented with the appropriate antibiotics (Carbenicillin (for Ampicillin resistance) at 100$\mu$g/mL (Merck Cat. No C1389-5G), Kanamycin at 50$\mu$g/mL (Sigma Aldrich Cat No. 60615), and Tetracycline at 15$\mu$g/mL (Sigma Aldrich Cat No. 87128). Minipreps were grown in 800$\mu$L L-Broth, using the fast protocol in the Zyppy Plasmid Miniprep Kit for extraction (Zymo Research, Cat. No. D4037). For larger DNA preps, midipreps were grown in 30mL of L-Broth, extracted using the QIAGEN Plasmid Plus Midi-Prep kit (Qiagen, Cat. No. 12945). In both cases, plasmids were eluted in nuclease-free water. All plasmids used in the study were validated with whole-plasmid Nanopore sequencing, provided by Full Circle Labs Ltd (UK).

#### 5.3 FL-synCRE reporter constructs

Based on our design-space survey of Olig2 CRE fragment re-orderings, we built sets of FL-synCREs that were predicted to have either boosted or diminished activity. Specifically, we synthesised these 1400bp FL-synCREs (without the 4bp spacer sequences inherent to the synCREs studied with NeMECiS) with appropriate Golden Gate adaptor sequences as Integrated DNA Technologies gBlocks, and cloned them into pLV-GG-CRErep-Puro2AmScar3.

#### 5.4 FL-synCRE-Olig2 Piggybac plasmids

We established a base vector for experiments in which the transcription factor Olig2 is placed under the control of FL-synCREs by assembling using Esp3I-based Golden Gate: (1) a Piggybac backbone derived from [Benzinger and Briscoe, 2024]; (2) an expression cassette comprising the Sonic Hedgehog minimal promoter , 3×FLAG-FKBP12$^{F36V}$-GSGSGS-Olig2 (mouse CDS, synthesised as a gBlock), and an SV40 polyA; (3) upstream placing, a superfolder GFP, PaqCI-flanked Golden-Gate landing pad; (4) downstream placing an [Ef1a]-[Blasticidin]-[SV40 polyA] resistance cassette:

`[Landing Pad][Shh mP][FKBP-Olig2][polyA][Ef1a][BSD][polyA]`

FL-synCREs were cloned into this backbone using the same synthesised sequences as above.

#### 5.5 synCRE construction: pooled and individual

##### 5.5.1 Overview

synCREs are assembled by sequential cloning of CRE fragments into a lentivirus backbone. For trimer synCREs, this is achieved by three sequential cloning reactions, inserting into the backbone a single fragment and a single barcode per step. This sequential nested cloning approach builds up CRE fragments and barcodes one by one from the outside inwards. Once inserted, fragments and barcodes remain physically linked via the plasmid backbone, meaning the composition and order of fragments can be read out (in reverse) by the composition and order of their cognate barcodes. Additionally, at each step, a distinct secondary antibiotic resistance (in addition to the backbone's existing ampicilin resistance) is introduced, allowing successful cloning events to be selected for during bacterial culture. In a final step, the landing pads and associated secondary antibiotic resistances are replaced by the reporter construct.

This is achieved by alternate usage of pairs of Type IIS (Golden Gate) restriction enzymes and secondary bacterial antibiotic resistance genes. Specifically, each part is designated either an 'odd' structure:

`CACCTGCatatACCA[CRE fragment]CTGAtGAGACG[KanR]CGTCTCaAGCA[8bp Barcode]GATTatatGCAGGTG`

or an 'even' structure

`CGTCTCaCTGA[CRE fragment]ACCAatatGCAGGT[TetR]CACCTGCatatGATT[8bp Barcode]AGCAtGAGACG`

with the following key:

- Esp3I recognition sequence
- Esp3I associated 4bp overhang
- PaqCI recognition sequence
- PaqCI associated 4bp overhang
- Spacer sequence
- [Feature]

Here, parts with an 'odd' structure can be inserted into plasmid with a compatible PaqCI ('odd') landing pad. 'Odd' parts – and consequently their cloned products – contain an internal Esp3I ('even') landing pad. In the 'even' structure, the reverse is true, where the part can be inserted into a compatible Esp3I ('even') landing pad, and contains within it a PaqCI landing pad.

Consequently, inserting an 'odd' part into an 'odd' backbone yields the following reaction

`[LV backbone]ACCAatatGCAGGT...CACCTGCatatGATT[LV backbone]`

`+ 'odd' part`

$\Longrightarrow$ `[LVbb]ACCA[Frag]CTGAtGAGACG[KR]CGTCTCaAGCA[BC]GATT[LVbb]`

This has centrally an 'even' landing pad (`CTGAtGAGACG[KR]CGTCTCaAGCA`) into which an 'even' part can be inserted in a subsequent step

$\Longrightarrow$ `[LVbb]ACCA[Frag]CTGA[Frag]ACCAatatGCAGGT[TR]CACCTGCatatGATT[BC]AGCA[BC]GATT[LVbb]`

This recovers the initial structure with an 'odd' landing pad (`ACCAatatGCAGGT[TR]CACCTGCatatGATT`), allowing for repeated cycling, for one more time in our case, but indefinitely in principle.

$\Longrightarrow$ `[LVbb]ACCA[Frag]CTGA[Frag]ACCA[Frag]CTGAtGAGACG[KR]CGTCTCaAGCA[BC]GATT[BC]AGCA[BC]GATT[LVbb]`

(Note here we use the following shorthand notation: Frag = CRE fragment; KR = Kanamycin Resistance; TR = Tetracycline Resistance; BC = 8bp Barcode; LVbb = Lentivirus backbone).

In a final step, the landing pads and associated secondary antibiotic resistances are replaced by the reporter construct, using a *terminal* plasmid, which has the following structure:

CGTCTCtCTGA[ShhmP-ZsGreen-polyA-Ef1a-Puro±t2AmScarlet3][Nextera Read 1]AGCAaGAGACG

where we insert the Nextera Read 1 recognition sequence (TCGTCGGCAGCGTCAGATGTGTATAAGAGACAG)) directly into the plasmid. The terminal plasmid containing Puro alone is termed *pNeMECiS-term-Puro*, and the plasmid additionally containing mScarlet3 is termed *pNeMECiS-term-Puro2AmScar*.

Note that this terminal plasmid is equally compatible for insertion into plasmids with one or three fragments/barcodes, allowing us to build single-fragment controls.

##### 5.5.2 Preparation of synCRE backbone and assembly of the barcode concatemer

To perform the above nested sequential cloning, we established an 'odd' compatible backbone, containing the necessary parts for barcode extraction later. Specifically, using *pLV-GG-WPRE* as a base vector, we cloned in the following insert

ACCAatatGCAGGTG[lacZ cassette]CACCTGCatatGATT[min-pA][Nextera Read 2 RC]

Here, Nextera Read 2 RC is the reverse complement of the Nextera Read 2 recognition sequence (RC = CTGTCTCTTATACACATCTCCGAGCCCACGAGAC), and min-pA is a 56bp truncated variant of the beta-globin polyadenylation signal (GATTAATAAAGGAAATTTATTTCATTGCAATAGTGTGTTGGAATTTGTGTCTCTCA). We term this backbone *pLV-eNextpA-WPRE*. Note that this plasmid contains an 'odd' landing pad (ACCAatatGCAGGTG[...]CACCTGCatatGATT).

Upon sequential nested cloning, and introduction of the terminal sequence, one yields the following 163bp sequence built into the plasmid:

[Nextera Read 1]AGCA[BC]GATT[BC]AGCA[BC]GATT[min-pA][Nextera Read 2]

This can be directly amplified from the plasmid, or upon lentivirally mediated integration, using Nextera-compatible primers with structures

5'-AATGATACGGCGACCACCGAGATCTACAC[seqBC]TCGTCGGCAGCGTCAGATGTG-3'

5'-CAAGCAGAAGACGGCATACGAGAT[seqBC]GTCTCGTGGGCTCGGAGATGT-3'

Giving a total Illumina sequencing compatible amplicon with the following structure

5'-[P5][seqBC][Nextera Rd1]AGCA[BC]GATT[BC]AGCA[BC]GATT[min-pA][Nextera Rd2][seqBC][P7]-3'

##### 5.5.3 Allocation of CRE fragments

200bp CRE fragments were derived from validated CREs for three well-established Sonic Hedgehog dependent TFs: Olig2, Nkx22, Pax6 [Oosterveen et al., 2012, Peterson et al., 2012]. Their full sequences are provided in Table 1. Note here that we use a combination of the sequences for the Olig2 CRE defined in [Oosterveen et al., 2012] and [Peterson et al., 2012], which is 1305bp in length.

We generated a 200bp fragmentation starting from the 5' end of each of the above sequences, with the final fragment overlapping and extending beyond the natural sequence's *3'* extent. We then generated a 200bp fragmentation with a 100bp 3' offset. Sequences are provided in Table 2

##### 5.5.4 Preparation of 'even' and 'odd' subcloning vectors

Nested cloning proceeded by sequentially inserting 'odd' parts into 'odd' backbones to generate 'even' plasmids, followed by the insertion of 'even' parts to return the plasmid to 'odd'. Practically, this was achieved by Golden Gate reactions between two types of plasmid: a lentivirus backbone and a (pool of) subcloning vectors. Each subcloning vector has an overall structure:

...[Ori][Esp3I/PaqCI][Frag][PaqCI/Esp3I][TetR/KanR][PaqCI/Esp3I][BC][Esp3I/PaqCI]...

These plasmids were cloned as follows. Integrated DNA Technologies eBlocks were synthesised the following structure, allocating the connection between 8bp barcodes ([BC]) and 200bp CRE fragments ([Frag]):

'Odd':

attaCACCTGCatatACCA[Frag]CTGAatatGCAGGTG[Spacer Odd]CACCTGCatatAGCA[BC]GATTatatGCAGGTGttaa

'Even':

attaCGTCTCtCTGA[Frag]ACCAaGAGACG[Spacer Even]CGTCTCtGATT[BC]AGCAaGAGACGttaa

Where [Spacer Odd] = CGTCGGGCTCGACATCGGCAAGGT

and [Spacer Even]=CGTCGGGCTCGACATCGGCAAGGTCGGACTGAATTCATGC.

NB: these spacer sequences act as fillers and are removed in the assembly of the subcloning vectors.

Next, a high-copy-number ColE1/pMB1/pBR322/pUC origin of replication ([Ori]) was amplified by PCR from Addgene: 123943 [Fonseca et al., 2019], adding on 'odd' or 'even' adaptors, introducing the outer Type IIS restriction sites into the plasmid that are subsequently used for sequential, nested cloning (note the inverted architecture of recognition cut-site pairs at the amplicon termini):

Odd: 5'-ttttGATTatatGCAGGTG[Ori]CACCTGCatatACCAaaaa

Even: 5'-ttttAGCAtGAGACG[Ori]CGTCTCaCTGAaaaa-3'

Cognate resistance cassettes were then amplified by PCR: [KanR] (Kanamycin resistance gene, amplified from Addgene: 123943 [Fonseca et al., 2019]) for 'odd' subcloning vectors; and [TetR] (Tetracycline resistance gene, amplified from Addgene: 92394 [Williams et al., 2018] for 'even' subcloning vectors.

Even: 5'-ttttCGTCTCtACCAatatGCAGGTG[TetR]CACCTGCatatGATTaGAGACGaaaa-3'

Odd: 5'-ttttCACCTGCatatCTGAtGAGACG[KanR]CGTCTCaAGCAatatGCAGGTGaaaa -3'

where the multicoloured sequences NNNN denote dual potential usage by Esp3I and PaqCI. PCRs were performed using Phusion Flash High-Fidelity PCR Master Mix (Thermo Scientific, Cat No. F548S) by manufacturer's instructions, and purified using the QIAquick PCR Purification Kit (QIAGEN, Cat No. 28104).

PCRs and eBlocks were then normalised to 25fmol/$\mu$L and cloned to generate plasmids using Golden Gate ('odd' with PaqCI; and 'even' with Esp3I), using a modified reaction protocol (10 cycles at 37℃ for 2 minutes, 16℃ for 5 minutes; 30 minutes at 16℃), mixed at a 1:1 ratio with a T4 ligase mix (per 10$\mu$L: 0.5$\mu$L T4 ligase, 1$\mu$L T4 ligase buffer, New England Biolabs, Cat No. M0202L), and incubated for 16 hours at 4℃. Cloning products were then transformed in NEB-5alpha Competent E. coli (New England Biolabs, Cat No. C2987H), miniprepped (Zyppy Plasmid Miniprep Kit; Zymo Research, Cat No. D4037), verified by Sanger Sequencing, and midiprepped (QIAGEN Plasmid Plus Midi-Prep kit, Cat. No. 12945).

Additionally, 1mL live cultures (L-Broth) for each set of 'odd'/'even' subcloning vectors (25 each) were serially cultured twice in 96 deep well plates (Axygen, Cat No. P-2ML-SQ-C-S), pooled in an equivolumetric ratio and midiprepped (QIAGEN Plasmid Plus Midi-Prep kit, Cat. No. 12945) together to generate equimolar pools.

##### 5.5.5 Pooled sequential nested cloning

Firstly, pLV-eNextpA-WPRE (containing an 'odd' landing pad) and the *'odd'* subcloning plasmid pool were individually secondarily cleaned up using DNA-Concentrator 5 (Zymo Research, Cat No. D4014) according to manufacturer's instructions, eluting in nuclease-free water and normalising (using Qubit quantification of DNA concentration using the broad-range kit, Invitrogen, Cat No. Q32850) to 25fmol/$\mu$L. 24$\times$ 10$\mu$L Golden Gate reactions with PaqCI were run (50 cycles), and immediately cleaned up using DNA-Concentrator 5, eluting in 50$\mu$L water. 2$\mu$L of this elution was transformed with Endura Electrocompetent Cells (Biosearch Technologies, Cat No. 60242-1) according to manufacturer's instructions, then cultured directly at 28℃ (170rpm) in 1L L-Broth supplemented with Carbenicillin and Kanamycin until the $OD_{600}$ reached 1.0 ($\sim 14$ hours). 100mL of the culture was then maxiprepped using the QIAGEN Plasmid Plus Maxi Kit (Cat No. 12965).

Next, an equivalent procedure was performed by cleaning up the *round 1* cloned pool (containing an 'even' landing pad) with the *'even'* subcloning plasmid pool with DNA-Concentrator 5, normalising, and running a Golden Gate reaction with the same specifications, this time with Esp3I. An identical transformation, culture and DNA extraction procedure was then performed, this time using Tetracycline in place of Kanamycin, to establish the *round 2 cloned pool*. After this, the same protocol was performed as in round 1, but using *round 2 cloned pool* in place of *pLV-eNextpA-WPRE*. Finally, the same procedure was performed as in round 2, but combining *round 3 cloned pool* with *pNeMECiS-term-Puro*, this time culturing the transformation with Carbenicillin alone, and performing and pooling $4\times$ maxipreps.

##### 5.5.6 Arrayed preparation of individual synCRE reporter constructs

synCRE reporter constructs were established in an analogous manner to the above, but in an arrayed fashion, allowing for the construction of desired fragment combinations and orderings. In this, clean-up steps before Golden-Gate reaction preparation and transformation were omitted, and MAX Efficiency Stbl2 Competent Cells (Invitrogen, Cat No. 10268019) were used. Here, 1$\mu$L of the reaction was combined with 10$\mu$L competent cells, heat-shocked and recovered according to manufacturer's instructions, and grown at 37℃ for 16 hours in 800$\mu$L L-Broth, split across two wells of 1.1mL 96 deep well plates (Axygen, Cat No. 736-0339), purifying minipreps using the Zyppy Plasmid Miniprep Kit for extraction (Zymo Research,

Cat No. D4037). In the final step, *pNeMECiS-term-Puro2AmScar3* was used to establish dual-colour reporter plasmids. Single fragment reporter constructs were established in an identical manner, cloning in *pNeMECiS-term-Puro2AmScar3* in the first step rather than the third. Matched full-length natural CRE and CRE-less, minimal-promoter only (spacer sequence = `CGAGCGTCTCT`) reporters were cloned in a similar manner.

##### 5.5.7 Barcode allocation

NeMECiS relies on sequencing barcode concatemers to retrieve compositional information. Consequently, it is crucial that the barcodes used are maximally informative, given potential misassignment. We built a Markov chain Monte Carlo Metropolis Hastings algorithm to scan sequence space for optimal combinations of 25 barcodes. The algorithm is summarised below:

1. Generate 25 random sequences of 8bp. Add the overhangs for cloning to each side. Re-generate if any of the random sequences has: (i) >2 repeated bases; or (ii) a GC content below 40% or above 60%.
2. Calculate the sum $H$ of Hamming distances among each pair of the 25 sequences (=300 pairs).
3. Randomly select one of the 25 sequences and replace the central 8bp with another random sequence that follows the two conditions above.
4. Calculate $H_{new}$ of the new set of sequences.
5. If $H-H_{new} < 0$, accept the swap. Else, accept the swap if $p < exp((H_{new} - H)/T)$, where $p$ is a random number between [0,1]. (T=0.01). Otherwise, reject the swap.
6. Repeat 2-5 for another 2989 iterations.
7. Repeat 2-5 for another 10 iterations, logging the sequence combinations.
8. Repeat 1-7 500 times to establish an ensemble of sequence combinations.
9. Calculate E for each logged sequence:
   - $E = H + H_{shift} + H_{delete}$
   - Where $H_{shift}$ is the sum of the minimum of the Hamming distances among pairs of sequences where sequences are shifted 1bp 5' or 3'.
   - And $H_{delete}$ is the sum of the minimum Hamming distances among pairs of sequences where each of the base-pairs of one of the sequences is deleted.
10. Determine the combination of 25 barcodes with the maximum E.

The resulting barcodes are given in Table 3

#### 5.6 Gli1/2 binding site mutant reporter

Using the motif enrichment analysis described below, we identified a putative Gli1/2 binding site in a strongly signal responsive synCRE f20-f11-f16:

`...GTGAAATCTTGGGTGGTATGGGCACCG...`

`Key: surrounding sequence, expanded motif,MOODS/JASPAR2024 called motif`

Adding 3bp to each side of the called motif to ensure complete disruption, we computationally scrambled the base-pairs to generate the sequence:

`...GTGAAGGTTGAGCGTGGGTATCACCG...`

We then synthesised both the WT and Gli-mutated variants of f20-f11-f16 as Integrated DNA Technologies gBlocks, and cloned them into *pLV-GG-CRErep-Puro2AmScar3*.

### 6 NeMECiS sequencing processing

#### 6.1 Alignment and barcode counts

A white-list of all 15,625 possible barcode concatemers was generated using a custom script. These were appended with left and right adaptor sequences to establish `amplicons.fa`, containing sequences each of length 163bp, labelled with the corresponding composition and ordering of CRE fragments. These have a generic form:

```
TCGTCGGCAGCGTCAGATGTGTATAAGAGACAGAGCANNNNNNNNGATTNNNNNNNNAGCANNNNNNNNGATT
GATTAATAAAGGAAATTTATTTCATTGCAATAGTGTGTTGGAATTTGTGTCTCTCACTGTCTCTTATACACATCTCCGAGCCCACGAGAC
Key:  Nextera Rd1 Golden Gate overhang 8bp-BC sequence polyA terminator sequence Nextera
Rd2
```

Bowtie 2 was used for barcode calling. Index files were generated using the command:

```
bowtie2-build amplicons.fa amplicons
```

Global alignment of barcode concatemers for each *fastq* file from the NextSeq 500 demultiplexed output was achieved by running the following command, trimming alignment to exclude Nextera adaptors:

```
bowtie2 --threads 32 --trim-to 3:96 -x amplicons -1 fastq_file_read1.fastq -2 fastq_file_read2.fastq
```

From the alignments recorded in the *.sam* file output, a custom python script `process_sam_files.py` was used to generate a count-matrix for each barcode in each sample. Here, reads by barcode were counted that had a perfect match (CIGAR output of `96M`), and successfully aligned to a member of the white-list.

#### 6.2 Bayesian inference of fluorescence from barcode distributions

The output of NeMECiS is the frequencies of each of the 15,625 synCRE barcodes in the entire population (background distribution) and the frequencies of each barcode after flow-sorting. We converted these frequencies into estimated fluorescence intensities, with associated uncertainties in estimates, by a Bayesian inference approach. This allows for quantitative comparison and batch-normalisation across conditions, by placing all values on a common scale, and further aggregates the multiple measurements per barcode into a single summary statistic.

##### 6.2.1 Outline of approach

The basic principle of the approach is that reporter fluorescence for each synCRE, and corresponding barcode $i$, will follow a distribution in intensity ($y$) as measured by flow cytometry ($P(y|i)$). We assume that this is a log-normal distribution for simplicity, or rather is normally distributed in log transformed intensities ($x = \log y; P(x|i) \sim \mathcal{N}(\mu_i, \sigma_i^2)$). Our task is to learn the parameters $\{\mu_i, \sigma_i\}$ given the measured data.

Measurements are performed in pool. This means that all fluorescence distributions measured during quality-control flow cytometry are a mixture of the barcode-specific distributions, weighted by their relative frequencies in the pool $\pi_i$. Therefore, the global distribution of (log) fluorescence in the pool (for a specific condition/replicate) is given by a Gaussian Mixture Model.

$$P(x) = \sum_i \pi_i \mathcal{N}(\mu_i, \sigma_i^2) \tag{1}$$

To deconvolve the barcode-specific components of this global distribution, we can compare the frequencies of barcodes in the unsorted pool to the frequencies in each of the sorted bins $b_j \in \{1, 2, 3, 4\}$. The flow sorter is assumed to transfer cells into bin $b_j$ if the log ZsGreen fluorescence level lies between the limits of the sort gate $[b_j^L, b_j^U]$. This assumes that sorting is 'perfect' i.e. that the sort bin only contains cells with log fluorescence values within these limits. Thus the probability of sorting into gate $b_j$ given a particular log fluorescence $x$ is given by:

$$P(b_j|x) = \frac{1}{b_j^U - b_j^L} \text{ for } b_j^L \le x \le b_j^U \tag{2}$$

Flow-sorting is performed in a manner that is indiscriminate to synCREs (once conditioned on fluorescence) i.e. $P(b_j|x, i) = P(b_j|x)$. Consequently, we can calculate the probability of being sorted into a bin $b_j$ for a particular synCRE $i$ through marginalisation to obtain $P(b_j|i)$,:

$$P(b_j|i) = \int_{-\infty}^{\infty} P(b_j|x)P(x|i)dx \tag{3}$$

$$= \phi(\frac{b_j^U - \mu_i}{\sigma_i}) - \phi(\frac{b_j^U - \mu_i}{\sigma_i}) \tag{4}$$

where $\phi(z)$ is the cumulative distribution function of the normal distribution with parameters $\mathcal{N}(0, 1)$. To then calculate the proportion of a synCRE $i$ in a bin $b_j$, we apply Bayes's rule:

$$P(i|b_j) = \frac{P(i)P(b_j|i)}{P(b_j)} \tag{5}$$

Note that $P(b_j)$ represents the probability of cells displaying fluorescence values between the bin-limits of $b_j$ across the entire unsorted pool. Consequently, $P(b_j)$ can be determined empirically from the flow-cytometry distribution of log ZsGreen fluorescences in the unsorted pool $P(x)$

$$P(b_j) = \int_{-\infty}^{\infty} P(b_j|x)P(x)dx \tag{6}$$

$$= \int_{b_j^L}^{b_j^U} P(x)dx \tag{7}$$

Therefore, defining $\pi_i$ as the density of each synCRE in the unsorted pool, we have:

$$P(i|b_j) = \frac{\pi_i[\phi(\frac{b_j^U - \mu_i}{\sigma_i}) - \phi(\frac{b_j^U - \mu_i}{\sigma_i})]}{\int_{b_j^L}^{b_j^U} P(x)dx} \tag{8}$$

##### 6.2.2 Finite number effects

The above describes the probability a given cell is sorted into a particular bin, given it contains a particular barcode. As we sorted a finite, while large, number of cells, we additionally consider and integrate the implications of these finite number effects on the confidence of our measurements.

There are two additional potential sources of noise: (a) the relative frequencies of cells containing each barcode found in cells actually sorted in each bin; (b) the relative frequencies of sequencing reads given the frequencies of the barcodes in each bin. Both, strictly speaking, follow multinomial distributions, containing the frequencies/probabilities for each of the 15,625 barcodes. In practice, such scenarios are well approximated by binomial distributions (given the covariance among any pair of different barcodes in this multinomial distribution will be vanishingly small).

$P(i|b_j)$ is the probability that a cell contains barcode $i$ given it is allocated to bin $b_j$. If $N_{sort,j}$ cells are sorted into bin $b_j$, $N_{cell}(i|b_j)$ is the number of those cells that contain barcode $i$. Specifically, this follows: $N_{cell}(i|b_j) \sim Binom(N_{sort,j}, P(i|b_j))$. Correspondingly, the probability that a cell contains barcode $i$ among a finite number of sorted cells is given by: $P(i|b_j, SORTED) = N_{cell}(i|b_j)/N_{sort,j}$. For computational tractability, this distribution is approximated by a normal distribution

$$P(i|b_j, SORTED) \sim \mathcal{N}\left(P(i|b_j), \frac{P(i|b_j)(1 - P(i|b_j))}{N_{sort,j}}\right) \tag{9}$$

From this, the absolute read counts $C$ by barcode is:

$$C(i|b_j) \sim Binom(N_{read,j}, P(i|b_j, SORTED)) \tag{10}$$

For the unsorted pool, one can likewise write:

$$C_{pool}(i) \sim Binom(N_{read}^{pool}, \pi_i) \tag{11}$$

##### 6.2.3 Calibrating bin gates

To achieve the scale of the flow-sort required for this screen, each sample was run on independent flow sorting machines. Consequently, we have to calibrate the bin gates among conditions to ensure that all estimated fluorescence values are on a common scale and without systematic errors. This was done using the small fraction of sorted/unsorted cells we kept out for quality control, run on a common flow cytometer.

Firstly, the raw fluorescence distributions of cells (gating on single, live cells) were log transformed and independently fit to Gaussian Mixture Models ($n_{component} = 3$ using the *sklearn* package), providing a continuous approximation of the empirical fluorescence probability distributions. Let us call these $P_{emp}(x|b_j)$ for each gate and $P_{emp}(x)$ for the global, unsorted pool. We then fit the following function to each $P_{emp}(x|b_j)$ to obtain the bin specifications:

$$P_{fit}(x|b_j) = P_{emp}(x) \cdot \frac{\alpha_j}{2} \left[\tanh(m_j(x - b_j^L)) + \tanh(m_j(b_j^U - x))\right] \tag{12}$$

which describes the convolution between the global fluorescence distribution of the pool and a step-like function that goes between 0 and $\alpha_i$, smoothed by the parameter $m_j$, which accounts for experimental error in the measurement. Parameters were fit using *scipy.optimize*'s implementation of the Newton-CG minimisation method, using *JAXlib*'s autodifferentiation capacities to calculate Jacobians and Hessians numerically. We adopted the following cost function on which to perform this optimisation with respect to the free parameters $\alpha_j, b_j^L, b_j^U$:

$$\begin{aligned}\mathcal{E} = &\sum_r \sum_j \left[\int_{-\infty}^{\infty} (P_{fit}(x|b_j, r) - P_{emp}(x|b_j, r))^2\right] \\ &+ \beta \sum_r \sum_j \left[(\tilde{b}_{j,r}^U - b_{j,r}^U)^2 + (\tilde{b}_{j,r}^L - b_{j,r}^L)^2\right] \\ &+ \gamma \sum_i \left[(\Delta_r \hat{\mu}_i)^2 \left(\sum_j C_{i,j,r}/4\right)\right]\end{aligned} \tag{13}$$

where $r$ defines the replicate number $r \in \{1, 2\}$. Here, the latter two sets of terms are used to guide fit values towards the theoretical values. The first term, scaled by the hyperparameter $\beta$, compares the fit upper and lower bin limits $b_{j,r}^U, b_{j,r}^L$ to the values expected from the global distribution if they were placed at exactly their desired percentile positions (see experimental methods). For example, for $b_j = 2$, $\int_{\tilde{b}_{j,r}^L}^{\tilde{b}_{j,r}^U} P_{emp}(x|r) = 0.15$ and $\int_0^{\tilde{b}_{j,r}^L} P_{emp}(x|r) = 0.15 + 0.1333 = 0.2833$. The second term, scaled by the hyperparameter $\gamma$ compares the approximate mean fluorescence levels of each of a set of barcodes $\mu_{i,r}$ across replicates, weighted by their frequency in the data. This can be viewed as guiding batch correction. Specifically:

$$\hat{\mu}_{i,r} = \frac{\sum_j G_{j,r} C_{i,j,r}}{\sum_j C_{i,j,r}} \tag{14}$$

where $C_{i,j,r}$ is the read-count matrix by barcode, gate and replicate, and $G_{j,r}$ is the mean empirical log fluorescence level for a given gate $b_j$ and replicate $r$. The value $\Delta_r \hat{\mu}_i$ is then calculated as $\Delta_r \hat{\mu}_i = \hat{\mu}_{i,1} - \hat{\mu}_{i,2}$. Together this defined the the inferred bin-positions in subsequent calculations (Fig. S2F shaded regions).

##### 6.2.4 Implementation of Bayesian inference

Provided with the read-count matrix $C_{i,j,r,s}$ across barcodes, bins, replicates, and SAG concentrations, we used *pySTAN* [Stan Development Team, 2024] to infer $\{\mu_{i,r,s}, \sigma_{i,r,s}, \pi_{i,r,s}\}$. This is done in a per-barcode/replicate/SAG concentration fashion, given the binomial approximation to the multinomial problem described above, allowing for parallel implementation.

We specify the following priors for these fit parameters:

$$\mu_{i,r,s} \sim \mathcal{N}(6,1) \tag{15}$$

$$\sigma_{i,r,s} \sim \psi inv\text{-}\chi^2(1, 0.1) \tag{16}$$

$$\pi_{i,r,s} \sim Beta(1,1) \tag{17}$$

(where scaled inverse $\chi^2$ distribution is denoted $\psi inv\text{-}\chi^2$)
The *Stan* script used in the fitting, takes the following measurements as input:

- Read-counts by gate for a particular barcode, replicate and SAG concentration ($C_{i,j,r,s}$)
- Number of cells sorted by gate ($N_{sort,j,r,s}$)
- Number of reads by gate ($N_{read,j,r,s}$)
- Number of reads for the unsorted, pooled population ($N^{pool}_{read,r,s}$)
- Upper and lower bounds of each gate, as inferred from the above
- Normalisation constant in the calculation of $P(i|b_j)$, i.e. $P(b_{j,r,s}) = \int_{b^L_{j,r,s}}^{b^U_{j,r,s}} P_{emp}(x|r,s)$. This is calculated empirically once the gate boundaries are inferred using the above.

Using the approach described above, we model using binomial functions the number of reads expected in the pool and in each of the bins, and use the No-U-Turn-Sampler (NUTS) sampler to generate posterior distributions and maximum-likelihood estimates with the following specifications (*n_chains*=8, *iter_warmup*=500, *iter_sampling*=500). We additionally specify the initial condition around which to sample as: $\mu_i \sim \mathcal{N}(\hat{\mu}, 0.5); \sigma_i \sim [0.8, 1.5]; \pi_i = C^{pool}_{i,r,s}(i)/\sum_j C^{pool}_{j,r,s}$. From this, we gain estimates of the expected value of each parameter ($\mathbb{E}(\theta)$), and its associated sample standard deviation ($s(\theta)$).

### 7 Linear modelling framework for learning modular contributions and interactions

#### 7.1 Overview

The overarching strategy of our linear modelling framework is to describe synCRE structure as a set of categorical features, and fit a joint model for CRE activity with and without SAG.

Let the vector $\boldsymbol{y}$ denote the expected activities of each synCRE×SAG combination. Let $\boldsymbol{X}$ be a $N_{observation} \times N_{feature}$ count-matrix of features. Here, $N_{feature} = 2n_{feature}$ where, depending on if the observation was without or with SAG, either the first (approximately) half or second half (respectively) of the entries in a given column are set to zero. From this, we find the optimal parameters of the following ridge-regression loss function:

$$\mathcal{E} = \sum_{i}^{N_{observation}} \left[\hat{w}_i (y_i - \boldsymbol{\beta}^T \boldsymbol{X}_i + c)^2\right] + \lambda \|\boldsymbol{\beta}\|^2 \tag{18}$$

Here, $\beta$ is a $(2n_{feature})$ coefficient vector of regressed values, with the first and second halves corresponding to the coefficients without and with SAG respectively, and $c$ is the intercept. We additionally include a ridge regularisation factor $\lambda$ which performs two functions: (a) to guard against over-fitting; (b) to encourage a parsimonious fit (i.e. involving the fewest number of features) in cases of parametric degeneracy. We additionally include a weighting parameter $\hat{w}_i$ which weights the fit towards observations with lower uncertainty. $\boldsymbol{y}$ and $\hat{\boldsymbol{w}}$ are taken directly from the Bayesian inference of log fluorescence levels, as described above, where $y_i = \mathbb{E}(\mu_i)$ and $w_i = 1/s(\mu_i)^2; \hat{w}_i = (N_{observation} w_i)/\sum_{j=1}^{N_{observation}} w_j$.

#### 7.2 Remark about scale of data

We choose to fit the ridge regression on log-transformed data. As discussed in 12, motivated by thermodynamic models of CRE activity, we anticipate that contributions from each feature (modular contributions and interactions) should multiply to give the overall activity of a synCRE. Consequently, if fluorescence $f = \prod_j f_j$ over $f_j$ feature-decomposed contributions, then $log(f) = \sum_j log(f_j)$, hence allowing for a functional form amenable to linear regression modelling.

#### 7.3 Data filtering

For all models, the following filtering steps were performed on the data. Firstly, we excluded fragment 18 in position 2 as inspection of fragment sequences revealed mutations, and fragment 17 in position 3, due to significantly fewer reads observed for this fragment-position pair. Secondly, we removed all data-points where $s(\mu_i) > 0.1$, deemed to be too large an uncertainty for further analysis. This left 28520 and 28468 data-points for replicate 1 and 2 respectively (out of a total of $25^3 \cdot 2 = 31250$).

#### 7.4 Feature extraction

To extract the features that comprise $\boldsymbol{X}$ in all of the models from the synCRE fragment composition, the following operations are performed. Let $z_i$ be a triple of fragment indices $(z_i = (z_i^1, z_i^2, z_i^3); z \in \{1, 2, ..., 25\}$ in a 5' to 3' direction from furthest to closest to the minimal promoter. For example, for a synCRE $12 - 14 - 16$, $z_i^1 = 12, z_i^2 = 14, z_i^3 = 16$. Let $\boldsymbol{Z}_i$ be the one-hot encoding of $z_i$ ($\boldsymbol{Z}_i = (\boldsymbol{e}_i^1, \boldsymbol{e}_i^2, \boldsymbol{e}_i^3)$, where $\boldsymbol{e}_i^j$ for a given synCRE $i$ and position $j$ is given by: $\boldsymbol{e}_i^j[z_i^j] = 1$ and is $0$ otherwise.

The fragment count matrix $\boldsymbol{X}_F^S$ for a given SAG is given by:

$$\boldsymbol{X}_F^S|_i = \left[ \sum_{j=1}^{3} \boldsymbol{e}_i^j \right] \tag{19}$$

Note that

$$\boldsymbol{X}_F^S|_i \cdot \mathbb{1} \equiv 3 \tag{20}$$

The fragment pair count matrix $\boldsymbol{X}_P^S$ for a given pair of fragments $\{u, v\}$ for a given synCRE $i$ is given by

$$\boldsymbol{X}_P^S|_{i,u,v} = \boldsymbol{e}_i^1[u]\boldsymbol{e}_i^2[v] + \boldsymbol{e}_i^2[u]\boldsymbol{e}_i^3[v] + \boldsymbol{e}_i^3[u]\boldsymbol{e}_i^1[v] \tag{21}$$

Note that, summing over pairs in $\boldsymbol{X}_P^S$ for a given synCRE:

$$\sum_u \sum_{v \geq u} [\boldsymbol{X}_P^S|_{i,u,v}] \equiv 3 \tag{22}$$

The fragment spaced pair count matrix $\boldsymbol{X}_{SP}^S$ for a given pair of fragments $\{u, v\}$ offset by a distance $d$ (e.g. $1 - x - 2 \implies (u = 1, v = 2, d = 2) or (v = 1, u = 2, d = -2)$ where $x$ is a wildcard), is given by the following set of expressions:

$$\boldsymbol{X}_{SP}^S|_{i,u,v,d} = \begin{cases} \boldsymbol{e}_i^3[u]\boldsymbol{e}_i^1[v]; \text{ if } d = -2 \\ \boldsymbol{e}_i^3[u]\boldsymbol{e}_i^2[v] + \boldsymbol{e}_i^2[u]\boldsymbol{e}_i^1[v]; \text{ if } d = -1 \\ \boldsymbol{e}_i^2[u]\boldsymbol{e}_i^3[v] + \boldsymbol{e}_i^1[u]\boldsymbol{e}_i^2[v]; \text{ if } d = 1 \\ \boldsymbol{e}_i^1[u]\boldsymbol{e}_i^3[v]; \text{ if } d = 2 \end{cases} \tag{23}$$

Note that the sum of $\boldsymbol{X}_{SP}^S|_{i,u,v,d}$ over non-degenerate spaced pairs of fragments is always $3$.

$\boldsymbol{X}_P^S$ and $\boldsymbol{X}_{SP}^S$ are constructed such that descriptions of (spaced) pairs is not degenerate. Thus there are $325$ columns in $\boldsymbol{X}_P^S$ and $1250$ columns in $\boldsymbol{X}_{SP}^S$. The overall count matrices are then constructed, for a given feature class $c$, as the concatenation $\boldsymbol{X}_c = \{\boldsymbol{X}_c^-, \boldsymbol{X}_c^+\}$, where the rows are all data-points for a given replicate (across SAG levels).

#### 7.5 Fitting procedure, cross-validation and hyperparameter tuning

Models were fit using the *sklearn.linear_model.Ridge* package, specifying $w_i$ with the *sample_weight* optional field. Models were fit separately (including independent hyperparameter tuning) on each of the two replicates.

To identify the optimal value of the regularisation parameter $\lambda$, we performed hyperparameter scans over the following range $\lambda = 10^q$ ($q$ is evenly sampled between $-2, +2$ inclusive in $32$ regularly spaced steps). Specifically, we identify a $5-$fold split of the data that preserves feature abundance. To do this, we randomly partition $N_{observation}$ into $5$ bins, holding out each of the 5 bins $b$ individually in turn ($\boldsymbol{X}_{\neg b}$), and calculated $\sum_b (\langle \boldsymbol{X}_{\neg b} \rangle_j - \langle \boldsymbol{X} \rangle_j)^2$. Iterating 100 times, we take the minimum of this cost function to find an optimal split.

We then trained the model on each of the $4/5$ subsets, for each specified value of $\lambda$, and calculate the explained variance in each of the withheld bins, and take a mean. This produces a graph with a maximum, identifying an optimal value of $\lambda$ ($\lambda^*$) the balances over and underfitting.

Next, we used $\lambda^*$ to fit the full model, using all $N_{observation}$. For sensitivity analysis, we fit the model on a bootstrapped resampling of the data ($N_{boot} = 50$).

#### 7.6 Model 1: Fragment contributions alone

We start with a model considering fragment contributions alone. Hence here $N_{feature} = 2 \cdot 25 = 50$, corresponding to each of the 25 fragments, with and without SAG. When fit, the values of $\boldsymbol{\beta}$ approximately correspond to the contributions of each fragment with respect to the population average, thus defining modular effects beyond this average.

If we define $\boldsymbol{\beta} = \{\boldsymbol{\beta}^-, \boldsymbol{\beta}^+\}$ as the decomposition of feature coefficients without and with SAG respectively, we can determine the SAG-dependent and SAG-independent effects as $\Delta\boldsymbol{\beta} = \boldsymbol{\beta}^+ - \boldsymbol{\beta}^-$= 'log FC $\Delta$SAG', and $\langle\boldsymbol{\beta}\rangle = \frac{1}{2}(\boldsymbol{\beta}^+ + \boldsymbol{\beta}^-)$, respectively.

##### 7.6.1 Fragment class allocation

Inspection of the distribution of fragment coefficients across $\{\langle\boldsymbol{\beta}\rangle, \Delta\boldsymbol{\beta}\}$ prompted us to score fragments into three categories based on their response to SAG: increased, repressed, neutral. To do this, we used the bootstrapped distributions of the coefficients and calculated $\bar{\beta}_{boot}$ and $s(\beta_{boot})$. We performed a two tailed Z-test, with $z_{crit} = 4.265$ to assign neutrality, then for fragments significantly different from neutrality, we define increasing fragments as those for which $\bar{\beta}_{boot} > 0$. We performed the test across replicates by using $\bar{\beta}_{boot} = \frac{1}{2}(\bar{\beta}^{r=1}_{boot} + \bar{\beta}^{r=2}_{boot})$, and $s(\beta_{boot}) = \sqrt{s(\beta^{r=1}_{boot})^2 + s(\beta^{r=2}_{boot})^2}$. We also use the Z-test to calculate logP-values of fragment contribution.

#### 7.7 Model 2: Fragment contributions + interactions + spacing effects

##### 7.7.1 Scoring interactions

Upon fitting a model with interactions and spacing effects, we first analysed the role of interactions. To do this, we define an interaction parameter $\alpha_{i,j} = \beta_P|_{\{i,j\}} + \sum_{d\in\{-2,2\}} \beta_{SP}|_{\{i,j,d\}} + 2\sum_{d\in\{-1,1\}} \beta_{SP}|_{\{i,j,d\}}$, summing over both position and spacing dependent effects, and weighting the latter with respect to the frequency it appears in the data. From this, bootstrapped means and sample standard deviations could be calculated, and Z-tests could be performed as above.

For the comparison between the multiplicative model (incorporating fragment effects alone) and the interaction model, we calculated the overall SAG-dependent activity of a pair $\Delta y_{i,j} = \Delta\beta_i + \Delta\beta_j + \Delta\alpha_{i,j}$ and compared it to $\Delta y_{i,j} = \Delta\beta_i + \Delta\beta_j$. When $\alpha_{i,j} \neq 0$, points on the graph that plot these models against each other deviate from the diagonal.

##### 7.7.2 Positionality score

We devised a positionality score to rank the importance of spacing on the interactions between fragments. Define $M_{u,v,d,r,q}$ as an array of shape $(25 \times 25 \times 6 \times 2 \times N_{boot})$, holding the values of $\Delta\beta_{SP}$ for a given pair of fragments $\{u, v\}$, at a given spacing $d \in \{-2, -1, -1, 1, 1, 2\}$, for a given replicate $r \in \{1, 2\}$, and a given bootstrap sample $q \in \{1, 2, ..., N_{boot}\}$. Note here that $d = \pm 1$ is duplicated to reflect the doubled frequencies of these features in the data. Let $s_q(M)$ be the standard deviation of the array $M$ along the $q$ dimension, and likewise for $s_d$. We can therefore define $\rho_{u,v}$, the positionality score for the pair $\{u, v\}$, as:

$$\rho_{u,v} = \frac{6 \sum_r \sum_q s_d(M_{u,v,d,r,q})^2}{N_{boot} \sum_r \sum_d s_q(M_{u,v,d,r,q})^2} \tag{24}$$

This is a measurement of the ratio of the variance attributed to spacing compared with the variance attributed to uncertainty in inference (via bootstrapping). When $\rho_{u,v} \gg 1$, spacing-dependent effects are strong. We additionally perform significance testing using the Kruskal-Wallis H-test, over each of the four possible spacings $d \in \{-2, -1, 1, 2\}$.

###### 7.7.3 Identifying syntax modes with k-means clustering

Having identified cases of strong spacing-dependent effects, we wanted to map the 'modes' of these spacing rules onto a continuum. Specifically, our fitted model generates a discrete function $\alpha_{u,v}(d)$ as a function of spacing $d$ (or for simplicity the vector $\boldsymbol{\alpha_{u,v}}$ of size 4).

To perform spatial clustering, we constructed a feature vector from the above. Specifically, we define:

$$\mathbf{Q}_{u,v} = \left\{ \sum_{d \in D} \alpha_{u,v} : D \subseteq \{-408, -204, 204, 408\}, 1 \leq |I| \leq 4 \right\} \tag{25}$$

Filtering by fragment-pairs where the positionality score exceeds $2$, we perform k-means clustering on $\boldsymbol{Q}_{u,v}$ on all unique pairs of fragments, with $n_{cluster} = 3$, using *sklearn.cluster.KMeans*. Given pairs can be considered in either order, we choose $u$ and $v$ such that $\alpha_{u,v}(d)$ is on average increasing in $d$.

###### 7.7.4 Orthogonal basis decomposition of syntax contributions

For a continuum description, we constructed a set of four orthonormal bases to describe $\alpha_{u,v}(d)$.

$$\hat{\boldsymbol{v}}_0 = (\frac{1}{2}, \frac{1}{2}, \frac{1}{2}, \frac{1}{2}) \tag{26}$$

$$\hat{\boldsymbol{v}}_1 = (0, -\frac{\sqrt{2}}{2}, \frac{\sqrt{2}}{2}, 0) \tag{27}$$

$$\hat{\boldsymbol{v}}_2 = (-\frac{1}{2}, \frac{1}{2}, \frac{1}{2}, -\frac{1}{2}) \tag{28}$$

$$\hat{\boldsymbol{v}}_3 = (\frac{\sqrt{2}}{2}, 0, 0, -\frac{\sqrt{2}}{2}) \tag{29}$$

from which one can note immediately that $\hat{\boldsymbol{v}}_0$ specifies the average interaction over spacing, and the other three vectors describe the grammatical modes. The coefficients of this transformed vector space ($m_{u,v,k} = \hat{\boldsymbol{v}}_k \cdot \boldsymbol{\alpha_{u,v}}$) thus define the contributions of each of these basis vectors to the overall shape of $\alpha_{u,v}(d)$. Noting that $Var_d(\alpha_{u,v}(d)) = \sum_{k=1}^{3} m_{u,v,k}^2$, one can therefore define the percentage of variance owing to each of the remaining three components $\omega_{u,v,k} = m_{i,j,k}^2 / \sum_{l=1}^{3} m_{u,v,l}^2$. This therefore allowed us to plot the spacing-dependent rules of each pair of fragments on a ternary plot, describing the relative contribution of spacing-dependent variance with respect to each of the three remaining modes ($\hat{\boldsymbol{v}}_1, \hat{\boldsymbol{v}}_2, \hat{\boldsymbol{v}}_3$). To perform the calculations, we let $\alpha_{u,v}(d) = \frac{1}{2}(\beta_{SP}|_{u,v,d}^{r=1} + \beta_{SP}|_{u,v,d}^{r=2})$.

###### 7.7.5 Inferring fragment order-dependent effects from spacing rules

Given the above analysis, we sought to determine whether the spacing rules inferred from NeMECiS are sufficient to predict changes in SAG-dependent synCRE activity upon permuting the order of its comprising fragments. In this model, permutation of fragments leaves the contributions owing to individual fragments ($\beta_F$) and owing to spacing-independent synergies ($\beta_P$) unchanged, meaning $\beta_{SP}$ can be solely used to infer order-dependent effects. Put another way, this experimental test is determining whether pairwise contributions are sufficient to explain the activity of a synCRE as a whole.

For a given triple of fragment indices $\{u, v, w\}$, for which exhuastive permutation sets (size $6$) were generated and tested experimentally ($\mathcal{P}_{\{u,v,w\}} \mapsto \{(u, v, w), (u, w, v), ...\}$) we calculated the normalised activity of each ordering as:

$$\Delta y_{(u,v,w)} = \alpha_{u,v}(d = 1) + \alpha_{v,w}(d = 1) + \alpha_{u,w}(d = 2) \tag{30}$$

$$\Delta \hat{y}_{(u,v,w)} = \Delta y_{(u,v,w)} - \langle \Delta y_{(p,q,r)} \rangle_{(p,q,r) \in \mathcal{P}_{\{u,v,w\}}} \tag{31}$$

This was performed on coefficients from each replicate individually (defining the lower and upper error-bars), as well as their average.

###### 7.7.6 FL-synCRE activity predictions

Extending the above, we next asked whether our model could extrapolate to larger permutation sets, testing if the spacing-dependent rules learned from NeMECiS could generalise to arbitrarily lengthed sequences. To do this, we took $7\times$ contiguous fragments comprising the Olig2 CRE (fragments $8, 9, ..., 14$) and calculated the SAG-dependent activities upon exhaustive permutation ($N_{\text{Design Space}} = 7! = 5040$). Defining $\boldsymbol{f} = \{f_1, f_2, ..., f_7\}$ as a particular ordering of fragments $8-14$, we could calculate the normalised activities upon shuffling (including the natural ordering where $\boldsymbol{f} = (8, 9, ..., 14)$) as:

$$\Delta y_{\boldsymbol{f}} = \sum_{i=1}^{6} \alpha_{f_i,f_{i+1}}(d = 1) + \sum_{i=1}^{5} \alpha_{f_i,f_{i+2}}(d = 2) \tag{32}$$

This was performed on the average of coefficients across replicates, as well as each replicate in turn to determine reproducibility.

###### 7.7.7 Modelling of individualised synCRE flow cytometry analysis

For the individualised tests of the multiplicative model, considering the exhaustive construction of triplets of fragments taken with replacement from a pair of fragments $\{i, j\}$, we fit the function $\Delta y_{N_i,N_j} = exp(\alpha + N_i\beta_i + N_j\beta_j)$ (where $N_i + N_j = 3$) using `scipy.optimize.curve_fit`. For analyses comparing the number of inducing fragments against $log\ \Delta y_{N_i,N_j}$, we use the linear regression function `scipy.stats.linregress`.

For the individualised tests for interactions between fragments 2 and 16, we fit an expanded integrative model on all triplets containing $\{2, 8\}$ and $\{16, 8\}$ (where $8$ is a neutral fragment and $2$ and $16$ are inducing fragments): $\Delta y_{N_2,N_8,N_{16}} = exp(\alpha + N_2\beta_2 + N_8\beta_8 + N_{16}\beta_{16})$ (where $N_2 + N_8 + N_{16} = 3$). To determine sensitivity to experimental error, we performed a bootstrap analysis, re-fitting the above model to the data sampled evenly by synCRE structure, but sampled with replacement for 500SAG-0SAG comparisons ($N_{boot} = 1000$), plotting the interval $\Delta y_{pred} \pm \sigma_{\Delta y_{pred}}$.

### 8 Comparison of deep learning approaches in predicting synCRE activity

#### 8.1 Overview

We evaluated the predictive performance of four convolutional neural network (CNN) based deep learning models – ExplaiNN [Novakovsky et al., 2023], DeepCNN [Kelley et al., 2016], DanQ [Quang and Xie, 2016], and DeepSTARR [de Almeida et al., 2022] – for predicting the activity of cis-regulatory elements (CREs) with and without SAG treatment. These models have been widely used in genomic applications, including transcription factor binding prediction, chromatin accessibility inference, and enhancer activity characterization [Novakovsky et al., 2023, Kelley et al., 2016, Quang and Xie, 2016, de Almeida et al., 2022].

#### 8.2 Model Architecture

Each model received as input a one-hot-encoded representation of synCRE sequences and was trained to predict the corresponding CRE activity levels with and without SAG treatment. The architectures were adapted from the original publications [Novakovsky et al., 2023] with modifications tailored to the task as follows.

##### 8.2.1 ExplaiNN

ExplaiNN was implemented as described in [Novakovsky et al., 2023] and consisted of units with a single convolutional layer followed by fully connected layers. The architecture included:

- a convolutional layer with one 19×4 filter, batch normalization, an exponential activation function, and max pooling (kernel size = 7, stride = 7);
- a fully connected layer with 100 nodes, batch normalization, ReLU activation, and 30% dropout;
- a second fully connected layer with a single node, batch normalization, and ReLU activation;
- a final linear output layer predicting CRE activity with and without SAG treatment.

##### 8.2.2 ConvNetDeep (DeepCNN)

ConvNetDeep, referred to as DeepCNN in [Novakovsky et al., 2023]) and originally presented in [Kelley et al., 2016], was adapted by modifying filter sizes and the final output layer. In detail, this implementation included

- three convolutional layers with 100 filters each, using kernel sizes of 19×4, 11×1, and 7×1, respectively, each followed by batch normalization, ReLU activation, and max pooling (kernel size = 3, stride = 3);
- two fully connected layers with 1,000 nodes, batch normalization, ReLU activation, and 30% dropout;
- a final fully connected layer generating two outputs.

##### 8.2.3 DanQ

DanQ [Quang and Xie, 2016] integrates convolutional and recurrent layers to capture dependencies. The original DanQ architecture was altered to incorporate a fully connected output layer producing two outputs[Novakovsky et al., 202[illegible]
The architecture included:

- a first convolutional layer with 320 filters (26×4), ReLU activation, 20% dropout, and max pooling (kernel size = 13, stride = 13);
- two bidirectional long short-term memory (biLSTM) layers with a hidden state size of 320 and 50% dropout
- a fully connected layer with 925 nodes and ReLU activation;
- a final fully connected output layer generating two outputs.

##### 8.2.4 DeepSTARR

DeepSTARR [de Almeida et al., 2022, Novakovsky et al., 2023], was implemented as follows:

- four convolutional layers with 256, 60, 60, and 120 filters, respectively, each using batch normalization, ReLU activation, and max pooling;
- two fully connected layers with 256 nodes each, batch normalization, ReLU activation, and 40% dropout;
- a final fully connected layer generating two outputs.

All models were implemented in PyTorch [Paszke et al., 2019].

#### 8.3 Data processing and model training

##### 8.3.1 Data processing

We first filtered the data following the criteria outlined in Section 3.3. Raw fluorescence measurements for CRE sequences were then preprocessed by averaging replicate values to obtain a single activity measurement per sequence. This resulted in 14,301 data-points of synCRE sequences paired with their corresponding CRE activity values with and without SAG. Data were split into training, validation, and test sets in a 7:1.5:1.5 ratio.

Each dataset was loaded using the PyTorch DataLoader function. Training data were shuffled during each epoch to enhance generalization, whereas validation and test sets remained fixed to ensure consistency.

##### 8.3.2 Hyperparameter optimization

Hyperparameters were optimized independently for each model.

- **ExplaiNN**. The number of convolutional units (`num_cnns`), batch size (`bs`), and learning rate (`lr`) were explored over the following ranges: $\texttt{num\_cnns} \in \{10, 20, 30, \ldots, 250\}$, $\texttt{bs} \in \{96, 168, 322\}$ and $\texttt{lr} \in \{5 \times 10^{-5}, 1 \times 10^{-4}, 5 \times 10^{-4}\}$
- **Others**. For ConvNetDeep, DanQ, and DeepSTARR we optimized the batch size (`bs`) and learning rate (`lr`) over: $\texttt{bs} \in \{96, 168, 322\}$ and $\texttt{lr} \in \{1 \times 10^{-4}, 5 \times 10^{-4}, 1 \times 10^{-3}, 5 \times 10^{-3}\}$.

Each model was trained five times per hyperparameter setting with different random seeds.

##### 8.3.3 Model Training

All models were trained for up to 200 epochs with early stopping to prevent overfitting. An early stop was triggered if the validation loss (loss calculated for the validation set) did not improve for 15 consecutive epochs.

The validation loss was computed using mean squared error (MSE):

$$\text{MSE} = \frac{1}{n} \sum_{i=1}^{n} (y_i - \hat{y}_i)^2$$

where $y_i$ is the true label, $\hat{y}_i$ is the predicted label ($i \in \{1, 2\}$ in this case), and $n$ is the number of data points in the validation set. The total loss was the sum of MSE values for CRE activity with and without SAG:

$$\text{Total Loss} = \text{MSE}_{\text{with\_SAG}} + \text{MSE}_{\text{without\_SAG}}$$

##### 8.3.4 Model evaluation

Performance was assessed using Pearson correlation between predicted and true activity values on the test set:

$$\text{pearson}_{\text{corr}} = \frac{\sum_{i=1}^{n} (y_i - \overline{y})(\hat{y}_i - \overline{\hat{y}})}{\sqrt{\sum_{i=1}^{n} (y_i - \overline{y})^2} \sqrt{\sum_{i=1}^{n} (\hat{y}_i - \overline{\hat{y}})^2}}$$

where $y_i$ and $\hat{y}_i$ still denote the true and predicted labels, $\overline{y}$ is the mean of $y_i$, and $\overline{\hat{y}}$ is the mean of $\hat{y}_i$. The epoch with the highest Pearson correlation was selected as the best-performing model.

### 9 Motif analysis

#### 9.1 Motif calling

Pulling all vertebrate non-redundant `.pfm` motif files from https://jaspar.elixir.no/downloads/ [Rauluseviciute et al., 2024], we ran `MOODS` ([Korhonen et al., 2009],https://github.com/jhkorhonen/MOODS) to enrich for motifs in each of the twenty-five 200bp fragment sequences using the following command:

```
python MOODS/python/scripts/moods-dna.py -m reference/JASPAR2024_CORE_PROCESSED/*.pfm
-s fasta/fragment_i.fa -p 0.0001 > alignment/fragment_i.csv
```

This returns `.csv` files with the following columns: `[Fragment, Motif, Start Position, Orientation, Match Score, Matched Sequence]`.

#### 9.2 Clustering of motifs into archetypes

Given substantial sequence similarity among sets of motif sequences, we followed [Vierstra et al., 2020] by clustering motifs based on sequence similarity into 'archetypes'. Specifically, we pulled pre-computed pairwise similarity scores from `JASPAR2024` and performed agglomerative clustering (`sklearn.cluster.AgglomerativeClusteri` `n_clusters=150`. This provided a grouping of motifs by similarity, which were used for subsequent feature-enrichment analysis.

#### 9.3 Motif enrichment analysis

We next scored whether particular motif clusters showed a correlation with SAG-dependent activity. Specifically, we took the `MOODS`-enriched motifs from each fragment, and subsetted by `Match Score`>6.5. We then built a series of sparse, boolean matrices: $\boldsymbol{M_{mtf,arch}}$, a motif to archetype association built on the above clustering; and $\boldsymbol{M_{frag,mtf}}$, a fragment to motif matrix; and $\boldsymbol{M_{synCRE,frag}}$, a boolean matrix determining the presence of a synCRE. From this, the below matrix product describes the archetype-count matrix (where $B(\boldsymbol{M})$ describes a boolean transformation).

$$B(\boldsymbol{M_{synCRE,frag}M_{frag,mtf}M_{mtf,arch}}) = \boldsymbol{M_{synCRE,arch}} \tag{33}$$

We then calculate the Spearman's correlation coefficient across synCREs between the presence of a given archetype and $\Delta y$ (change in predicted fluorescence with respect to SAG) for each replicate separately, filtering synCRE data-points by quality as in our linear modelling framework. We subsequently filtered for archetypes for which at least one of the corresponding genes is expressed (TPM$> 20$ for any day 4, 5, 6 sample from [Delas et al., 2023]). Our strongest enriched archetype was cluster `90`, containing the motifs [Gli1, Gli2, Zbtb7b, Zbtb7c].

### 10 Dynamical modelling of neural tube patterning

We use the previously established and parameterised model of neural progenitor patterning [Cohen et al., 2014, Pezzotta and Briscoe, 2023, Exelby et al., 2021] to explore the effects of changes in expression amplitude in response to a gradient of Shh. This describes the gene-regulatory network underpinning Shh morphogen decoding during neural progenitor patterning as a set of ordinary differential equations for the level of four transcription factors $\boldsymbol{S} = (P, O, N, I)$, corresponding to Pax6, Olig2, Nkx22, and Irx3 respectively. The Shh gradient is integrated into the model by graded activation of a fixed amount of Gli ($G$) protein ($G = A + R = 1$), which was either in an activated ($A$) is repressive ($R$) form.

$$\frac{d\boldsymbol{S}}{dt} = f_{\boldsymbol{S}} = (f_P, f_O, f_N, f_I) \tag{34}$$

$$H^+(y) = \frac{y}{1+y} \tag{35}$$

$$A = e^{-0.15x} \tag{36}$$

$$R = 1 - A \tag{37}$$

$$f_P = \alpha_{Pax} H^+ \left( \frac{K_{Pax,Pol} c_{Pol}}{(1 + K_{Pax,Oli} O)^2 (1 + K_{Pax,Nkx} N)^2} \right) - \beta_{Pax} P \tag{38}$$

$$f_O = \alpha_{Oli} H^+ \left( \left[ \frac{K_{Oli,Pol} c_{Pol}}{(1 + K_{Oli,Nkx} N)^2 (1 + K_{Oli,Irx} I)^2} \right] \left[ \frac{1 + f_A K_{Oli,Gli} A}{1 + K_{Oli,Gli}(A + R)} \right] \right) - \beta_{Oli} O \tag{39}$$

$$f_N = \alpha_{Nkx} H^+ \left( \left[ \frac{K_{Nkx,Pol} c_{Pol}}{(1 + K_{Nkx,Pax} P)^2 (1 + K_{Nkx,Oli} O)^2 (1 + K_{Nkx,Irx} I)^2} \right] \left[ \frac{1 + f_A K_{Nkx,Gli} A}{1 + K_{Nkx,Gli}(A + R)} \right] \right) - \beta_{Nkx} N \tag{40}$$

$$f_I = \alpha_{Irx} H^+ \left( \frac{K_{Irx,Pol} c_{Pol}}{(1 + K_{Irx,Oli} O)^2 (1 + K_{Irx,Nkx} N)^2} \right) - \beta_{Irx} I \tag{41}$$

In all cases, the model was simulated using *scipy.integrate.solve_ivp* subject to an initial condition of $\boldsymbol{S} = (0.1, 0, 0, 0.1)$.

Varying the parameter $\alpha_{Olig2}$ for a fixed range ($x \times 10^q$, where $q$ is linearly evenly spaced between X and Y), we measured the predicted distribution of $O$ (Olig2) as a function of position, identifying shifts in the patterning.

#### 10.1 Fitting FL-synCREs

To predict the changes in Olig2 spatiotemporal expression upon GRN rewiring, we calibrated the model. In vitro differentiation under SAG of neural progenitors yields multiple cell fates that shift in proportions as the concentration of SAG changes. Thus to calibrate the in vivo model to the in vitro differentiation, we used fate proportions of p3, pMN and p2+ cells at day 6 500nM SAG determined in [Delas et al., 2023] (p3=$20\%$; pMN=$45\%$; p2+ = $35\%$). By binarising cell fate in the in silico model as a function of dorsoventral position $y$, we specify pMN$^+(y) = O(y) > \max_y(O(y))/2$, p3$^+(y) = N(y) > \max_y(N(y))/2$ and p2$^+(y) = \neg($pMN$^+(y) +$ p3$^+(y))$. We then suggest that 500nM SAG in vitro differentiations maps to a Gaussian kernel of spatial coordinates in the in vivo model:

$$K(y) = \exp\left( \frac{-(y - y_0)^2}{\sigma_y^2} \right) \tag{42}$$

From this, we can determine the values of $y_0$ and $\sigma_y$ for which the following cost-function is minimised using a grid-search.

$$\mathcal{C} = \left( \frac{\int_0^1 \text{pMN}^+(y)K(y)dy}{\int_0^1 \text{p2}^+(y)K(y)dy} - \frac{P(\text{pMN})}{P(\text{p2+})} \right)^2 + \left( \frac{\int_0^1 \text{p3}^+(y)K(y)dy}{\int_0^1 \text{p2}^+(y)K(y)dy} - \frac{P(\text{p3})}{P(\text{p2+})} \right)^2 \tag{43}$$

Then using the measured ZsGreen fluorescence values, we calculated the fold changes with respect to the WT Olig2 CRE for the boosted and diminished FL-synCREs. We then calculated the kernel-averaged value of Olig2 expression for 500nM SAG as a function of $\alpha_{Oli}$, normalised to the WT value:

$$\tilde{O}(\alpha_{Oli}) = \int_0^1 O(y|\alpha_{Oli}) K(y) dy \tag{44}$$

$$\hat{o}(\alpha_{Oli}) = \frac{\tilde{O}(\alpha_{Oli})}{\tilde{O}(\alpha_{Oli} = \text{WT value})} \tag{45}$$

From this, we can compute $\alpha_{Oli}$ for the FL-synCREs as the value that satisfies:

$$\hat{o}(\alpha_{Oli}^{\text{FL-synCRE}}) = \frac{\text{ZsGreen}_{\text{FL-synCRE}}}{\text{ZsGreen}_{\text{WT}}} \quad (46)$$

To determine uncertainty, error was propagated for $\text{ZsGreen}_{\text{FL-synCRE}}$, $\text{ZsGreen}_{\text{WT}}$.

### 11 Image analysis

#### 11.1 Pre-processing and segmentation

We exported a $4\times$ $x, y$ downsampled version of our 3D images inherent to the pyramidal `.vsi` files from Olympus Spinning Disk microscopy to `.ome.tiff` using the `Bio-Formats` [Linkert et al., 2010] converter in Fiji-ImageJ [Schindelin et al., 2012], ensuring that physical spatial dimensions are preserved.

We then repurposed the cell-volume segmentation software `giani` [Barry et al., 2022] to segment entire organoids from the 3D imaging volumes, using the following command:

```
srun java -Xmx750G -jar path-to-giani.jar path-to-joblist.txt path-to-properties.xml
index
```

This produces a directory of `Nucleus_masks`, with `.tiff` files for each z-slice (where here nuclei imply organoids). We use these masks to compile a series of `.tiff` files for each uniquely identified organoid, taking the bounding-box of a given organoid index, and concatenating into it the original microscopy data. This now allows for parallelisation of subsequent analysis for each organoid.

From here, we perform a set of operations to clean the mask by organoid:

- Perform `scipy.ndimage.binary_erosion` on the mask with `iterations=2`
- Rescale the image and mask such that each voxel has dimension $(1\mu m \times 1\mu m \times 1\mu m)$ using `scipy.ndimage.zoom` with `order=0`.

#### 11.2 Organoid geometry metrics

From our cleaned, isometric organoid mask ($\boldsymbol{M}$), we calculated a series of geometric metrics:

- Volume ($V_{mask} = \sum_{\{x,y,z\}} M_{i,j,k}$) using `mask_iso.sum()` [units = $\mu m^3$]
- Convexity ($\mathcal{C} = V_{hull}/V_{mask} - 1$), where we calculate the convex hull of the mask, and compare its volume to the mask volume. The ratio $V_{hull}/V_{mask}$ will tend to $1$ when the mask is perfectly convex (i.e. all boundary pixels lie on their convex hull).

#### 11.3 FoxA2+ floor-plate scoring

We next scored each organoid for their floor-plate composition. As an overview, we generated a mask for where the FoxA2 channel exceeds a critical intensity, and sequentially processed it by merging clusters and filling holes, such that we can group sets of FoxA2 voxels as being members of distinct floor-plates.

Specifically, we performed the following operations on the isometrised FoxA2 channel:

- Define $\boldsymbol{M}^{FA2}$, the mask of FoxA2+ voxels within its isometrised image, as $M_{i,j}^{FA2} = \{1 \text{ if } F_{i,j} > F_{thresh}; \text{ else } 0\}$, where $F_{thresh} = 1000$.
- Perform the operation `process_3d_mask`. This identifies the connected components with `scipy.ndimage.label`, and from the corresponding labelled mask, performs `scipy.ndimage.remove_small_objects` with `min_size=200`.
- Smooth the mask by performing `scipy.ndimage.gaussian_filter`, then binarise with the threshold $0.1$.
- Perform `process_3d_mask` a second time, this time with `min_size=3000`.
- We update our FoxA2+ voxel classification (removing the spurious positive pixels removed in the above smoothing) by taking the intersection between $\boldsymbol{M} \otimes \boldsymbol{M}^{FA2} \otimes \boldsymbol{M}_{processed}^{FA2} = \boldsymbol{M}_{final}^{FA2}$

- Use `scipy.ndimage.label` to label the product of the above. This label mask is used to classify voxels in $\boldsymbol{M}^{FA2}_{final}$, and the corresponding number of connected components is taken as the number of floor-plates.

##### 11.4 Organoid quality control filtering

We perform analysis only on organoids with regular and equivalent geometries, and with precisely one floor-plate, for valid subsequent comparisons. Specifically, we require:

- Number of floor plates = 1
- Volume constraints: $10^6/3 < V_{mask} < 3 \times 10^3$
- Convexity constraints: $\mathcal{C} < 0.5$

##### 11.5 Olig2 spatial expression analysis

We then determine the euclidian distance from each voxel within the processed isometrised organoid mask to its nearest FoxA2+ voxel in $\boldsymbol{M}^{FA2}_{final}$ using `scipy.ndimage.distance_transform_edt` on the inverse of $\boldsymbol{M}^{FA2}_{final}$. We then bin these distances to a linear range of bins ($\{0, 5, 10, ...\}\mu m$), converting exact distances to bin mid-points (i.e. $\{2.5, 7.5, ...\}\mu m$), excluding voxels that lie outside of $\boldsymbol{M}$, the processed isometrised organoid mask.

We then perform spatial analysis on the distribution of Olig2 expression. Specifically, we binarise the (isometrised) Olig2 channel with a threshold $3000$. From this, we can compare the fraction of Olig2+ voxels at each spatial bin position in an organoid-by-organoid fashion ($p_j^{Olig2}(x)$, where $x$ is a set of distances (bin mid-points) from the nearest FoxA2+ voxel, for organid $j$). To understand the average behaviour by genotype, we take the mean of this spatial profile (i.e. effectively removing an otherwise inherent weighting based on variations in organoid volume; $p^{Olig2}_{ensemble}(x) = \frac{1}{N_{organoid}} \sum_{j=1}^{N_{organoid}} p_j^{Olig2}$). We report these values in a replicate-by-replicate fashion, excluding replicates for which $N_{organoid} \leq 20$.

For domain-size calculations, we calculate a binary spatial profile for positions where $p^{Olig2}$ exceeds a threshold percentage ($b_j^{Olig2}(x) = (p_j^{Olig2}(x) > 0.1)$). This is then processed such that $b_j^{Olig2}(x > x_0) = 0$ where $x_0$ is the lowest value of $x$ for which $b_j^{Olig2} = 0$, so as to consider only called Olig2 domains near the floor-plate source, removing rare instances where iso-lines of Olig2 expression exceed the threshold far from the source (likely due to imaging noise). We then take a per-replicate ensemble mean as a function of $x$: $\langle b_j^{Olig2}(x)\rangle_j$. Next, we calculate $d^{Olig} = \sum_{x \in \text{bin mid-points}} \left[x \cdot \langle b_j^{Olig2}(x)\rangle_j\right] / \sum_{x \in \text{bin mid-points}} \langle b_j^{Olig2}(x)\rangle_j$ to determine the domain size by organoid. This is then normalised by the wild-type domain size averaged across replicates $\tilde{d}^{Olig2}_{cond.,rep.} = d^{Olig2}_{cond.,rep}/\langle d^{Olig2}_{WT,rep.}\rangle_{rep.}$.

#### 12 Relating thermodynamic state ensemble models to synCRE activity

##### 12.1 Transcriptional activation by a single site

We started from a basic model of gene regulation, which offers a baseline framework for understanding the composability rules of regulatory sequences. This approach draws heavily from prior work [Bintu et al., 2005, Buchler et al., 2003, Veitia, 2003]. We consider a regulatory element containing a single binding site for a transcriptional activator $A$ and a binding site for RNA polymerase II (RNAPII) $P$ at the transcription start site (TSS). Additionally, we assume that $A$ and $P$ can physically interact when bound simultaneously. The system can occupy four distinct states: neither molecule bound, only A bound, only P bound, or both bound. Under the assumption of thermodynamic equilibrium, the partition function describing this system can be written as:

$$Z = 1 + Ae^{-E_a/kT} + Pe^{-Ep/kT} + APe^{-(E_a+E_p+E_{ap})/kT} \tag{47}$$

where $T$ is the temperature and $k$ is the Boltzmann constant. Here $E_{xy}$ relates to a binding/unbinding process by:

$$X + Y \underset{k_{xy}^-}{\overset{k_{xy}^+}{\longleftrightarrow}} XY \tag{48}$$

$$k_{xy}^+/k_{xy}^- = K_{xy} = e^{-E_{xy}/kT} \tag{49}$$

$$\tag{50}$$

From this, the probability that RNAPII is bound can be calculated by:

$$p_{bound} = \frac{Pe^{-Ep/kT} + APe^{-(E_a+E_p+E_{ap})/kT}}{Z} \tag{51}$$

$$= K_p P(1 + K_a A e^{-E_{ap}/kT})/(1 + K_a A + K_p P + K_a K_p A P e^{-E_{ap}/kT}) \tag{52}$$

$$= \frac{1}{1 + \frac{1}{K_p P} \cdot \frac{1+K_a A}{1+K_a A e^{-E_{ap}/kT}}} \tag{53}$$

which can be written more simply as

$$p_{bound} = \frac{1}{1 + \frac{\alpha}{F_A}} \tag{54}$$

$$\alpha = \frac{1}{K_p P} \tag{55}$$

$$F_A = \frac{1 + f_a K_a A}{1 + K_a A} \tag{56}$$

$$f = e^{-E_{ap}/kT} \tag{57}$$

$$\tag{58}$$

This reparameterisation provides an intuitive interpretation [Bintu et al., 2005], where $F_A$ quantifies the regulatory effect of $A$ on polymerase binding. When $F_A < 1$, the polymerase binds less frequently than in the absence of the TFBS, indicating that $A$ acts as a repressor. In contrast, if $F_A > 1$, the polymerase is more likely to bind, implying that $A$ functions as an activator. To express this in terms of the transcription rate $r$, we assume that polymerase recruitment is the rate-limiting step in transcription initiation, such that:

$$r = \beta_m \cdot p_{bound} \tag{59}$$

Consequently, assuming the system is at steady state, the mean level of the corresponding protein is a read-out for $p_{bound}$

$$\frac{d[mRNA]}{dt} = r - \lambda_m [mRNA] \tag{60}$$

$$\frac{d[Protein]}{dt} = \beta_p [mRNA] - \lambda_p [Protein] \tag{61}$$

$$\implies [Protein] = \frac{\beta_p \beta_m}{\lambda_m \lambda_p} p_{bound} = c \cdot p_{bound} \tag{62}$$

#### 12.2 Varying number of independently acting transcription factors

What happens when there are two different transcription factors $A$ and $B$ with two corresponding TFBSs? Suppose $A$ and $B$ are not directly interacting, such that binding is not cooperative. If we further assume that $A$ and $B$ independently interact with RNAPII, this yields a partition function:

$$Z_{AB} = 1 + K_a A + K_b B + K_p P + K_a K_b AB + K_a K_p f_a AP + K_b K_p f_b BP + K_a K_b K_p f_a f_b ABP \tag{63}$$

implying:

$$F_{AB} = \frac{(1 + f_a K_a A)(1 + f_b K_b B)}{(1 + K_a A)(1 + K_b B)} \tag{64}$$

$$p^{AB}_{bound} = \frac{1}{1 + \frac{\alpha}{F_{AB}}} \tag{65}$$

Comparing a CRE with either of the two TFBSs to the two-site CRE, one can see that individual "contributions" (i.e. $F$s) multiply:

$$F_{AB} = F_A \cdot F_B \tag{66}$$

Therefore, the function of a cis-regulatory element made up of a set $S$ of non-interacting TFBSs is expected to follow a logistic curve based on their individual effects.

$$p^{\{S\}}_{bound} = \frac{1}{1 + exp(-(log\ 1/\alpha + \sum_{s\in\{S\}} log\ F_s))} \tag{67}$$

Additionally, when $F_s \ll \alpha$, meaning RNAPII loading to the TSS is far from saturation, we can approximate $p_{bound}$ as the product of the individual contributions of each TFBS. As a result, the activity of a CRE consisting of different TFBSs should be directly proportional to the product of the activities of each TFBS independently.

$$p^{\{S\}}_{bound} \approx \frac{1}{\alpha} \prod_{s\in\{S\}} F_s \propto \prod_{s\in\{S\}} p^s_{bound} \tag{68}$$

When multiple identical TFBSs are concatenated, with $F_s$ held constant, the activity will increase exponentially with the number of binding sites, following an exponential distribution.

$$p_{bound}(N) \approx \frac{1}{\alpha} F^N \propto exp(N \cdot log\ F) \tag{69}$$

This outcome also suggests that combining two CREs, each containing distinct sets of TFBSs $S_1$ and $S_2$, should adhere to a multiplicative rule, a principle that can be generalized to any N-way fusion.

$$p^{\{S_1,S_2\}}_{bound} \propto p^{\{S_1\}}_{bound} p^{\{S_2\}}_{bound} \tag{70}$$

Together, the activity of a regulatory sequence consisting of independently acting TFBSs (those that independently bind and recruit RNAPII) adheres to a logistic composability rule based on the separate contributions of its TFBSs. In sub-saturating conditions, this behavior simplifies to a multiplicative rule. Consequently, when TFBSs act independently, the activity of any fusion of sequence fragments—regardless of the number or type of TFBSs—can be approximated by multiplying the activities of each individual fragment.

#### 12.3 Breaking independence requirements

The previous section supports a multiplicative (with saturation) composability rule based on the contributions of TFBSs. However, it assumes that TFs act independently, both in terms of binding to the CRE and recruiting the polymerase. To relax these assumptions, we introduce additional energies that account for cooperative binding and post-binding synergies. For a CRE containing two TFBSs, this can be generically represented by the partition function.

$$Z_{AB} = 1 + K_a A + K_b B + K_p P + c_{ab} K_a K_b AB + f_a K_a K_p AP + f_b K_b K_p BP + c_{ab} f_{ab} f_a f_b K_a K_b K_p ABP \tag{71}$$

Here, $c_{ab}$ quantifies how much the likelihood of both TFs binding together deviates from the expected probability if each TF were to bind independently, assuming the polymerase is not bound. If $c_{ab} \neq 1$, this signifies that the binding of the two TFs is cooperative.

$$c_{ab} = \frac{P(A \cap B|\neg P)}{P(A \cap \neg B|\neg P)P(B \cap \neg A|\neg P)} \tag{72}$$

$f_{ab}$ reflects the difference in polymerase recruitment probability when both TFs are present, as compared to their individual recruitment abilities. When $f_{ab} \neq 1$, it indicates that the recruitment process is cooperative.that the TFs engage in post-binding synergy.

$$f_{ab} = \frac{1}{c_{ab}} \frac{P(A \cap B|P)}{P(A \cap \neg B|P)P(B \cap \neg A|P)} \tag{73}$$

We can then determine $F_{AB}$:

$$F_{AB} = \frac{1 + f_a K_a A + f_b K_b B + f_{ab} c_{ab} f_a f_b K_a K_b AB}{1 + K_a A + K_b B + c_{ab} K_a K_b} \tag{74}$$

This expression is no longer the product of $F_A$ and $F_B$.

$$\frac{F_{AB}}{F_A F_B} = \frac{1 + (f_{ab} c_{ab} - 1)\left(\frac{f_a f_b}{F_A F_B}\right)\left(\frac{K_a K_b AB}{(1+K_a A)(1+K_b B)}\right)}{1 + (c_{ab} - 1)\left(\frac{K_a K_b AB}{(1+K_a A)(1+K_b B)}\right)} \tag{75}$$

The regulatory complexity in principle grows exponentially with the addition of more TFBSs. For instance, the partition function for a sequence with three TFBSs will include cooperative binding and post-binding synergy terms for each of the two possible interacting TFBS pairs ($c_{ac}, c_{bc}, f_{ac}, f_{bc}$), as well as higher-order terms ($c_{abc}, f_{abc}$).

#### 12.4 Defining regulatory modules

One expects that, at least for larger regulatory elements, not all pairs of TFBSs engage in synergistic interactions (pre- or post-binding). Therefore, certain combinations of TFBSs may combine under the multiplicative rule. As a result, some TFBS combinations will follow the multiplicative rule, while others will show deviations from it. This observation leads to the concept of a regulatory module. In this view, a regulatory module is a group of TFBSs where no synergy exists between any TFBS in the module and any TFBS from another.

Consider a graph where the TFBSs in a CRE are represented as nodes, and the synergies (pre- or post-binding) between pairs or higher-order combinations are represented as edges. For instance, there would be an edge between nodes $A$ and $B$ if any of the following terms differ from $1$: $f_{ab}, c_{ab}, f_{abc}, c_{abc}, f_{abcd}, c_{abcd}, \dots$. Modules are therefore defined as the connected components of the graph.

Define the partition function of the TFBSs comprising a module as $Z_m$. A CRE is then defined by a set of modules. A generic description of the partition function for a CRE with an assortment of potentially interacting TFBSs for TFs $T = \{A_1, A_2, \dots, A_n\}$ is given by

$$Z = 1 + K_p P + \sum_{\{A_1, A_2, \dots, A_n\}} K_i A_i (1 + f_i K_p P) + \sum_{S \subseteq \{A_1, A_2, \dots, A_n\}, |S|>1} c_S \prod_{i \in S} K_i A_i \left(1 + f_S K_p P \prod_{i \in S} f_i\right) \tag{76}$$

$$= \left[1 + \sum_{A_i \in T} K_i A_i + \sum_{S \subseteq T, |S|>1} c_S \prod_{i \in S} K_i A_i\right] + K_p P \left[1 + \sum_{A_i \in T} f_i K_i A_i + \sum_{S \subseteq T, |S|>1} c_S f_S \prod_{i \in S} K_i A_i f_i\right] \tag{77}$$

$$\implies F_T = \frac{1 + \sum_{A_i \in T} f_i K_i A_i + \sum_{S \subseteq T, |S|>1} c_S f_S \prod_{i \in S} K_i A_i f_i}{1 + \sum_{A_i \in T} K_i A_i + \sum_{S \subseteq T, |S|>1} c_S \prod_{i \in S} K_i A_i} \tag{78}$$

If we can divide TFBSs into sets of modules ($M = \{M_1, M_2, M_3\}$, $M_1 = \{A_1, A_2, ..., A_{m-1}\}, M_2 = \{A_m, A_{m+1}, ...A_{p-1}\}, ...$), one can show that

$$Z = \prod_j \left[1 + \sum_{A_i \in M_j} K_i A_i + \sum_{S \subseteq M_j, |S|>1} c_S \prod_{i \in S} K_i A_i\right] + K_p P \prod_j \left[1 + \sum_{A_i \in M_j} f_i K_i A_i + \sum_{S \subseteq M_j, |S|>1} c_S f_S \prod_{i \in S} K_i A_i f_i\right] \tag{79}$$

This therefore implies that

$$F_{\{M_1,M_2,...\}} = \prod_j F_{M_j} \tag{80}$$

Importantly, this demonstrates that we can decompose the activity of a CRE into the product of the activity of regulatory modules. As with the individual, non-interacting, TFBS examples, in in the sub-saturating the limit, this implies that the activity of a CRE will be linearly proportional to the product of the activities of each individual module.

Crucially, this shows that the activity of a CRE can be broken down into the product of the activities of its regulatory modules. Similar to the case with individual, non-interacting TFBSs, in the sub-saturating limit, this means that CRE activity will be directly proportional to the product of the activities of each module.

$$p_{bound}(\{M_1, M_2, ...\}) \propto \prod_j p_{bound}(M_j) \tag{81}$$

#### 12.5 Linear modelling

Due to the combinatorial design of synCREs in the NeMECiS dataset, it is possible to directly connect experimental measurements to the thermodynamic model of modularity. Assuming the thermodynamic model accurately represents transcriptional regulation, and that ZsGreen mRNA and protein levels reach steady state—with ZsGreen protein levels being linearly proportional to fluorescence—it becomes feasible to predict the SAG-dependent fluorescence of a synCRE ($y^S; S \in -, +$). This fluorescence can be described in terms of the SAG-dependent regulatory contribution of a synCRE composed of fragments $A$, $B$, and $C$ at a given SAG concentration ($F^S_{ABC}$).

$$y^S \propto \frac{1}{1 + \alpha / F^S_{ABC}} \tag{82}$$

In this context, $F^S_{ABC}$ represents a scalar value rather than the function. Specifically, we consider the overall function $F_{ABC}(\boldsymbol{x})$, which is, in principle, a function of many components in the cell's state-space, $\boldsymbol{x}$, with measurements focusing on evaluating the overall function $F_{ABC}(\boldsymbol{x}(S))$ at two specific SAG concentrations: $F^-_{ABC} = F_{ABC}(\boldsymbol{x}(0))$ and $F^+_{ABC} = F_{ABC}(\boldsymbol{x}(500))$.

We first test the hypothesis that all fragments in the NeMECiS dataset behave as independent modules. Consequently, one can calculate the SAG-dependent regulatory contributions of each individual fragment, such as $F^S A$, $F^S B$, $F^S_C, ...$. Therefore:

$$F^S_{ABC} = F^S_A \cdot F^S_B \cdot F^S_C \tag{83}$$

Therefore, for a given SAG concentration, let each fragment contribute a regulatory value $F^S_i$ ($50$ parameters in total). A synCRE can be represented by a triplet of indices $\boldsymbol{X} = (X_1, X_2, X_3)$. For instance, the synCRE labeled "4-5-6" would correspond to $X_1 = 4$, $X_2 = 5$, and $X_3 = 6$. Consequently, we can express the ZsGreen fluorescence of the synCRE as:

$$log(y^S_{\boldsymbol{X}}) \approx c + \sum_{i=1}^{3} log(F^S_{X_i}) \tag{84}$$

Here, $c$ is a proportionality constant. As a result, the log-transformed ZsGreen levels of synCREs measured by NeMECiS ($\log(y^S_{\boldsymbol{X}})$) can be linked to the by-fragment weights ($\beta_i = \log(F^S_{X_i})$) through linear regression, where the intercept parameter ($c$) represents the intrinsic transcription rate of the promoter.

It is important to note that this represents an under-determined system with 51 parameters and 50 variables. Intuitively, since synCREs always consist of three fragments, it is not feasible to determine an absolute baseline. However, differences in $\beta$ are still meaningful. Thus, it is reasonable to choose a value for $c$ that reflects the average activity of synCREs across the dataset, with the values of $\beta$ capturing the deviations from this baseline. A common approach to address this is through regularization. Specifically, we chose to fit the model using Ridge regression, minimizing a cost function of the form:

$$C = \|\boldsymbol{Z\beta} + y_0 - \boldsymbol{Y}\|_2^2 + \lambda\|\boldsymbol{\beta}\|_2^2 \tag{85}$$

### 13 Tables

| CRE | Sequence | Genomic Coordinates (mm10) |
|---|---|---|
| Nkx22 | TTGGTTCTATTCATTTTTCCCTCTAACTGTCCTTGTTTGGGTTGCCTTAG<br>TCCAGGCGAAGCAGAACGCTCAGGCATCGAGTCCTCCAAGCTCACAGCTG<br>CTTGGGCCGCGGCGGCGCCCCCACCACCCCCGGGTTTCTAGAGCCTTGTG<br>ACAGCGCCTCACTCCGAGCTGCTCAGCGGCTTTCTGCAACTCTCTCTGCC<br>TCGCCCCCAGTATGTGACGTGGGTGACAATGGCCCAGGTTGGAGCGAGCC<br>CCACGTCGGCGCGTCCTGGGTGGTCGGACCCGGGCAAACACAAATACAAA<br>CCGATTGCTAAGCTGCGGACAATGAGGGAAATGTAGACAAATGTCCCGCT<br>CCTGTTGGAAGCCTTTGTCCAGGCCCGGTTTTTGCATTTATTTCAGTGGC<br>GAAATAATACATGATTGACGCTCTCTTTCAATGTGTCCTAACTGTTTGGA<br>ATAAATCTAAGGTTGTCCCTAGTTGTCATGGCATTCAACCCTTTTCAATG<br>GACTATCTCTTCATTCATTTCTGAATCCGTCCACAATATTAAGGAAACCC<br>CTTCATGTCGACGTTCCCCATTCCTCTTTCTCCTGCTTTTCTTTTCTCCT<br>TTCTTCCTCCCTTTTCCCCTTGCAAGAATTAGTATCTGGTTTGAAACGTG<br>TTCCTCAGACTCATTACTTTCTCTGTCTTCCCCCCTTCTGTTTTTCCAGC<br>TTCTTTCTCAGCGTTAGAGCCTGTTGCCTCTCCCTCCCTCTTTTCAGGC | chr2<br>147187985-147188733 |
| Olig2 | TGATTGCCAACTGCCTCCAACCCAACTTTGTAAGCCGAGCCCTCATCCCT<br>ACCCACTCCCGGTGTGTCAGATGGAACACCTAGGTGGCCCACGGGGACCT<br>CTGACCTCTATATCCTCTTCCTTTCTCCCTCCTTTGCTACTTTTCTACTG<br>GATAAAAGGAGAGAGTGAGAGATAATTAACAAAAAACATGGCCCCGGGAC<br>AATGAAGCAACTGGCCTTGGCCGGCAAGCAACGATCCTGGTTTTCTAGGT<br>AGAGTTTCTCCCATCAATCTTTCCTTTAACCTCCCTGTTCGTGGAAGCAA<br>TAGAAACACCACCCCCTCCCCTGAGCAAATGCTTTCTTTTGACTGGAAAC<br>AAAAAGGGGGCCCGGCAAAGACGGAGGTGAAATCTGGGTGGTATGGGCAC<br>CGCACAATGGCCCCGCTGTTCCTGGCCCTGCTTGTGTTTTACAACAGGGG<br>AGGGGCAGGCGCGAATGGTCCGATGGTGGAGACAATCCCCCTGATTCAGG<br>CTACAAATGCATCTTCTATTCCACACGGAGCTGAGCAGAAAGGATGGGGG<br>TGACAAAGAGCATGGGCGGGGAGAGGGAAAACAAAATGTTTTCAGTTGAA<br>AAAAAAAATCTCTCATATCCTACACATCCTCAGAAGAGCTTCTATGGAGA<br>AGGCCTTCGGAGAGTCCCAGCCCACAACTCAAGGGCTTTGTCTGAACTCT<br>GATTTATTGATGAAGCTTAAGCGGCTCGCTAAGAAAGGCCTGGGGGTGTC<br>TTTGTCTTGAAGATAAAGTACAATAGGCCACAAGGGCCAAGATCTCTCGG<br>GATGCTCTCAGGTCCTGCCTCTCTCTTGCCCTCTCCTCCCTGCAAATGCC<br>AGCAGATGCTGAAAAAAAAAAAAACCCATCGGTGGTGTGGCTGGGAGTGC<br>TGGGGACAAGCTGGGCCACTTGAGGTCTCCTTAAGAGGGTATTATGGCCA<br>GGGAAAATATTTGCGCTCTAAGGATGGCACACTCCATTTGATAATGGCTC<br>TCATCTGCCTCAGATAATCGCCTCCCTCCCGGCTGTCAGGGGTGCAGCCA<br>CTGCCAATTCACAGCGCCCTCCGAGAAAGTACCCTTGTCTGTGATGACCA<br>AGATGGGGACATTGTGTTTACCTACTTGAGCAGAGGAGAAGGTGACCGTG<br>AGGGCAGCCTGCATTGTAAATTACAATTAAAACAGAAACAGACAGTTCCT<br>GCTCTGCCCTGGGACCCCCACCAATAAATTATGGGTGGACATTAGGGGAG<br>AGCCCAGGAAAGGTTGGGTCCTGGGGAGGATCCCCCCATCCCATAGCCTA<br>CCGAC | chr16<br>91192436-91193740 |

| | | |
|---|---|---|
| Pax6 | CTAAATAGCACCGCGGCGCCCGCTCTCCGGACAGTGATTAATGATAGCAG<br>TGCAGAGGGGTTAACACACTTCGCTGAAAAGTCTGTTGACTGAGCTTCTT<br>GTAACACAATGTGGCCCGCTGCACGCCTCGAGAGAATCCTTTTGTTGTCC<br>GTGCTTATTGTGGCCTCAAAATTCTGCCCACGAAAGTTTGCCAACGCTCC<br>TGCCCCAGGAGTTTAATAGTTTCCCTTACTCGCGGGGCATTGTGTGGTGC<br>TGAAAAGCAGCCCCTCGCTATTCAAGTGTTGGTGGTCATCTCAATAGATC<br>TCCAAGGGCCCATATGGTGGCCAGTGCCGATGAATCCGCCTGTTTAAATG<br>GGGGAGAAAGTTGGGGTTTTAAAACATTTCAAAGTTCCTGAAAAGATCCC<br>ACTAGATCCTGTCACAATTCCCTGAACGCTTTGAAGGCGCGGCCTATTGT<br>CTCCTGGTTATAAATGATATTCCTGGCCAAGTCGATTCCCACA | chr2<br>105689933-105690425 |

Table 1: Natural CRE sequences

| Fragment | Sequence | Origin |
|---|---|---|
| 1 | TTGGTTCTATTCATTTTTCCCTCTAACTGTCCTTGTTTGGGTTGCCTTAG<br>TCCAGGCGAAGCAGAACGCTCAGGCATCGAGTCCTCCAAGCTCACAGCTG<br>CTTGGGCCGCGGCGGCGCCCCCACCACCCCCGGGTTTCTAGAGCCTTGTG<br>ACAGCGCCTCACTCCGAGCTGCTCAGCGGCTTTCTGCAACTCTCTCTGCC | Nkx22<br>chr2<br>147187985-147188184 |
| 2 | TCGCCCCCAGTATGTGACGTGGGTGACAATGGCCCAGGTTGGAGCGAGCC<br>CCACGTCGGCGCGTCCTGGGTGGTCGGACCCGGGCAAACACAAATACAAA<br>CCGATTGCTAAGCTGCGGACAATGAGGGAAATGTAGACAAATGTCCCGCT<br>CCTGTTGGAAGCCTTTGTCCAGGCCCGGTTTTTGCATTTATTTCAGTGGC | Nkx22<br>chr2<br>147188185-147188384 |
| 3 | GAAATAATACATGATTGACGCTCTCTTTCAATGTGTCCTAACTGTTTGGA<br>ATAAATCTAAGGTTGTCCCTAGTTGTCATGGCATTCAACCCTTTTCAATG<br>GACTATCTCTTCATTCATTTCTGAATCCGTCCACAATATTAAGGAAACCC<br>CTTCATGTCGACGTTCCCCATTCCTCTTTCTCCTGCTTTTCTTTTCTCCT | Nkx22<br>chr2<br>147188385-147188584 |
| 4 | TTCTTCCTCCCTTTTCCCCTTGCAAGAATTAGTATCTGGTTTGAAACGTG<br>TTCCTCAGACTCATTACTTTCTCTGTCTTCCCCCCTTCTGTTTTTCCAGC<br>TTCTTTCTCAGCGTTAGAGCCTGTTGCCTCTCCCTCCCTCTTTTCAGGCT<br>GGGGCTCTCTGCCCGGCAGGGTGGGAGGGGGGAAAGTCAGCTTTCCAGAA | Nkx22<br>chr2<br>147188585-147188784 |
| 5 | CTTGGGCCGCGGCGGCGCCCCCACCACCCCCGGGTTTCTAGAGCCTTGTG<br>ACAGCGCCTCACTCCGAGCTGCTCAGCGGCTTTCTGCAACTCTCTCTGCC<br>TCGCCCCCAGTATGTGACGTGGGTGACAATGGCCCAGGTTGGAGCGAGCC<br>CCACGTCGGCGCGTCCTGGGTGGTCGGACCCGGGCAAACACAAATACAAA | Nkx22<br>chr2<br>147188085-147188284 |
| 6 | CCGATTGCTAAGCTGCGGACAATGAGGGAAATGTAGACAAATGTCCCGCT<br>CCTGTTGGAAGCCTTTGTCCAGGCCCGGTTTTTGCATTTATTTCAGTGGC<br>GAAATAATACATGATTGACGCTCTCTTTCAATGTGTCCTAACTGTTTGGA<br>ATAAATCTAAGGTTGTCCCTAGTTGTCATGGCATTCAACCCTTTTCAATG | Nkx22<br>chr2<br>147188285-147188484 |
| 7 | GACTATCTCTTCATTCATTTCTGAATCCGTCCACAATATTAAGGAAACCC<br>CTTCATGTCGACGTTCCCCATTCCTCTTTCTCCTGCTTTTCTTTTCTCCT<br>TTCTTCCTCCCTTTTCCCCTTGCAAGAATTAGTATCTGGTTTGAAACGTG<br>TTCCTCAGACTCATTACTTTCTCTGTCTTCCCCCCTTCTGTTTTTCCAGC | Nkx22<br>chr2<br>147188485-147188684 |
| 8 | TGATTGCCAACTGCCTCCAACCCAACTTTGTAAGCCGAGCCCTCATCCCT<br>ACCCACTCCCGGTGTGTCAGATGGAACACCTAGGTGGCCCACGGGGACCT<br>CTGACCTCTATATCCTCTTCCTTTCTCCCTCCTTTGCTACTTTTCTACTG<br>GATAAAAGGAGAGAGTGAGAGATAATTAACAAAAAACATGGCCCCGGGAC | Olig2<br>chr16<br>91192436-91192635 |
| 9 | AATGAAGCAACTGGCCTTGGCCGGCAAGCAACGATCCTGGTTTTCTAGGT<br>AGAGTTTCTCCCATCAATCTTTCCTTTAACCTCCCTGTTCGTGGAAGCAA<br>TAGAAACACCACCCCCTCCCCTGAGCAAATGCTTTCTTTTGACTGGAAAC<br>AAAAAGGGGGCCCGGCAAAGACGGAGGTGAAATCTGGGTGGTATGGGCAC | Olig2<br>chr16<br>91192636-91192835 |
| 10 | CGCACAATGGCCCCGCTGTTCCTGGCCCTGCTTGTGTTTTACAACAGGGG<br>AGGGGCAGGCGCGAATGGTCCGATGGTGGAGACAATCCCCCTGATTCAGG<br>CTACAAATGCATCTTCTATTCCACACGGAGCTGAGCAGAAAGGATGGGGG<br>TGACAAAGAGCATGGGCGGGGAGAGGGAAAACAAAATGTTTTCAGTTGAA | Olig2<br>chr16<br>91192836-91193035 |

| | | |
|---|---|---|
| 11 | AAAAAAAATCTCTCATATCCTACACATCCTCAGAAGAGCTTCTATGGAGA<br>AGGCCTTCGGAGAGTCCCAGCCCACAACTCAAGGGCTTTGTCTGAACTCT<br>GATTTATTGATGAAGCTTAAGCGGCTCGCTAAGAAAGGCCTGGGGGTGTC<br>TTTGTCTTGAAGATAAAGTACAATAGGCCACAAGGGCCAAGATCTCTCGG | Olig2<br>chr16<br>91193036-91193235 |
| 12 | GATGCTCTCAGGTCCTGCCTCTCTCTTGCCCTCTCCTCCCTGCAAATGCC<br>AGCAGATGCTGAAAAAAAAAAAAAACCCATCGGTGGTGTGGCTGGGAGTGC<br>TGGGGACAAGCTGGGCCACTTGAGGTCTCCTTAAGAGGGTATTATGGCCA<br>GGGAAAATATTTGCGCTCTAAGGATGGCACACTCCATTTGATAATGGCTC | Olig2<br>chr16<br>91193236-91193435 |
| 13 | TCATCTGCCTCAGATAATCGCCTCCCTCCCGGCTGTCAGGGGTGCAGCCA<br>CTGCCAATTCACAGCGCCCTCCGAGAAAGTACCCTTGTCTGTGATGACCA<br>AGATGGGGACATTGTGTTTACCTACTTGAGCAGAGGAGAAGGTGACCGTG<br>AGGGCAGCCTGCATTGTAAATTACAATTAAAACAGAAACAGACAGTTCCT | Olig2<br>chr16<br>91193436-91193635 |
| 14 | GCTCTGCCCTGGGACCCCCACCAATAAATTATGGGTGGACATTAGGGGAG<br>AGCCCAGGAAAGGTTGGGTCCTGGGGAGGATCCCCCCATCCCATAGCCTA<br>CCGACAGGTCTTGGATATAGGGATAGGGCTACTTGGGAGTCAAGGTAGAC<br>TGGCTGGTTGACCACACACACACTGGGATCCTCAGGAGGTTCCCCCCACT | Olig2<br>chr16<br>91193636-91193835 |
| 15 | CTGACCTCTATATCCTCTTCCTTTCTCCCTCCTTTGCTACTTTTCTACTG<br>GATAAAAGGAGAGAGTGAGAGATAATTAACAAAAAACATGGCCCCGGGAC<br>AATGAAGCAACTGGCCTTGGCCGGCAAGCAACGATCCTGGTTTTCTAGGT<br>AGAGTTTCTCCCATCAATCTTTCCTTTAACCTCCCTGTTCGTGGAAGCAA | Olig2<br>chr16<br>91192536-91192735 |
| 16 | TAGAAACACCACCCCCTCCCCTGAGCAAATGCTTTCTTTTGACTGGAAAC<br>AAAAAGGGGGCCCGGCAAAGACGGAGGTGAAATCTGGGTGGTATGGGCAC<br>CGCACAATGGCCCCGCTGTTCCTGGCCCTGCTTGTGTTTTACAACAGGGG<br>AGGGGCAGGCGCGAATGGTCCGATGGTGGAGACAATCCCCCTGATTCAGG | Olig2<br>chr16<br>91192736-91192935 |
| 17 | CTACAAATGCATCTTCTATTCCACACGGAGCTGAGCAGAAAGGATGGGGG<br>TGACAAAGAGCATGGGCGGGGAGAGGGAAAACAAAATGTTTTCAGTTGAA<br>AAAAAAAATCTCTCATATCCTACACATCCTCAGAAGAGCTTCTATGGAGA<br>AGGCCTTCGGAGAGTCCCAGCCCACAACTCAAGGGCTTTGTCTGAACTCT | Olig2<br>chr16<br>91192936-91193135 |
| 18 | GATTTATTGATGAAGCTTAAGCGGCTCGCTAAGAAAGGCCTGGGGGTGTC<br>TTTGTCTTGAAGATAAAGTACAATAGGCCACAAGGGCCAAGATCTCTCGG<br>GATGCTCTCAGGTCCTGCCTCTCTCTTGCCCTCTCCTCCCTGCAAATGCC<br>AGCAGATGCTGAAAAAAAAAAAAAACCCATCGGTGGTGTGGCTGGGAGTGC | Olig2<br>chr16<br>91193136-91193335 |
| 19 | TGGGGACAAGCTGGGCCACTTGAGGTCTCCTTAAGAGGGTATTATGGCCA<br>GGGAAAATATTTGCGCTCTAAGGATGGCACACTCCATTTGATAATGGCTC<br>TCATCTGCCTCAGATAATCGCCTCCCTCCCGGCTGTCAGGGGTGCAGCCA<br>CTGCCAATTCACAGCGCCCTCCGAGAAAGTACCCTTGTCTGTGATGACCA | Olig2<br>chr16<br>91193336-91193535 |
| 20 | AGATGGGGACATTGTGTTTACCTACTTGAGCAGAGGAGAAGGTGACCGTG<br>AGGGCAGCCTGCATTGTAAATTACAATTAAAACAGAAACAGACAGTTCCT<br>GCTCTGCCCTGGGACCCCCACCAATAAATTATGGGTGGACATTAGGGGAG<br>AGCCCAGGAAAGGTTGGGTCCTGGGGAGGATCCCCCCATCCCATAGCCTA | Olig2<br>chr16<br>91193536-91193735 |
| 21 | CTAAATAGCACCGCGGCGCCCGCTCTCCGGACAGTGATTAATGATAGCAG<br>TGCAGAGGGGTTAACACACTTCGCTGAAAAGTCTGTTGACTGAGCTTCTT<br>GTAACACAATGTGGCCCGCTGCACGCCTCGAGAGAATCCTTTTGTTGTCC<br>GTGCTTATTGTGGCCTCAAAATTCTGCCCACGAAAGTTTGCCAACGCTCC | Olig2<br>chr16<br>105689933-105690132 |
| 22 | TGCCCCAGGAGTTTAATAGTTTCCCTTACTCGCGGGGCATTGTGTGGTGC<br>TGAAAAGCAGCCCCTCGCTATTCAAGTGTTGGTGGTCATCTCAATAGATC<br>TCCAAGGGCCCATATGGTGGCCAGTGCCGATGAATCCGCCTGTTTAAATG<br>GGGGAGAAAGTTGGGGTTTTAAAACATTTCAAAGTTCCTGAAAAGATCCC | Pax6<br>chr2<br>105690133-105690332 |
| 23 | ACTAGATCCTGTCACAATTCCCTGAACGCTTTGAAGGCGCGGCCTATTGT<br>CTCCTGGTTATAAATGATATTCCTGGCCAAGTCGATTCCCACAGAGGCCT<br>GGCGCCCCCTCCCCCCAAACTAGCCAAGTTTCTAGGATCCGGAGAGAACA<br>GTTCTTGTTGGTATCCCGTGGGGGCCACAGGCCTCAGGACCCAAGACAGC | Pax6<br>chr2<br>105690333-105690532 |

| | | |
|---|---|---|
| 24 | GTAACACAATGTGGCCCGCTGCACGCCTCGAGAGAATCCTTTTGTTGTCC<br>GTGCTTATTGTGGCCTCAAAATTCTGCCCACGAAAGTTTGCCAACGCTCC<br>TGCCCCAGGAGTTTAATAGTTTCCCTTACTCGCGGGGCATTGTGTGGTGC<br>TGAAAAGCAGCCCCTCGCTATTCAAGTGTTGGTGGTCATCTCAATAGATC | Pax6<br>chr2<br>105690033-105690232 |
| 25 | TCCAAGGGCCCATATGGTGGCCAGTGCCGATGAATCCGCCTGTTTAAATG<br>GGGGAGAAAGTTGGGGTTTTAAAACATTTCAAAGTTCCTGAAAAGATCCC<br>ACTAGATCCTGTCACAATTCCCTGAACGCTTTGAAGGCGCGGCCTATTGT<br>CTCCTGGTTATAAATGATATTCCTGGCCAAGTCGATTCCCACAGAGGCCT | Pax6<br>chr2<br>105690233-105690432 |

Table 2: CRE Fragments

| Fragment | 'Odd' Barcodes (5'-3-) | 'Even' Barcodes (5'-3-) |
|---|---|---|
| 1 | CACTGGAT | GACAACAC |
| 2 | CTCCTCTA | CCTGATCA |
| 3 | CTAAGGTG | CACTTGGC |
| 4 | TCGCGATA | GCCATTCT |
| 5 | GGCCTATT | GTACGAGT |
| 6 | TGTGTGCA | GATGAACG |
| 7 | GGATATGC | CGAGTCTA |
| 8 | ACTCTCTC | GTCAGTGT |
| 9 | TATCCAGC | CTACCATG |
| 10 | GAGTATCC | GAGTCTAC |
| 11 | CCTTCAAG | ATGTGCTG |
| 12 | TACGCCGT | GTTCGTAC |
| 13 | AGAACACG | AGGCTGTT |
| 14 | GCTAGTAG | CCGACGTA |
| 15 | AGCAACTC | AGCTCACT |
| 16 | CCAACTGA | AGAGACAC |
| 17 | ATCGACCA | ACGAACAC |
| 18 | AGAGTGGT | AGAGCGGA |
| 19 | TGGATGCT | AACCGAAG |
| 20 | GAACACAC | GCTTGTCG |
| 21 | TCGTGAGT | CTTCCGTG |
| 22 | GTGTAGCG | ACAGACCA |
| 23 | CATGCTCA | GAGTGCGT |
| 24 | ATGGCGAG | CCTCTAGT |
| 25 | GTGCGTAC | CGAGTGGC |

Table 3: Allocated Barcodes by Fragment
